# A tuned topographic map hierarchy formats visual working memory load

**DOI:** 10.64898/2026.11.14.766649

**Authors:** Ben M. Harvey, Martijn van Ackooij, Joeri F.L. van Helden, Evi Hendrikx, Nathan van der Stoep, Surya Gayet, Jacob M. Paul

## Abstract

Visual working memory tasks rely on neural systems that scale activity with task load. Traditional models assume higher demands are met by monotonically increasing activity within a single process, implying high informational redundancy across responses. However, theories of cognitive control suggest processing strategies change with task demands, flexibly linking sensory inputs to behavior. Using 7T fMRI and neural encoding models, we hypothesized and confirmed that association cortices format task load-dependent responses using sensory-like neural tuning. We identified populations selectively tuned to specific loads, forming 11 topographic maps per hemisphere hierarchically organized from posterior sensory to anterior executive regions across a fronto-parieto-occipital network. Tuning persisted across performance and trial-order controls, was absent without memory task demands, and predicted behavioral accuracy after load changes. These findings demonstrate that human association cortices distribute task demands using continuously varying processing pathways, adapting sensory-like tuning to decorrelate responses and reduce information redundency during cognitive control.

## Introduction

Visual short-term memory (VSTM) allows us to briefly retain visual information to guide goal-directed behavior, such as remembering the road ahead while checking a rear-view mirror. Neuroimaging studies have predominantly focused on how memory content (what is remembered) is represented in the brain^1–6^. Far less is known about how brain activity changes with task demands like the amount of visual information remembered (VSTM load), a common behavioral metric^7^.

Increasing task demands require increasing cognitive resource allocation^8,9^. Standard neuroimaging accounts assume that neural responses in supporting brain areas scale monotonically with VSTM load^10,11^, implying that the brain simply ramps up activity within a single, widespread processing strategy. However, theories of cognitive control propose more complex responses representing task goals and strategies^12^. Such responses dynamically reconfigure processes throughout the brain linking sensory inputs to adaptive behaviors. Conceptually, as task demands increase, our cognitive strategy and the required neural processing may gradually adapt and shift, rather than simply allocating more effort to a single strategy. This framework predicts diverse, task demand-dependent neural responses distributed across neural populations, rather than uniform monotonic scaling throughout the brain.

In sensory systems, efficient processing depends on three core principles: neural tuning, where distinct populations respond preferentialy to specific stimulus states^13,14^; topographic mapping, where preferences progress continuously across the cortical surface^15–20^; and hierarchical processing, where response properties systematically transform across successive brain areas^19–22^. Information-theoretically, selective tuning decorrelates population signals, minimizing redundancy and maximizing coding efficiency compared to monotonic scaling^23–26^. Encoding cognitive task demands using these mechanisms could optimize how the brain reconfigures processing to flexibly link sensory inputs to behavior. We therefore hypothesized that neural responses to VSTM load follow a tuned organizational framework, with distinct neural populations within and between brain areas selectively responding to different VSTM loads.

To test whether neural responses to VSTM load follow a tuned organizational framework, we designed a paradigm that parametrically manipulated load (Fig. 1A,B; Video S1). Participants viewed a central six-item Memory Display of colored bars and remembered the orientations of 1 to 6 unique items (i.e. loads) over a brief delay before judging whether a colored probe bar matched the orientation remembered. Holding spatial layout, item count, and extent (<1.6° eccentricity) identical across loads minimized physical visual stimulation changes, ensuring neural response changes reflected load.

**Figure 1.**
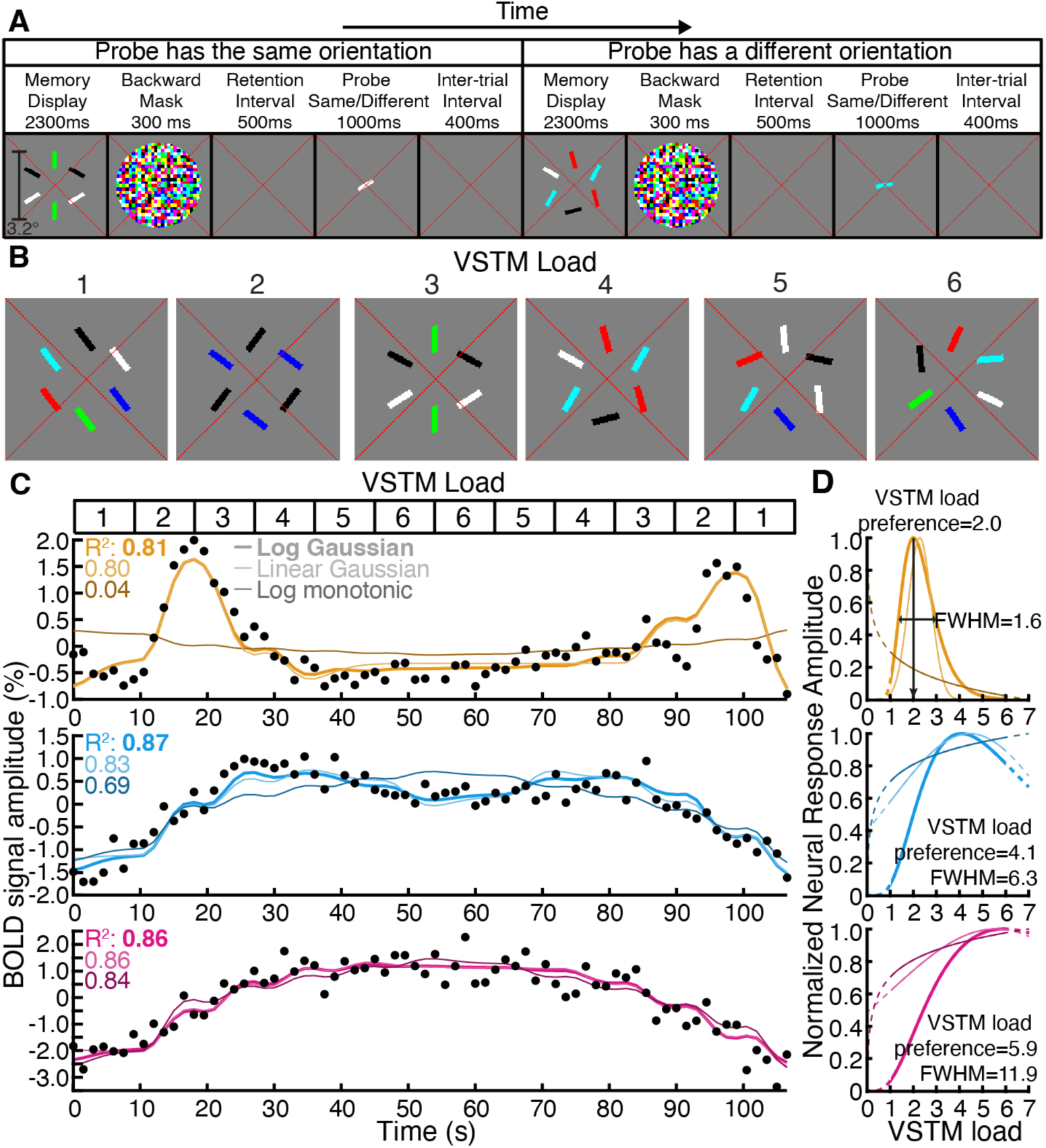
Experimental paradigm, stimuli, and load-tuned fMRI response time courses. **(A)** Trial sequence illustrated for two consecutive trials (loads 3 and 4). **(B)** Example Memory Displays for each VSTM load. At load 1, all bars shared a single orientation but were uniquely colored to maintain local color contrast. For higher loads, bars sharing the same orientation were identically colored and non-adjacent. **(C)** Observed fMRI BOLD time courses (black dots) from three nearby voxels in one participant’s intraparietal sulcus during a sequence of changing VSTM loads (top inset). **(D)** Best-fitting neural response functions for each voxel in (C). Differences in these estimated parameters of the winning log-Gaussian model account for the distinct time courses in (C). Dashed lines show functions extended beyond the tested load range.

## Results

### Behavioral performance and task controls

To dissociate effects of VSTM load from behavioral performance, participants underwent pre-scan staircased training to establish stable load-specific discrimination thresholds (Fig. S1A). During ultra-high field (7T) fMRI, we tested two conditions: a *constant performance* condition, where probe orientation differences were calibrated to equate accuracy across loads, and a *constant orientation difference* condition, where probe differences were fixed across loads (causing accuracy to drop as load increased). This isolated neural responses to load from responses following behavioral accuracy.

As expected, during fMRI scanning in the *constant performance* condition, behavioral accuracy did not change with VSTM load (*F*(1,34)=0.37, *p*=0.55; linear mixed-effects model with participant as a categorical random factor and VSTM load as a continuous fixed predictor; Fig. S1B). Conversely, in the *constant orientation difference* condition performance decreased significantly with increasing VSTM load (*F*(1,34)=100.72, *p*<10^-10^; Fig. S1C). Together, these behavioral results confirm that our thresholding procedure successfully equated objective behavioral accuracy across loads in the constant performance condition, demonstrating that VSTM load-tuning profiles can be isolated independently of changes in task accuracy.

### Tuned responses to VSTM load

In both task conditions (Fig. S2), variations in VSTM load elicited strong responses with remarkably different profiles across nearby recording sites (voxels), peaking after different loads (Fig. 1C, S3). To formally characterize these response profiles, we compared the observed time courses to predictions from five candidate encoding models linking VSTM load to neural response amplitude (Fig. 1C,D; Methods). These included three monotonic models (response proportional to VSTM load, its logarithm, or VSTM load raised to an estimated power parameter) and two tuned models (response proportional to a linear-Gaussian or log-Gaussian function of VSTM load). For tuned models, the preferred VSTM load (μ, Gaussian mean) and tuning width (σ, standard deviation i.e. range of VSTM loads producing a response) were fit as free parameters.

In cross-validated data, the log-Gaussian model best described neural responses across an extensive fronto-parietal network (Fig. 1C, S4). This model significantly outperformed the next-best candidate model (linear-Guassian, *t*=11.3, *p*<10^-10^, n=264, paired *t*-test across map examples; Fig. S4). The distinct time courses observed across nearby voxels (Fig. 1C, S3) were closely predicted by systematic variations in the load preference (μ) and tuning width (σ) parameters of this log-Gaussian function (Fig. 1D, S3).

### Model Validation via Simulations

To ensure our pipeline could reliably distinguish between tuned and monotonic profiles, we validated our model-fitting procedure using synthetic BOLD responses generated from ground-truth monotonic and tuned functions across a wide range of parameters and noise levels (Fig. S5). Ground-truth tuned responses were accurately identified and distinguished from monotonic responses whenever cross-validated variance explained (*R*^2^_CV_) exceeded 0.2 and estimated VSTM load preferences fell between 1.5 and 5 (and with >50% accuracy outside this range). While ground-truth monotonic responses were incorrectly classified as tuned in ∼40% of cases with *R*^2^_CV_ > 0.2, these false positives produced edge-of-range preference estimates (<1.5 or >5). Thus, monotonic and tuned profiles are cleanly distinguishable within the core tested range (1.5–5 items) when fit quality is robust (*R*^2^ > 0.2). Our validation also confirmed that the pipeline reliably distinguished linear-Gaussian from log-Gaussian functions under similar conditions. To ensure data quality and avoid classification artifacts, we excluded voxels with variance explained below 0.2 or estimated load preferences outside the tested range from model comparisons and subsequent analyses.

### VSTM load preferences organize into widespread topographic maps

Projecting estimated VSTM load preferences onto individual cortical surfaces revealed organized topographic progressions, where preferred loads changed systematically across the local cortical surface (Fig. 2, S7). Following visual field mapping conventions^20^, we identified 11 distinct topographic maps per hemisphere characterized by progressive increases from low to high load preferences across participants (Fig. S8, Table S1 for template locations).

**Figure 2.**
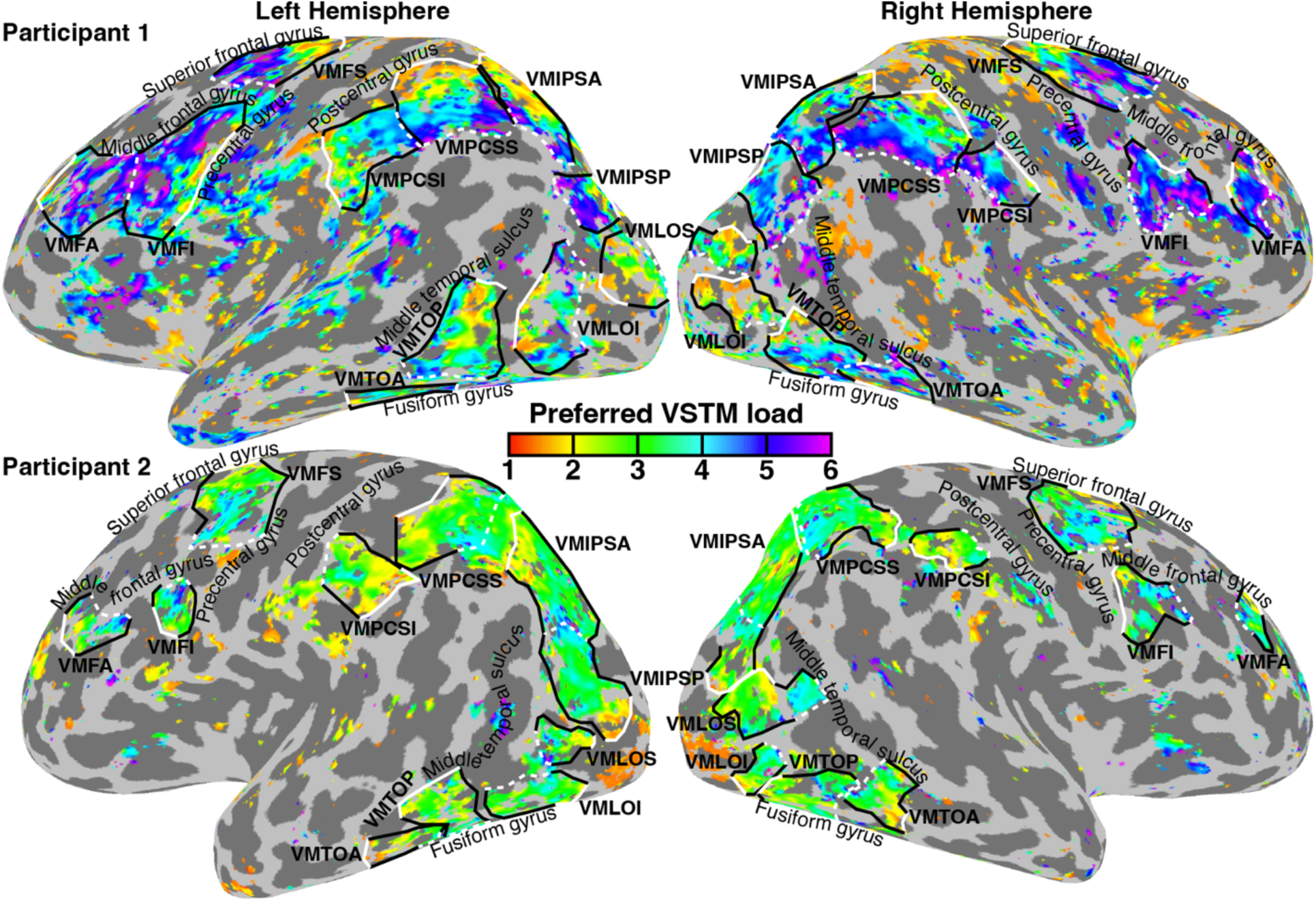
Topographic maps of VSTM load preferences across the cortical surface. Solid white lines indicate low-load preference borders; dashed white lines indicate high-load preference borders. Black text labels major anatomical gyri and sulci used to define map orientations. Voxels with variance explained *R*^2^<0.2 or estimated load preferences outside the tested range are uncolored. Two participants shown; see Fig. S7 for all participants.

We named these maps using their anatomical locations preceded by ‘VM’ for ‘visual memory’, following naming conventions for visual field maps and maps of physical quantity preferences^19,20,22^.

- Occipital & Temporal Cortices: VMLOI (lateral occipital inferior) and VMLOS (lateral occipital superior) in lateral occipital cortex; VMTOP (temporal-occipital posterior) and VMTOA (temporal-occipital anterior) on the inferior temporal-occipital boundary.
- Parietal Cortex: VMIPSP (intraparietal sulcus posterior) and VMIPSA (intraparietal sulcus anterior) in the intraparietal sulcus extending into superior parietal lobule; VMPCSS (post-central sulcus superior) and VMPCSI (post-central sulcus inferior) in postcentral sulcus, extending into postcentral and supramarginal gyri.
- Frontal Cortex: VMFS (frontal superior) and VMFI (frontal inferior) in precentral sulcus; VMFA (frontal anterior) on the middle frontal gyrus.

These tuned load maps were widely distributed across cortical regions implicated in visual sensory processing (occipital and parietal areas), attentional control (intraparietal, postcentral and precentral sulci), and executive function (middle frontal gyrus).

Within every map, voxels spanned a range of load preferences (Fig. 2, 3, S3, S7, S11). Split-half reliability analyses confirmed that these load preferences were highly reproducible across independent scan runs (odd vs. even runs correlated significantly in 108 of 132 maps [81%], Spearman’s rank, FDR-corrected *p*<0.05; Table S2). Furthermore, voxel load preferences correlated strongly between the constant performance and constant orientation difference conditions (correlated significantly in 103 of 132 maps [78%], Table S3, Fig. S9–S11). This high cross-condition consistency demonstrates that these topographic maps track internal VSTM load rather than objective behavioral accuracy. Data from both conditions were therefore averaged for subsequent analyses.

**Figure 3.**
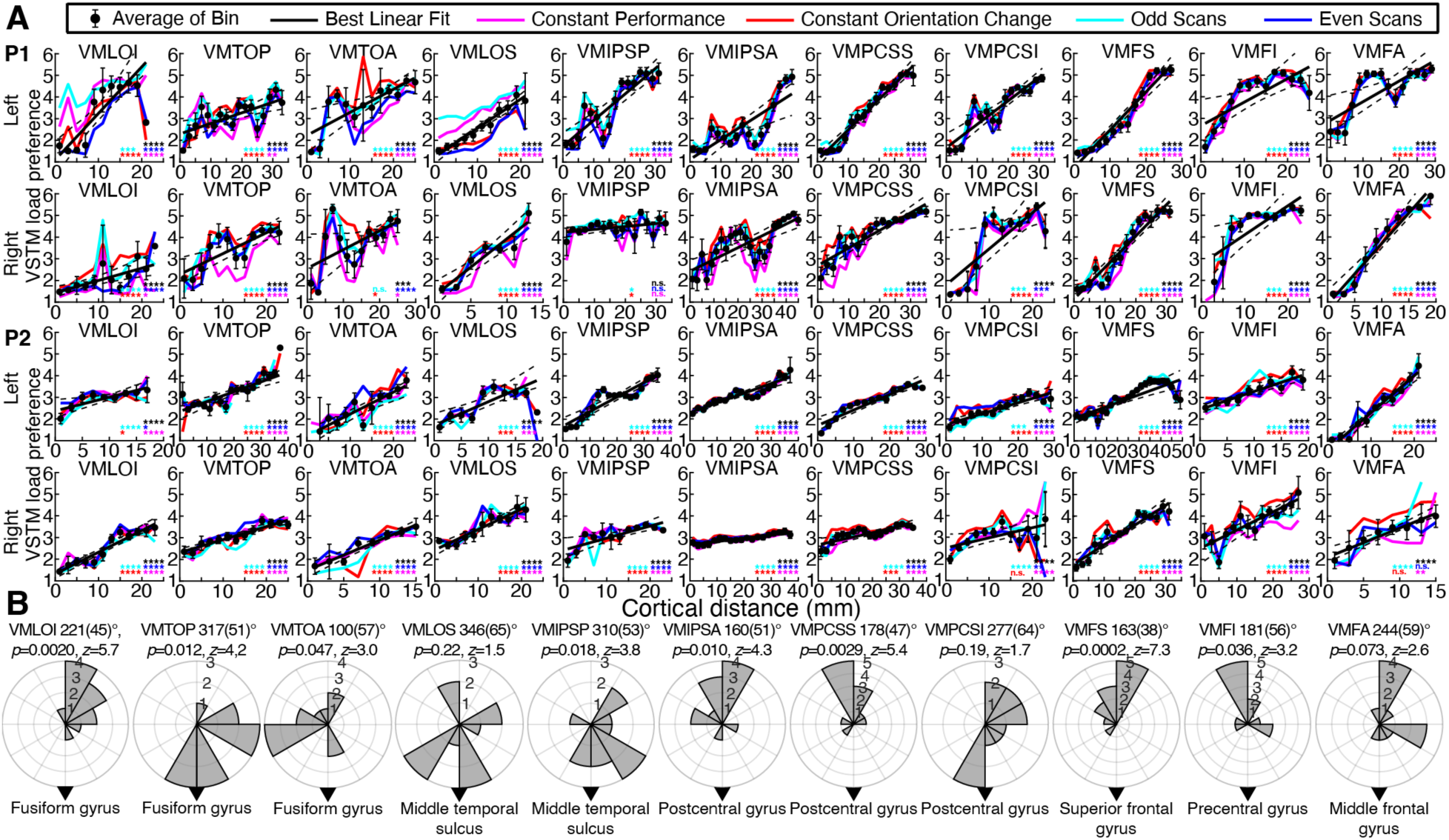
Topographic progressions and anatomical alignment of VSTM load preferences. **(A)** VSTM load preferences plotted per 2mm of cortical surface distance (from low- to high-preference map borders) for the hemispheres in Fig. 2 (all hemispheres in Fig. S11). Data are shown for the grand-averaged data (black) alongside split-half and task condition subsets (colored lines). Significance levels for unbinned correlations are indicated per condition (\**p*<0.05, \*\**p*<0.01, \*\*\**p*<0.001, \*\*\*\**p*<0.0001). **(B)** Orientation of topographic progressions relative to gross macro-anatomy. Polar plots show the distribution of mean preference progression directions across individual hemispheres relative to the shortest path to a nearby major sulcus or gyrus (black arrows). Radial axes indicate hemisphere counts per angular bin.

### Quantifying consistency of topographic load progressions

To quantify the spatial progression of load tuning along the cortical surface, we measured the relationship between each voxel’s preferred VSTM load and its distance from the low- and high preference map borders (Fig. 3A, S10). VSTM load preferences were significantly correlated with cortical distance in 121 out of 132 maps (92%) in the grand averaged dataset (Fig. 3A, S11, Table S4; Spearman’s rank, FDR-corrected *p*<0.05). These spatial progressions were highly repeatable across independent split-half splits and task conditions, showing significant correlations in: 84% of maps (111/132) in the constant performance condition (Table S5); 85% of maps (112/132) in the constant orientation difference condition (Table S6); 84% of maps across odd/even scan splits (109 and 112 out of 132; Tables S7, S8). Crucially, when map boundaries were cross-validated (maps defined on odd scans and tested on even scans, and vice versa), load preferences remained significantly correlated with cortical distance in 75–78% of maps (99 and 103 out of 132; Fig. S12, Tables S9, S10). These metrics confirm a robust, highly repeatable topographic progression of VSTM load preferences across the cortical surface within each map.

To test whether the spatial orientation of these progressions was consistent across participants, we computed the local gradient direction of preference increase relative to paths to nearby major anatomical landmarks (gyral crowns or sulcal fundi). Rayleigh’s test for circular uniformity revealed significant directional alignment in 8 of the 11 maps (73%, FDR-corrected) (Fig. 3B). This demonstrates that topographic progressions of VSTM load preferences follow consistent directional axes relative to gross anatomy across individuals.

### Tuning width increases systematically with VSTM load preference

Systematic relationships between response preferences and tuning widths are a hallmark of tuned sensory systems^13,18,21,27^. Similarly, for VSTM load-tuned responses, linearised tuning widths (full width at half maximum, FWHM) increased systematically with VSTM load preference across all 11 maps (all *p* < 0.001, Spearman correlation, FDR-corrected; Fig. S13). Thus, neural populations preferring low VSTM loads responded to a narrower range of load conditions, maintaining high response selectivity between low loads. Conversely, populations preferring high VSTM loads exhibited broader tuning profiles that overlap with more loads.

### Simulation of edge-of-range biases

Our validation simulations (Fig. S14) demonstrate that edge-of-range constraints can introduce systematic estimation biases when true VSTM load preferences exceed 4 items. Under these extreme conditions, empirical models tend to underestimate both the true load preference and true tuning width. Fully eliminating these edge ambiguities would require testing load levels far exceeding participants’ behavioral memory limits (>6 items). Crucially, because these simulation biases operate in the direction of underestimation, they do not generate false-positive progressions; rather, they indicate that the true progressions of load preferences and tuning widths across our topographic maps are likely more pronounced than reported.

### Response properties transform hierarchically across VSTM load maps

Beyond local intra-map progressions, neural response properties transformed systematically across the macro-level hierarchy of maps. Mean VSTM load preferences differed significantly across maps (*F*(10,116)=3.34, *p*=0.0007; linear mixed-effects model with map as a fixed factor and participant as a random factor), increasing progressively from posterior sensory to anterior executive areas (Fig. 4A).

**Fig. 4.**
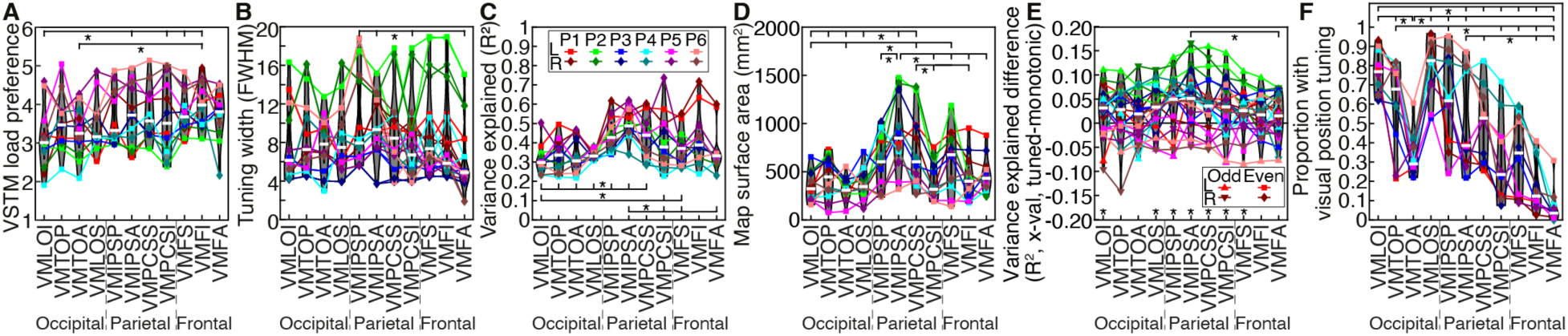
Hierarchical transformations of response properties across VSTM load maps. **(A)** Mean VSTM load preference per map increased from posterior (left) to anterior (right) locations. Mean load-tuning width **(B)**, model variance explained **(C)**, responsive surface area **(D)**, and fit advantage of tuned over monotonic models **(E)** were highest in superior parietal maps. **(F)** Proportion of VSTM load-tuned voxels also showing visual field position tuning (spatial pRFs) declined progressively from posterior visual toward anterior and inferior executive areas. In all panels, points indicate individual participant means per hemisphere; white bars represent across-hemisphere means. Horizontal brackets indicate statistically significant differences between maps (*p*<0.05; all maps connected to brackets on the left of the star differ significantly from those connected on the right). Asterisks below map labels in (E) highlight maps where Δ*R*^2^ was significantly greater than zero (*p*<0.05, FDR corrected).

Furthermore, mean tuning widths (*F*(10,116)=3.22, *p*=0.001; Fig. 4B), model variance explained (*F*(10,116)=9.71, *p*<10^-10^; Fig. 4C) and overall map surface area (*F*(10,116)=9.81, *p*<10^-10^; Fig. 4D) varied significantly across maps, peaking in superior parietal maps (VMIPSP, VMIPSA, VMPCSS). Thus, load-tuned neural responses undergo systematic hierarchical transformations: posterior maps predominantly respond under lower VSTM loads, whereas anterior and superior maps respond under higher loads, with superior parietal cortices exhibiting the clearest tuning, largest cortical coverage and slightly larger tuning widths.

We also evaluated the cross-validated variance explained difference between the best-fitting tuned and monotonic models (Δ*R*^2=^*R*^2^_tuned_-*R*^2^_monotonic_, isolating fit advantages unique to tuned functions), which varied significantly across maps (*F*(10,248)=2.18, *p*=0.019; Fig. 4E). While tuned models fit better on average across all maps (Fig. S4), post-hoc testing revealed that Δ*R*^2^ was significantly above zero in occipital-parietal and superior frontal maps, but did not reach statistical significance in temporal-occipital maps (VMTOP, VMTOA) or extreme frontal maps (VMFI, VMFA). This indicates that while load tuning is robust across the broader fronto-parietal network, evidence for selective tuning over monotonic scaling is strongest in parietal and superior frontal cortices.

### VSTM load tuning reflects cognitive demand, not stimulus complexity

Memory Displays in the main experiment necessarily varied in visual complexity alongside VSTM load (Fig. 1B). We isolated sensory response effects by presenting identical visual displays with reversed task demands (Video S2): Participants memorized a single-item target and determined whether that item appeared in a subsequent six-item display, maintaining a constant VSTM load of 1 item despite changing display complexity.

Modeling BOLD responses as a function of display complexity yielded preference estimates that correlated with the main experiment in only 16.7% of maps (11/66 maps in three participants; Spearman correlation, *p*<0.05 FDR-corrected). This absence of consistent tuning under constant memory load confirms that tuned neural responses require active variation in internal VSTM load rather than visual display features.

### Counterbalancing Trial Order Rules Out Sequence Artifacts and Load Expectations

The main experiment varied VSTM load progressively across trials to maximize encoding model power given fMRI’s slow timescale^28^. To rule out load anticipation, trial history, or order-related hemodynamic artifacts, a second control experiment (in five participants) presented an unpredictable, trial-counterbalanced load sequence (Video S3). The log-Gaussian model remained the winning model across cross-validated data (Fig. S4E–F, S15). Load preference estimates correlated significantly with those from the progressive design in 65.5% of maps (72/110) rising to 75.8% (72/95) in maps with robust responses in both experiments (Table S13). Furthermore, log-Gaussian models trained on the unpredictable control generalized to explain time courses in the main experiment (and vice versa) better than alternative candidate models (Fig. S4G–J). These results confirm that population tuning reflects current internal VSTM load.

### Trial-to-trial switching costs confirm distinct load-dependent cognitive strategies

To establish whether increasingly different VSTM loads recruit increasingly different cognitive processing strategies rather than monotonically varying resource allocation levels, we evaluated task accuracy across trial transitions in the counterbalanced unpredictable trial order control experiment. If different load levels demand distinct processing strategies, larger changes in load between consecutive trials (|Δload|) should incur greater strategy-switching costs. In line with this hypothesis, participant accuracy declined significantly as the magnitude of trial-to-trial VSTM load change increased (*F*(1, 24)=5.81, *p*=0.024).

Because load difficulty was threshold-matched and transition frequencies were fully counterbalanced, this performance cost is consistent with a cognitive overhead of reconfiguring processing strategies that elicit distinct load-tuned responses. Monotonically varying cognitive resource allocation predicts increasing and decreasing load changes would affect accuracy differently. However, this direction of change had no significant effect (*F*(1,47)=0.26, *p*=0.61) or interaction with load change (*F*(1,47)=0.96, *p*=0.33).

### VSTM load maps share cortical territory with independent visual field maps and numerosity field maps

Because many VSTM load maps partially overlap with visual field maps, we evaluated whether load tuning relates to retinotopic tuning. In five participants with visual field mapping data, the proportion of load-tuned voxels displaying spatial population receptive field tuning differed significantly across maps (*F*(10,95) = 25.6, *p* < 10^-10^), declining from a high degree of overlap in posterior and superior visual areas to negligible overlap in anterior executive regions (Fig. 4F). Within these overlapping regions, VSTM load preferences correlated with visual field eccentricity (Fig. S16) in only 26 out of 110 maps (24%) and polar angle (Fig S17) in only 11 out of 110 maps (10%). Goodness of fit (*R*^2^) correlated significantly between spatial and load-tuning models positively in only 5 maps (5%) and negatively in 4 maps (4%).

Finally, we previously identified visual numerosity-tuned responses near some VSTM load maps^22^. Although our Memory Displays held item count constant (always 6), we tested whether VSTM load tuning was related to numerosity tuning. In three participants with independent numerosity mapping data (Fig. S18), VSTM load preferences and visual numerosity preferences correlated significantly in only 7 out of 66 maps (11%). Unlike consistent relationships between numerosity and other sensory quantities^14,29–31^, VSTM load tuning is at most weakly related to numerosity tuning. Therefore, VSTM load preferences are independent of both visual numerosity tuning and retinotopic spatial representations.

## Discussion

### Computational mechanisms and benefits of tuned task demand responses

Our findings challenge classic univariate accounts that assume neural activity scales monotonically with task load to allocate effort within a widespread process^10,11^. Instead, we demonstrate that responses to visual short-term memory (VSTM) load are formatted using principles established in sensory systems: selective neural tuning, organized topographic mapping, and hierarchical transformations across fronto-parietal networks. Neural populations preferring loads >6 exhibit profiles within our tested range that resemble monotonic scaling, concentrated primarily in frontal cortex. These high-preference responses may represent drive signals that are transformed to generate tuned responses across fronto-parieto-occipital cortices, a mechanism well-established in visual and quantity processing^32–35^.

This distributed population encoding provides an information-theoretically efficient mechanism for reconfiguring neural responses between task contexts^23–26^. Whereas monotonic channels carry redundant information across similarly-responding neural populations, selective tuning decorrelates population activity, minimizing redundant overlap. This may optimize the brain’s capacity to process sensory inputs flexibly across cognitive demands,.

This architecture directly supports theories of flexible cognitive control^12^, which propose that the brain manages task demands by reconfiguring functional pathways to flexibly link sensory inputs to behavioral outputs. We demonstrate that association cortices establish continuously varying processes across topographically organized pathways rather than ramping up a single global process. Aligning with our behavioral switching costs, tuned subpopulations establish processes that become increasingly distinct as task differences grow. Furthermore, the broader tuning widths observed at higher load preferences reveal a computational trade-off: sharp tuning curves finely separate processing pathways between low loads, whereas broader tuning at high loads pools resources to maintain network robustness without requiring an unfeasible proliferation of narrow channels.

### Topographic maps and macro-level processing hierarchies

Once tuned responses emerge, topographic organization groups similarly tuned populations together, minimizing local wiring costs and facilitating efficient lateral interactions^36–38^. Similar to retinotopic organization, we observed a continuous series of abutting load maps across the cortex. Defining topographic borders in novel cortical territory relies on manual delineation. Although standard across mapping disciplines, manual delineation retains an inherent degree of subjectivity. Given the large number of newly identified maps, future studies may propose refined boundary definitions, echoing debates that have persisted for decades in visual field mapping.

Despite their consistent internal topographic progressions, we observed notable individual variation in map positions and orientations. This spatial variability is higher than that seen in early occipital retinotopic maps, which are tightly constrained by major anatomical landmarks and subcortical projections (such as the optic radiation terminating in the calcarine sulcus, alongside the interlocking architecture of V1–V3^22^). Instead, the macro-anatomical variability observed here resembles that of association cortices, including fronto-parietal visual field maps^39^, numerosity maps^22^, and timing maps^19^.

The observed hierarchy of VSTM load preferences closely mirrors established processing schemes across cortical systems, including visual field^20^, numerosity^22^, and visual timing maps^19^. In sensory systems, processing hierarchies reflect the progressive integration and refinement of stimulus representations. For VSTM load, however, this hierarchy reflects a progressive expansion of cognitive recruitment: increasingly anterior areas are recruited at higher VSTM loads. Posterior and inferior maps in sensory-adjacent regions may suffice to accommodate low task demands and simpler task strategies. In contrast, higher VSTM loads recruit more extensive parietal and frontal networks^40^ to configure higher-order processes and maintain task performance.

### VSTM sub-phases, effort, and task complexity

Although fMRI lacks the temporal resolution to isolate sub-trial events, several design choices suggest our load-tuned responses are primarily driven by VSTM encoding. The encoding phase was subjectively effortful and presented for 2300ms to allow accurate encoding even at high loads, generating a larger time-integrated BOLD signal. Conversely, maintenance (800ms) and single-item retrieval were brief and relatively effortless.

Higher VSTM loads inherently demand more encoding effort and involve greater task complexity, as displays with more unique orientations contain less redundant information and are less amenable to chunking^41^. Crucially, these load-tuned maps reflect task complexity and effort rather than physical stimulus complexity, as tuning vanished when identical displays were viewed without memory demands. These findings align with reports that fronto-parietal activity scales with the complexity of remembered sequences^42^ and that subjective encoding effort remains high across load conditions even when probe discrimination thresholds are eased.

### Executive control vs. VSTM content representation

The fronto-parietal network identified here heavily overlaps with core executive control networks^43–45^. Increased working memory load has long been linked to elevated frontal activity and fronto-parietal coupling^46,47^, consistent with the posterior-to-anterior hierarchy we observed across our maps.

Conversely, we found no load-tuned responses in early visual cortex. This does not contradict studies demonstrating that visual cortex maintains specific VSTM content^3,5,40^. Rather, it highlights a functional dissociation: early visual areas represent sensory content, whereas fronto-parietal association cortices respond to task demands and executive processing strategies^48^. Reusing sensory-like encoding schemes (tuning, topography, and hierarchies) for abstract cognitive demands likely optimizes bidirectional interactions between executive control and sensory content during working memory.

A critical question for future research is whether VSTM load tuning and its topographic organization generalize across different visual features, categories, or sensory modalities. Because sensory cortices recruit distinct neural populations for different features (e.g., motion, faces, or color), load-tuning profiles in association cortices may be similarly constrained by the feature-selective networks they recruit. Determining whether feature domains rely on distinct processing strategies and load responses or a single domain-general framework remains an open question. The paradigm and modeling pipeline established here provide a clear foundation to test how the brain applies its task processing architecture across diverse short-term memory representations.

In summary, association cortices adapt sensory-like tuning and topographic mapping principles to distribute cognitive task demands across selectively tuned sub-populations. This decorrelates neural signals to establish an information-theoretically efficient framework for flexible executive processing pathways that progressively transition with task demands. High-resolution neuroimaging offers powerful approaches to investigate these cognitive processing pathways in the human brain.

## Methods

### Experimental model and participant details

Six right-handed participants (aged 24–38 years, two female) with normal or corrected-to-normal vision participated in the study. In line with high-resolution 7T fMRI encoding and pRF mapping conventions^18,19,22,49,50^, we adopted a deep sampling approach, acquiring extensive data per individual to ensure robust within-subject replication across independent hemisphere measurements and cortical maps^51,52^. All participants provided written informed consent. Experimental procedures were approved by the Ethics Committee of the University Medical Center Utrecht (Protocol No. 17/414).

### Visual stimuli and VSTM task paradigm

Visual stimuli were generated using MATLAB (MathWorks) with Psychtoolbox-3 and displayed on a BOLDscreen 32 LCD monitor (1920 ×1080 pixels, 69.8 × 39.3 cm; Cambridge Research Systems) positioned at the head of the scanner bore. Participants viewed the display through a mirror at a total viewing distance of 220cm. A red diagonal cross spanning the screen was present throughout all runs to facilitate central fixation.

Each trial lasted 4.5 s (Fig. 1A, Video S1) and comprised:

1) Memory Display (2.3 s): An array of six colored bars each 0.8° × 0.2°, centered 1.2° from fixation and extending up to 1.6° from fixation. To vary VSTM load, the array contained between 1 and 6 unique bar orientations (Fig. 1B). Bar colors were easily distinguished (black, white, red, green, blue, cyan).
2) Backward Mask (0.3 s): A dynamic mask covering the array area to prevent visual afterimages.
3) Retention Delay (0.5 s): A blank gray screen.
4) Probe Display (1.0 s): A single colored probe bar presented with one of the array colors, presented at fixation. Participants indicated via button press whether the probe bar’s orientation matched the orientation of the corresponding bar in the Memory Display.
5) Inter-Trial Interval (0.4 s): A blank gray screen.

The response window lasted 3.7 s, from the start of the probe display until the end of the following trial’s Memory Display. Trials with no response (0.64%) were excluded from behavioral analyses.

### Main fMRI experiment paradigm and staircase training

In the main experiment, VSTM load varied progressively within each scanning run in an ascending and descending block design (Fig. 1C, top inset; Video S1) optimized for fMRI encoding models^28^. Each load level (1 to 6) was presented for two consecutive 4.5 s trials (9 s total; 6 TRs). Loads progressed sequentially from 1 to 6 and then returned from 6 down to 1. One complete cycle comprised 24 trials (108 s; 72 TRs), presenting each load level four times. Each 540 s (9 min; 360 TRs) scanning run repeated this cycle five times.

To ensure stable behavioral performance during scanning, participants underwent extensive pre-scan training across 1-hour sessions over several days (typically 4–6 total hours; 16–24 practice runs). Using an adaptive staircase procedure (Watson and Pelli, 1983) with trial order and timing identical to the fMRI experiment, we estimated the orientation discrimination threshold (75 correct) for each VSTM load. Thresholds were calculated over 32 trials per load (8 cycles; 14 min 24 s per practice run; un-responded trials were repeated at the end of the run). Training continued until threshold estimates stabilized below 90° for each load. Final scan thresholds for each load and participant (Fig. S1A) were derived from a linear fit to threshold estimates across the last 6 practice runs.

To separate effects of task difficulty from task performance accuracy and orientation change extent, the main fMRI experiment included two task conditions, alternated between scanning runs:

1) Constant Performance Condition: The probe-target orientation difference was set to the participant’s load-specific threshold established during practice, maintaining equal task performance across load levels (Fig. S1B).
1) Constant Orientation Difference Condition: The orientation difference was held constant across all loads at the participant’s mean threshold averaged across all load levels, allowing task accuracy to decrease with load (Fig. S1C).

Participants were informed of the condition prior to each run. To minimize potential hemodynamic lag or neural adaptation effects caused by sequential load presentations, encoding models were fit across combined ascending and descending load sequences, counterbalancing the influence of preceding trial history^18^.

To test whether VSTM load systematically affected task performance in each condition, accuracy was analyzed using a linear mixed-effects model with VSTM load as a continuous fixed factor and participant as a categorical random factor (without an interaction term). Statistical significance across all analyses was set at *p* < 0.05.

### Control experiments

We conducted two control experiments to determine whether load-tuned responses reflected physical stimulus complexity or trial sequence predictability.

#### 1) Constant VSTM Load / Reverse Display Role Control

To test whether tuning was driven by visual stimulus differences across Memory Displays, three participants (P1, P4, P6) completed a “reverse display role” control task (Video S2). Physical visual stimulation was identical to the main experiment (using the orientation thresholds from the constant performance condition), but the display roles were reversed: participants memorized the single probe bar and judged whether its orientation matched any item in the subsequent six-item array. VSTM load was therefore held constant at 1 item across all trials despite variations in display complexity.

#### 2) Unpredictable Trial Order Control

To evaluate whether responses depended on predictable load sequences, trial history, load expectation, or hemodynamic lag artifacts, five participants (P1, P3, P4, P5, P6) completed an “unpredictable trial order” control experiment (Video S3).

Trial sequences followed a Type-1, Index-1 counterbalanced sequence^53,54^ across loads. In each 36-trial cycle (162 s; 108 TRs), every VSTM load level appeared six times and was preceded and followed by every load level (including itself) exactly once, eliminating trial-to-trial load correlations. Each 486 s (8.1 min; 324 TRs) scanning run contained three repeats of a cycle. To prevent sequence learning across six scanning runs per session, six distinct sequences remapped load labels to different numerical values. Because pseudo-random sequences have inherently lower statistical power for fMRI model estimation than progressive designs^28^ and could not be repeatedly averaged (each sequence was presented 9 times total per participant, compared to ∼80 repeats of the progressive cycle in the main experiment), this experiment served as a stringent out-of-sample validation test.

### Behavioral switching cost analysis in unpredictable trial order control

To test whether different VSTM loads recruit distinct cognitive strategies rather than continuously scaling effort, we analyzed trial-to-trial behavioral accuracy in the unpredictable trial order control experiment.

Because orientation differences were calibrated to yield equal baseline accuracy across loads, and sequence counterbalancing presented every load transition pair with equal frequency, the effect of preceding trial history could be isolated.

For each participant, response accuracy was calculated across trials grouped by the absolute difference in VSTM load relative to the preceding trial (|Δload| = |load*_t_*-load*_t-1_*|). To test for transition costs, accuracy was modeled using a linear mixed-effects model with absolute VSTM load change as a continuous fixed factor and participant as a random categorical factor (intercept-only, no interaction term). Additional analyses also included the direction of load change as a categorical fixed factor, including an interaction with absolute load change.

### Magnetic resonance imaging MRI data acquisition

Data were acquired on a 7T Philips Achieva scanner using a 32-channel head coil. In an initial session, a *T*_1_-weighted whole-brain anatomical scan was acquired for each participant. Anatomical images were automatically segmented using FreeSurfer and manually edited in ITK-SNAP to correct grey-white matter boundary errors, yielding high-fidelity 3D cortical surface reconstructions.

*T*_2_*-weighted functional images were acquired with a gradient-echo EPI sequence at an isotropic resolution of 1.7 mm (57 interleaved slices; 128×128 matrix; field of view: 216×216×97 mm). Sequence parameters were:

- TR / TE: 1500 ms / 22.5 ms
- Flip Angle: 70°
- Acceleration: SENSE factor 2.4, Multiband factor 3
- Phase Encoding: Anterior–Posterior
- Shimming: 2^nd^ order image-based *B*_0_ shim across the functional FOV (MRCodeTool, Zaltbommel, Netherlands).

Coverage spanned the fronto-parieto-occipital cortex, excluding the cerebellum and anterior temporal lobes to avoid 7T susceptibility artifacts and distortions.

### Run structure

- Progressive Load & Reverse Display Role Control Runs (558 s total): Each run began with 12 dummy scans (18 s) to achieve steady-state magnetization (discarded), followed by 360 functional TRs (540 s). During the final 6 dummy scans (9 s), the final two trials of the load sequence were displayed to allow models to account for hemodynamic response lag at the start and end of the analyzed time series. Participants completed 6 to 10 runs per session across two sessions.
- Unpredictable Order Control Runs (510 s total): Each run began with 16 dummy scans (24 s) followed by 324 functional TRs (486 s). The final 9 dummy scans (13.5\text) presented the final three trials of the load sequence to model hemodynamic lag. Participants completed 6 runs per session across three sessions.

### Pre-processing, Distortion Correction, and Denoising

In each session, a top-up scan (10 TRs) with reversed phase-encoding direction (P–A) was acquired to correct susceptibility-induced distortions^55^. A session-specific 1.7 mm *T*_1_-weighted anatomical scan matching the functional slice placement was acquired to facilitate coregistration to the high-resolution whole-brain anatomical model.

To suppress thermal noise, fMRI time series were denoised using NORDIC^56^ for conditions where complex (phase) data were collected. Specifically, NORDIC denoising was applied to all scanning runs in participants P1, P2, and P5, and to the unpredictable trial order condition in participant P3. For all remaining runs and participants, standard magnitude images were analyzed without NORDIC.

### Preprocessing and functional data alignment

Functional data were co-registered to the high-resolution whole-brain *T*_1_-weighted anatomical space using a custom pipeline^35^ implemented in AFNI^57^. To minimize blurring and spatial interpolation artifacts, all alignment, motion correction, and distortion correction transformations were concatenated into a single transformation matrix, reducing functional resampling to a single interpolation step. No additional spatial or temporal smoothing was applied.

The registration pipeline for each session proceeded as follows:

1) Anatomical Alignment: The session-matched *T*_1_ scan was aligned to the high-resolution 1 mm whole-brain *T*_1_ anatomical space (3dUnifize, 3dAllineate).
2) Motion Correction: Volume registration was performed on functional series acquired in both phase-encoding directions (3dvolreg).
3) Distortion Correction: Nonlinear distortion transformation matrices were computed from the average of the functional and reversed phase-encoding series (3dQwarp).
4) Functional Co-registration: Within-session and between-scan motion parameters (3dNwarpApply) were combined with the registration matrix linking the functional space to the session-matched *T*_1_ anatomical volume (3dvolreg).
5) Single-Step Transformation: The composite product of these transformation matrices was applied to every functional volume, projecting all time series directly into the high-resolution anatomical space.

### Data aggregation, cycle averaging, and cross-validation splits

Preprocessed time series were imported into Vistasoft (mrVista) for signal averaging and model fitting:

- Main Experiment Data: Functional runs were averaged separately for the constant performance and constant orientation difference conditions. To evaluate domain-general load representations, both conditions were also combined into a single averaged dataset. For cross-validation, data were split into independent odd-scan and even-scan averages, ensuring equal numbers of constant performance and constant orientation difference runs in each split. Within each run type, stimulus cycles (72 TRs, 108 s) were averaged together to yield a single 72-TR response profile per voxel for encoding model fitting.
- Unpredictable Trial Order Control Data: Time series were averaged across identical pseudo-random sequences (3 cycles per run, across 3 sessions) to generate six distinct 36-trial cycle averages (one for each sequence variant). Encoding models were then fit simultaneously across all six concatenated cycle averages to capture response dynamics across diverse trial histories.

### fMRI forward modeling and candidate neural response functions

To characterize voxel-level response profiles, we evaluated forward neural encoding models that predict fMRI time series based on VSTM load^19,50,58^. Our modeling strategy involved two distinct steps: (1) Model Selection, comparing competing candidate neural response functions (functional forms) across the cortex; and (2) Parameter Estimation, fitting voxel-specific response parameters for each candidate function.

Following population encoding frameworks in event timing^19^ and visual numerosity^18,22,27,35^, we tested candidate functional forms that predict neural responses as either monotonic or tuned functions of VSTM load, evaluated on both linear and logarithmic scales.

#### 1) Candidate Functional Forms (Model Selection)

We evaluated five candidate neural response functions to determine which model family best accounts for voxel time courses:

- Linear Monotonic: Predicts neural response amplitude scales linearly with VSTM load:

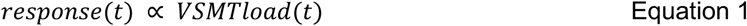
- Logarithmic Monotonic: Predicts neural response amplitude scales with the logarithm of VSTM load:

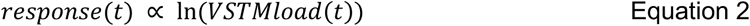
- Power-Law Monotonic: Predicts a compressive monotonic response function of VSTM load:

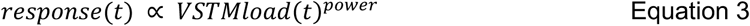
- Linear-Gaussian Tuned: Predicts selective tuning following a Gaussian profile on a linear VSTM load scale:

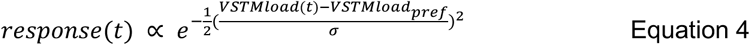
- Logarithmic-Gaussian Tuned: Predicts selective tuning following a Gaussian profile on a logarithmic VSTM load scale:

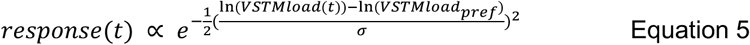

#### 2) Free Parameters Fit Per Voxel (Parameter Estimation)

For each candidate functional form independently, we solved for the optimal voxel-specific free parameters:

- Fixed/No Free Parameters (Eq. 1 & 2): Linear and logarithmic monotonic models have no shape parameters, predicting fixed scaling relative to load.
- Exponent Parameter (Eq. 3): The power-law model estimates a single free shape parameter, *power* (>0), governing response compression/expansion.
- Tuning Parameters (Eq. 4 & 5): The Gaussian tuned models estimate two free shape parameters for each voxel:

1) *VSTMload_pref_*: The preferred VSTM load (the peak or center of the tuning curve).
2) *σ*: The tuning width (standard deviation of the Gaussian profile), governing load selectivity.

Additionally, for all models during time-series prediction, an overall gain/amplitude scaling factor (g) was fit to map predicted neural responses to the observed fMRI signal.

### Candidate response model fitting and cross-validated model comparison

#### 1) Model Fitting Pipeline

Model fitting was executed using population receptive field (pRF) modeling principles^59^ adapted for the VSTM load dimension. For each candidate neural response function and each combination of candidate free parameters:

1) Neural Time Course Generation: The neural response function’s amplitude was evaluated at each stimulus time point based on the VSTM load presented in that trial, generating a predicted neural response time course across the scan.
2) Hemodynamic Convolution: The predicted neural time course was convolved with a canonical hemodynamic response function^60^ to yield a predicted BOLD fMRI response time course.
3) Parameter Optimization: A dense grid-search optimization evaluated candidate parameter combinations across a wide parameter space. For each voxel, the parameter set that minimized the sum of squared residuals between the predicted and observed fMRI time series, thereby maximizing response variance explained (*R*^2^), was selected.

#### 2) Cross-Validation and Model Selection

To compare candidate response models with differing numbers of free parameters without overfitting bias, we used a two-fold cross-validation procedure. For each voxel, parameters were estimated using the odd-numbered scan runs (or Constant Performance Condition runs) and evaluated by calculating the cross-validated variance explained (*R*^2^*_CV_*) in the observed fMRI time series of the even-numbered scan runs (or Constant Orientation Difference Condition runs), and vice versa.

For statistical model comparison across candidate functional forms:

- Initial Voxel Selection: Voxels were included where any candidate response model yielded a cross-validated variance explained of *R*^2^*_CV_* > 0.2 (for main experiment data) or *R*^2^*_CV_* > 0.1 (for Unpredictable Trial Order Control Data).
- Median Summary Statistics: Within these selected voxels, the median *R*^2^*_CV_* was computed for each candidate neural response function across each region of interest (ROI), hemisphere, and cross-validation split (Fig. S4A, C, E, G, I).
- Statistical Hypothesis Testing: Paired two-sided *t*-tests were conducted on these median values to determine statistical differences in goodness-of-fit between competing candidate response functions, yielding *t*-statistics and *p*-values across model comparisons (Fig. S4B, D, F, H, J).

#### 3) Winning Model Selection and Voxel Inclusion Criteria

Statistical comparisons revealed that the logarithmic-Gaussian tuned response function predicted fMRI time series significantly better than all competing monotonic and linear-Gaussian candidate models.

Consequently, voxel-level parameter estimates (*VSTMload_pref_*, *σ, R*^2^*_CV_*) from this log-Gaussian model were used for all subsequent spatial and organizational analyses.

### Region of interest identification

By projecting each voxel’s variance explained from its best fitting model and VSTM load preference from the logarithmic-Gaussian model onto reconstructed cortical surface models, we defined regions of interest representing topographic maps of VSTM load. Following established retinotopic mapping protocols^61^, maps were identified as contiguous, responsive cortical areas showing higher variance explained than surrounding locations (Fig. S6), localized consistently relative to major anatomical sulci across all hemispheres (Fig. S8), and containing a single, continuous progression of VSTM load preferences across the cortical surface (Fig. 2, S7).

We identified 11 distinct maps per hemisphere (22 per participant), yielding a total of 132 map instances across the cohort. For each map, two boundary lines were manually drawn on the cortical surface to demarcate the low VSTM load preference end and high VSTM load preference end, enabling quantitative distance analyses along the topographic gradient.

### Voxel Inclusion Criteria

For subsequent analyses, voxels were selected based on two stringent inclusion criteria:

- Goodness of Fit: *R*^2^ > 0.2 for the winning log-Gaussian model.
- Tuning Preference Within Range: Estimated VSTM load preference falling within the tested stimulus range (1< *VSTMload_pref_*<6). Voxels with estimated tuning peaks outside this range exhibit monotonic response scaling across the tested loads and thus do not show convincing evidence of selective tuning.

### Model validation via ground-truth fMRI simulations

To evaluate the sensitivity, specificity, and parameter estimation accuracy of our model fitting and cross-validation pipeline, we conducted simulations with known ground-truth neural response functions.

#### 1) Ground-Truth Data Generation and Noise Degradation

Synthetic fMRI time series were generated for four ground-truth response profiles: linear-monotonic, logarithmic-monotonic, linear-Gaussian tuned, and logarithmic-Gaussian tuned. For tuned functions, ground-truth shape parameters (*VSTMload_pref_* and *σ*) were drawn from broad, uniform distributions across the parameter space.

Simulation time series were processed as follows:

1) Normalization: Synthetic time courses were z-scored (zero mean, unit standard deviation).
2) Noise Injection: Normally distributed Gaussian noise (standard deviations ranging from 0 to 6) was added to yield signal-to-noise ratios (SNR) from 1/6 to infinity (noiseless).
3) Cross-Validation Paired Datasets: Independent noise set were generated over identical underlying signals to create paired time-series splits for two-fold cross-validation.
4) Rescaling: Simulated time series were re-scaled using the empirical mean and standard deviation of BOLD responses observed in our cortical maps, matching the amplitude dynamic range of human 7T fMRI data.

#### 2) Model Recovery and Classification Performance

Simulated datasets were processed through the identical model fitting, grid-search, and cross-validation pipelines used for experimental fMRI data. Cross-validated variance explained (*R*^2^*_CV_* was compared across all candidate models for each ground-truth response type.

Classification accuracy was evaluated separately for:

- In-Range Preferences (1<*VSTMload_pref_* <6): Categorized as valid tuned responses.
- Out-of-Range Preferences (*VSTMload_pref_* < 1 or > 6): Categorized as monotonic edge-case responses outside the tested load window.

Binary model classification accuracy was computed as a function of both SNR and *R*^2^*_CV_*.

#### 3) Parameter Estimation Bias and Error Mapping

Because fMRI responses exhibitted non-uniform parameter distributions, we systematically evaluated parameter recovery for voxels meeting our empirical inclusion threshold (*R*^2^*_CV_*>0.2).

For every parameter combination across the search grid, we calculated:

- Classification Accuracy: The proportion of correctly identified response models across combinations of estimated parameters.
- Parameter Estimation Bias: The signed difference between estimated and ground-truth parameters (Estimate - Ground Truth) for both *VSTMload_pref_* and *σ*.

### Surface template registration and standardized map coordinates

To evaluate group-level topographic overlap and establish standardized coordinates across participants:

- Group Overlap Mapping: Individual cortical surface models were co-registered to the N27 (Talairach) standard surface template using AFNI (3dQwarp, 3dAllineate). Cortical map ROIs from all participants were projected into this common surface space, and vertex-wise overlap density was calculated across hemispheres (Fig. S8).
- Centroid Coordinates: The geometrical center (centroid) of each map was manually identified on individual cortical surface reconstructions and transformed into Talairach space using AFNI. Centroids were subsequently converted into MNI coordinate space using SPM’s tal2mni tool. Mean coordinates and standard deviations were computed for each map across the cohort (Table S1).

### Spatial reliability and topographic progression analyses within ROIs

#### 1) Repeatability of VSTM Load Preference Patterns

To test whether voxel-level load preferences showed consistent spatial distributions across independent measurements (independent of specific spatial structure) we calculated non-parametric Spearman rank correlations between load preferences estimated from:

1) Odd-scan vs. Even-scan runs
2) Constant Performance vs. Constant Orientation Difference runs

To account for spatial oversampling when transforming data from the functional acquisition resolution (1.7³ mm³) to the higher-resolution cortical surface grid (1.0 mm³), *p*-values were adjusted for surface upsampling. Specifically, the number of surface vertices in each ROI was divided by a surface upsampling factor of 2.89 (1.7^2^, accounting for sampling restricted to a 2D folded surface) prior to converting rank correlations to *p*-values. All resulting *p*-values were corrected for multiple comparisons across all maps and hemispheres using False Discovery Rate (FDR^62^).

#### 2) Quantifying Topographic Progressions Across Cortical Surface Distance

To characterize the spatial gradient of load preferences between the defined low-preference and high-preference boundary lines within each map:

1) Cortical Distance Calculation: For every voxel within an ROI, geodesic distances along the cortical surface to the nearest point on the low-end line (*d*_low_) and high-end line (*d*_high_) were calculated.
2) Normalized Distance Metric: Cortical distance along the map progression was computed as:

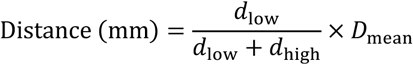 where *D*_mean_ is the mean surface distance between the two boundary lines^18,22^. This normalizes cortical distance in millimeters while accounting for non-parallel boundary lines across cortical folds.
3) Topographic Gradient Correlation: Spearman rank correlations were computed between voxel cortical distances and their estimated VSTM load preferences (*VSTMload_pref_*).

For visualization (Figs. 3A, S11), load preferences were binned in 2 mm cortical distance increments, plotting bin means and standard errors. Linear regression functions were fit to these binned means, and 95% confidence intervals were constructed using 10,000 bootstrap iterations.

#### 3) Circularity Controls

Because ROI ends were initially identified based on gradient continuity, assessing topographic gradients within the primary dataset introduces potential circularity. To rigorously confirm gradient repeatability in independent data:

1) Independent Data Splits: Topographic distance correlations were re-computed using VSTM load preferences estimated from independent data subsets (*odd runs*, *even runs*, *constant performance runs*, and *constant orientation difference runs*).
2) Independent ROI Definitions: ROI borders and ends were identified from odd and even run data subsets and correlations re-computed using VSTM load preferences estimated from the complementary subset.

### Topographic map orientations relative to cortical anatomy

To quantify the orientation of load-preference progressions relative to major anatomical structures (sulci and gyri)^31^:

1) Gradient Vector Computation: For each surface vertex in a map, pairwise cortical distances and signed differences in VSTM load preference (Δ*VSTMload_pref_*) were calculated relative to all other vertices within that map. A weighted average of these directional vectors was calculated, where vector magnitudes were weighted by the local slope ((Δ*VSTMload_pref_*/cortical distance). This yielded a vertex-wise vector representing the direction of steepest increase in load preference.
2) Anatomical Alignment: The direction of the shortest cortical surface path from each vertex to the nearest sulcal fundus or gyral crown was determined. To compute the angular difference between the functional gradient and this anatomical trajectory, we evaluated the cross product of each direction vector with the surface normal at that vertex.
3) Hemispheric Normalization and Circular Statistics: Angular differences were averaged across all vertices within each map per hemisphere. To correct for mirror symmetry across hemispheres, the sign of angular differences in the right hemisphere was inverted. Rayleigh’s test for circular uniformity^63^ was applied to test whether each maps orientations across hemispheres followed a directional von Mises (circular Gaussian) distribution rather than a uniform distribution, FDR-correcting for multiple map comparisons across all maps.

### Relationships between VSTM load preferences and tuning widths within ROIs

To characterize how tuning selectivity changes as preferred VSTM load increases across cortical maps:

1) For each responsive voxel in a map, tuning width was quantified as the Full Width at Half Maximum (FWHM) of its logarithmic-Gaussian response function. FWHM was determined as the difference between the upper and lower VSTM load values where response amplitude dropped to 50% of its peak value.
2) Preference vs. Width Rank Correlations: Non-parametric Spearman rank correlations were calculated between voxel VSTM load preferences (*VSTMload_pref_*) and FWHM tuning widths, for each map considering voxels from all hemisphere. To correct for spatial surface upsampling, vertex counts per map were divided by the surface upsampling factor of 2.89 (1.7^2^) prior to computing *p*-values. All resulting *p*-values were FDR-corrected across all maps.
3) Visualization & Trend Fitting: For visualization (Fig. S13), voxels were binned in *VSTMload_pref_* increments of 0.25, computing bin means and standard errors. Linear regression functions were fit to these bin means, with 95% confidence intervals established via bootstrapping across 10,000 iterations.

### Relationship between VSTM load preferences and model goodness of fit

To test whether model fits varied systematically as a function of preferred load within maps, we evaluated voxel-level relationships between preferred VSTM load and cross-validated variance explained for the logarithmic-Gaussian tuned model. Within each map, voxels were selected using standard inclusion criteria (*R*^2^ >0.2, 1< *VSTMload_pref_*<6). Spearman rank correlations were computed between *VSTMload_pref_* and and *R*^2^. Resulting *p*-values were adjusted for surface upsampling and FDR-corrected across all map instances.

### Between-ROI hierarchical difference analyses

To evaluate hierarchical transformations across the cortical map hierarchy, six key map-level parameters were quantified for each map instance:

1) Mean VSTM Load Preference: Averaged *VSTMload_pref_* across tuned voxels (Fig. 4A).
2) Mean Tuning Width: Averaged FWHM (Fig. 4B).
3) Mean Goodness of Fit: Averaged *R*^2^ for the log-Gaussian model (Fig. 4C).
4) Responsive Surface Area: Total cortical surface area (mm^2^) occupied by responsive voxels (*R*^2^ >0.2, 1< *VSTMload_pref_*<6) within the ROI (Fig. 4D).
5) Tuning Advantage: Mean cross-validated variance explained advantage (Δ*R*^2^*_CV_*= *R*^2^*_CV_*_, log-Gaussian_ -*R*^2^*_CV_*_, log-monotonic_) evaluated across odd and even splits (Fig. 4E).
6) Retinotopic Overlap Fraction: Proportion of VSTM load-tuned voxels that also exhibited significant spatial tuning in independent visual field mapping datasets (*R*^2^>0.1; Fig. 4F).

### Statistical Modeling of Hierarchical Differences

To evaluate differences in these metrics across maps while controlling for subject-level baseline variation:

- Linear Mixed-Effects Models: Analyzed each metric using models with map identity as a fixed categorical factor and participant as a random categorical factor (intercept-only, no interaction term).
- Degrees of Freedom & Post-Hoc Tests: Marginal tests for fixed effects were evaluated using the Satterthwaite approximation for degrees of freedom^64,65^. Pairwise differences between maps were assessed using Tukey’s Honestly Significant Difference (HSD) post-hoc tests^66^.
- Tuned vs. Monotonic Comparison: Differences in cross-validated variance explained between tuned and monotonic model fits across maps were evaluated using two-sided Wilcoxon signed-rank tests followed by FDR correction.

### Numerosity and visual field position preference comparisons

To determine whether VSTM load preferences were independent of other topographically mapped dimensions, load preferences were compared against visual spatial position (eccentricity, polar angle, *R*^2^) and numerosity preferences within the same cortical locations.

Data sources:

- Numerosity Mapping Data: Retrieved for participants P3, P4, and P6 from published datasets^22,35,67^.
- Visual Field Mapping Data: Retrieved for participants P4 and P6 from existing datasets^22,35^. New retinotopic mapping data were acquired for participants P1, P3, and P5 using standard sweeping bar stimuli and pRF modeling protocols^21^ with fMRI sequence and preprocessing parameters identical to the main experiment.

Voxel Selection & Inclusion Thresholds:

- *VSTM Load vs. Numerosity Comparison:* Included voxels showing evidence of selective tuning for both dimensions, defined as *R*^2^>0.1 for both models, 1< *VSTMload_pref_*<6 and numerosity preference between 1 and 7 items.
- *Visual Spatial Response Comparison:* Included voxels with variance explained *R*^2^>0.1 for both models, 1< *VSTMload_pref_*<6 and and spatial eccentricity preference within the stimulated field of view (6.5°).

### Statistical Analyses & Multiple Comparison Corrections

Non-parametric Spearman rank correlations were calculated within each VSTM load map between VSTM load preferences and:

1) Numerosity preferences
2) Spatial eccentricity preferences
3) Spatial polar angle preferences

Additionally, the variance explained (*R*^2^) of VSTM load response models was compared to that of visual field mapping models using the same correlation framework. All *p*-values were converted from correlation coefficients after adjusting for surface upsampling and were FDR-corrected across all map instances.

### Relationships between responses in the main and control experiments

To confirm that neural response models reflected internal VSTM load demands rather than sequence predictability or visual display features, we evaluated the consistency of estimated VSTM load preferences between the main experiment and the two control conditions (*unpredictable trial order* and *reverse display role*).

- Voxel Selection Thresholds: For these cross-experiment comparisons, inclusion criteria were set to *R*^2^>0.1 and 1<*VSTMload_pref_*<6 in both compared datasets. This threshold accounted for the lower statistical power inherent to pseudo-random sequences and reduced repeat counts in the control designs.

Cross-Experiment Correlation Analysis: Within each map instance per hemisphere, non-parametric Spearman rank correlations were calculated between the VSTM load preferences estimated from the main experiment and those estimated from each control experiment. All *p*-values were again adjusted for cortical surface upsampling and FDR-corrected.

## Resource availability

### Lead contact

Ben M. Harvey

### Materials availability

No materials were generated in this study.

### Data and code availability

Ethical constraints prevent us from sharing the medical imaging data sets (MRI scans) generated in the current study to public repositories. These raw data sets are available from the lead contact upon reasonable request, depending on agreements not to share these data publicly.

Response data and fit model parameters underlying all statistical analyses are publicly available at: - https://doi.org/10.6084/m9.figshare.32779608

The code that supports the findings of this study is available from the following repositories:

- vistasoft (https://github.com/vistalab/vistasoft)
- vistasoftAddOns (https://github.com/benharvey/vistasoftAddOns)
- fMRI_preproc (https://github.com/MvaOosterhuis/fMRI_preproc)
- AnalysisScript (https://github.com/MvaOosterhuis/VSTM_Analysis)

## Acknowledgments

Funding by Netherlands Organization for Scientific Research grant 452.17.012 (BMH).

## Author contributions

Conceptualization: B.M.H., J.H., J.M.P.

Methodology, Formal analysis: B.M.H., M.A., J.H., J.M.P. Software: B.M.H., M.A., J.H.

Validation, Visualization: B.M.H., M.A. Investigation: B.M.H., M.A., E.H., J.M.P. Data curation: M.A.

Funding acquisition, Project Administration: B.M.H. Supervision: B.M.H., N.S., J.M.P.

Writing – original draft: B.M.H., M.A., S.G.

Writing – review & editing: B.M.H., J.H., E.H., N.S., S.G., J.M.P.

**Declaration of interests:** The authors declare no competing interests.

## Supplemental information

Document S1. Figures S1–S18, Tables S1-S13, captions for Videos S1-S3. Videos S1-S3.

### Declaration of generative AI and AI-assisted technologies in the manuscript preparation process

During the preparation of this work, the authors used Google Gemini 3.6 for text editing and journal-specific formatting. The authors reviewed and edited the output as needed and take full responsibility for the content of the published article.

