## Supplementary Materials for "A tuned topographic map hierarchy formats visual working memory load"

**Figures**

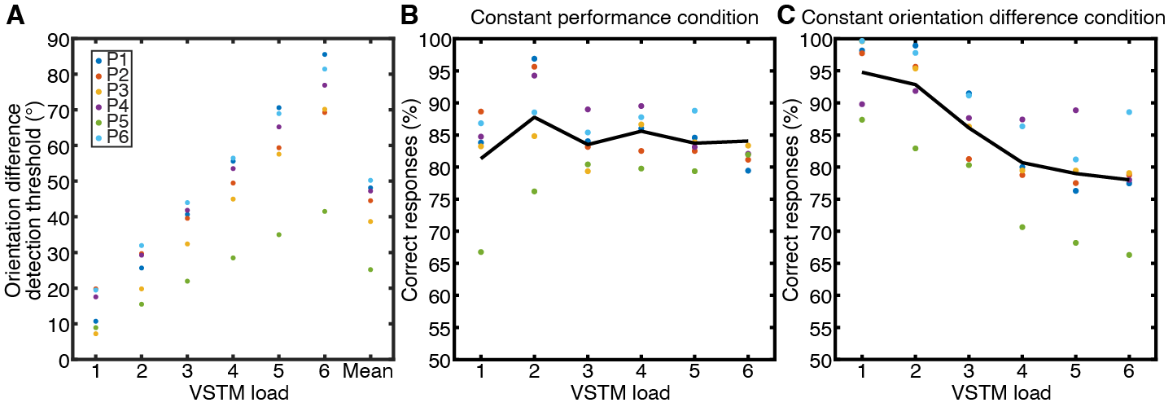


**Figure S1, Participants' task thresholds and accuracy at each VSTM load.** (**A**) Orientation difference thresholds measured before fMRI for each participant at each VSTM load. The mean across the six measured VSTM loads was use in the constant orientation difference condition. (**B**) In behavioral responses collected during fMRI in the constant performance condition, the task was performed at the measured orientation difference detection threshold for each VSTM load, so response accuracy was not significantly affected by VSTM load. Participants failed to respond on an additional 0.82% of trials, which are excluded here. (**C**) In the constant orientation difference condition, the task was performed with the same orientation difference regardless of VSTM load, so response accuracy decreased as VSTM load increased. Participants failed to respond on an additional 0.46% of trials.

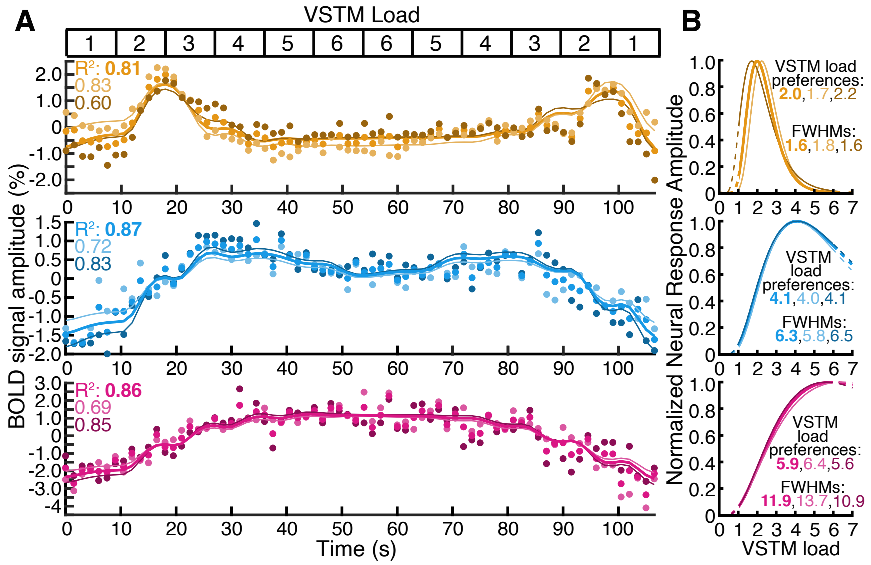


**Figure S2, Tuned fMRI responses to VSTM task load are similar between task conditions and repeated measurements, related to Figure 1C-D.** (**A**) Examples of observed fMRI blood-oxygen-level-dependent (BOLD) responses (dots) from three nearby recording sites with different VSTM load preferences in one participant's intraparietal sulcus, elicited by the sequence of changing VSTM loads (top inset). The BOLD responses were remarkably similar in the constant performance condition (light dots), constant orientation change condition (dark dots) and their average (saturated color dots). These responses were well captured by the VSTM load-tuned response model (light, dark and saturated lines respectively). (**B**) The best fitting neural response functions for each response (correspondingly colored lines) and the parameters of these response functions (colored text) were very similar between conditions. Dashed lines extend the response functions beyond the tested VSTM loads.

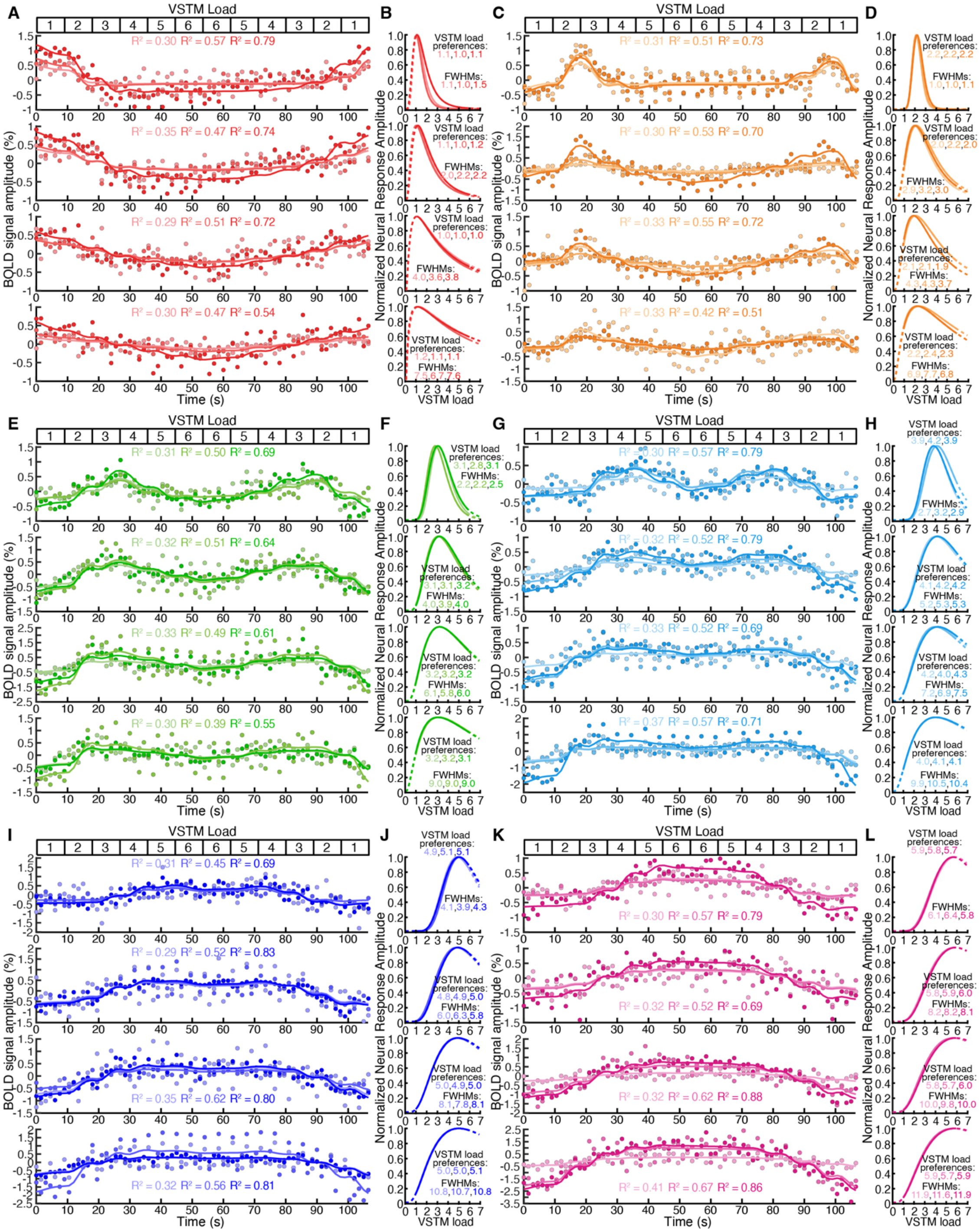

**Figure S3, Tuned fMRI responses to VSTM task load cover a broad range of VSTM load preference, tuning width and signal quality, related to Figure 1C-D.** (**A, C, E, G, I, K**) Examples of observed fMRI BOLD responses (dots) from recording sites with VSTM load preferences around 1, 2, 3, 4, 5 and 6 respectively, elicited by the sequence of changing VSTM loads (top inset). These responses were best captured by VSTM load-tuned response models whose predictions are shown as colored lines. Responses and fits with relatively low, medium and high variance explained ($R^2$) are shown with light, medium and dark dots and lines respectively. Tuning widths increase from top to bottom panels. (**B, D, F, H, J, L**) The best fitting neural response functions for each response (correspondingly colored lines) and the parameters of these response functions (colored text). All responses and fit parameters are in the online data.

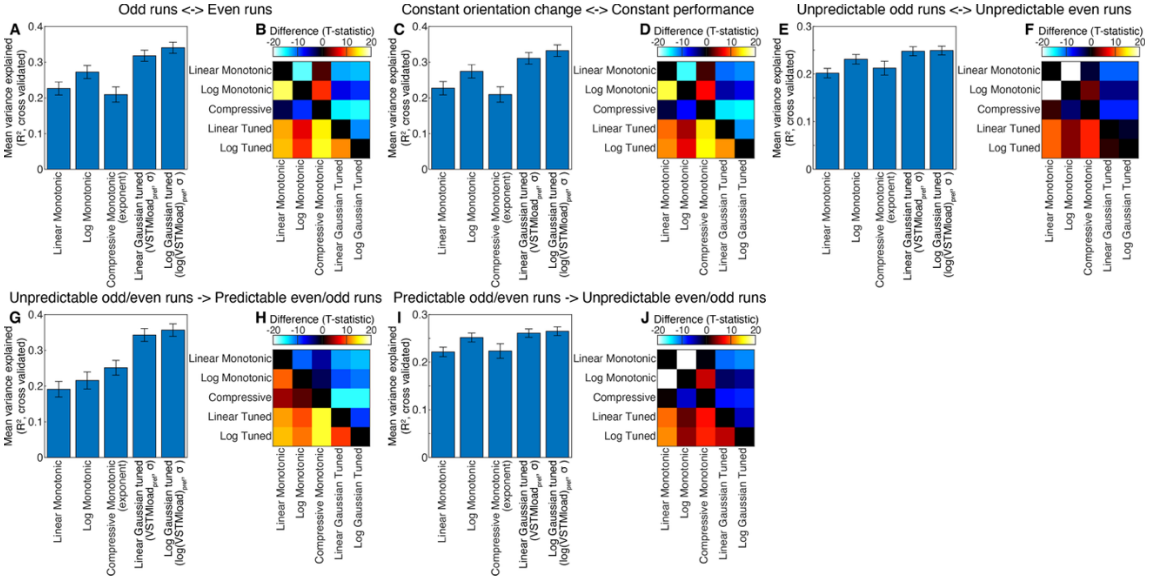


**Figure S4, Comparison of response variance explained by candidate Gaussian-tuned and monotonic models of neural population responses to VSTM load.** (**A**) Median cross-validated variance explained for candidate models (linear and logarithmic versions of Gaussian-tuned and monotonic functions, and a monotonic function with a free compressive exponent parameter) across all hemispheres, maps, and odd/even scan splits (n = 264). Error bars indicate the 95% confidence interval of the median. (**B**) T-statistics for the pairwise differences between candidate models matched for hemisphere, map, and cross-validation split. The log-Gaussian model outperformed all other models ($p<10^{-24}$, FDR-corrected for all comparisons). (**C**, **D**) Identical to (A) and (B), but evaluating median variance explained and model differences across constant orientation change and constant performance conditions. The log-Gaussian model outperformed all other models ($p<10^{-24}$). (**E**, **F**) Identical to (A) and (B), but for the unpredictable trial order control experiment (n = 220; a subset of participants). The log-Gaussian model outperformed all other models (p<=0.017). (**G**, **H**) Cross-experimental validation: Median variance explained for models fitted on the unpredictable trial order control experiment and evaluated on the main experiment (n = 220), along with corresponding pairwise differences. The log-Gaussian model outperformed all other models ($p<10^{-11}$). (**I**, **J**) Cross-experimental validation: Median variance explained for models fitted on the main experiment and evaluated on the unpredictable trial order control experiment, along with corresponding pairwise differences. The log-Gaussian model outperformed all other models ($p<10^{-10}$).

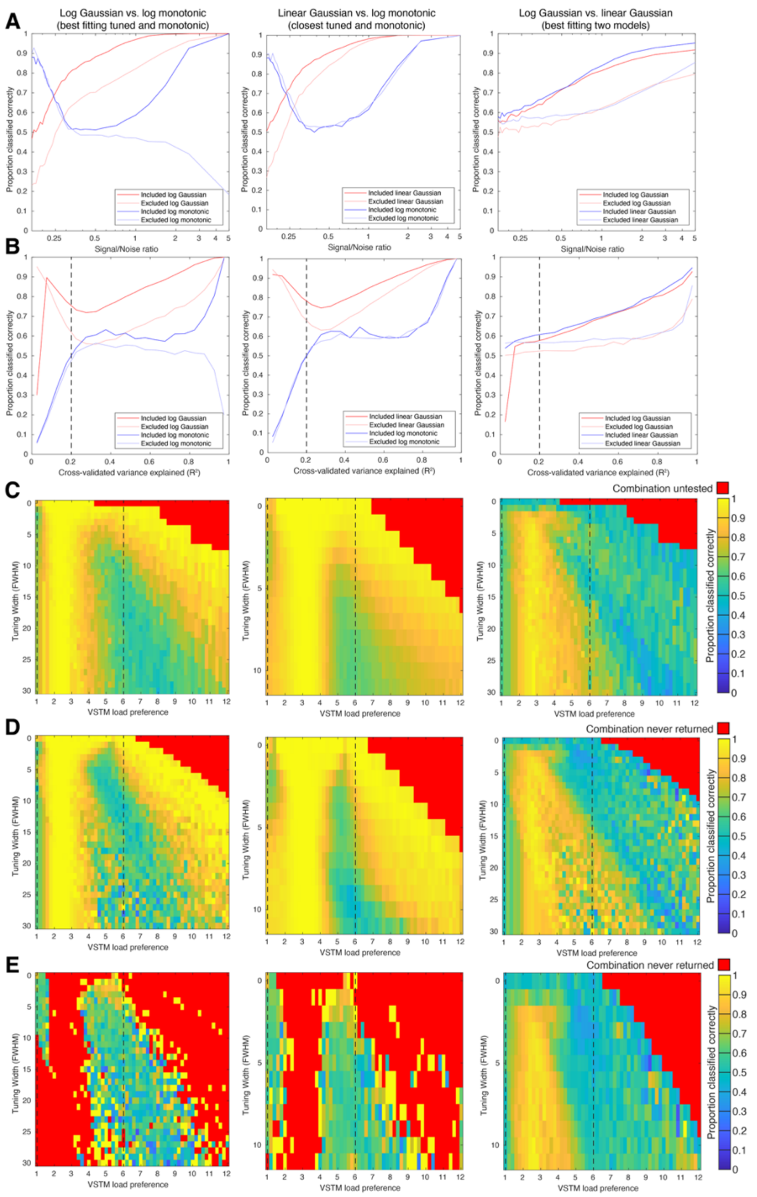


**Figure S5, Validation simulations reliably distinguish between simulated responses following different response functions within the parameter range included in subsequent analyses.** (**A**) Our modeling procedure correctly identifies ground-truth response functions, though classification accuracy decreases with higher noise. Accuracy is higher within the range of estimated VSTM load preferences included in subsequent analyses (dark colors) than in excluded ranges (light colors). (**B**) Classification accuracy decreases when cross-validated variance explained falls below our analysis threshold of 0.2. (**C**) Classification accuracy depends on the ground-truth parameters of the simulated response function (for the function listed first in the column title). Gaussian functions with VSTM load preferences below 4 are identified with high accuracy, whereas preferences near the edge of the tested VSTM load range (dashed lines) show lower accuracy. (**D**) Identical to (C), but shown as a function of estimated (rather than ground-truth) Gaussian parameters. (**E**) Classification accuracy for simulated responses following the second function listed in the title depends on the parameters estimated by the first model. Ground-truth monotonic response functions are rarely misidentified as Gaussian functions with VSTM load preferences below 4 (the range common in our fMRI data). However, monotonic responses are occasionally misidentified as Gaussian functions with high VSTM load preferences.

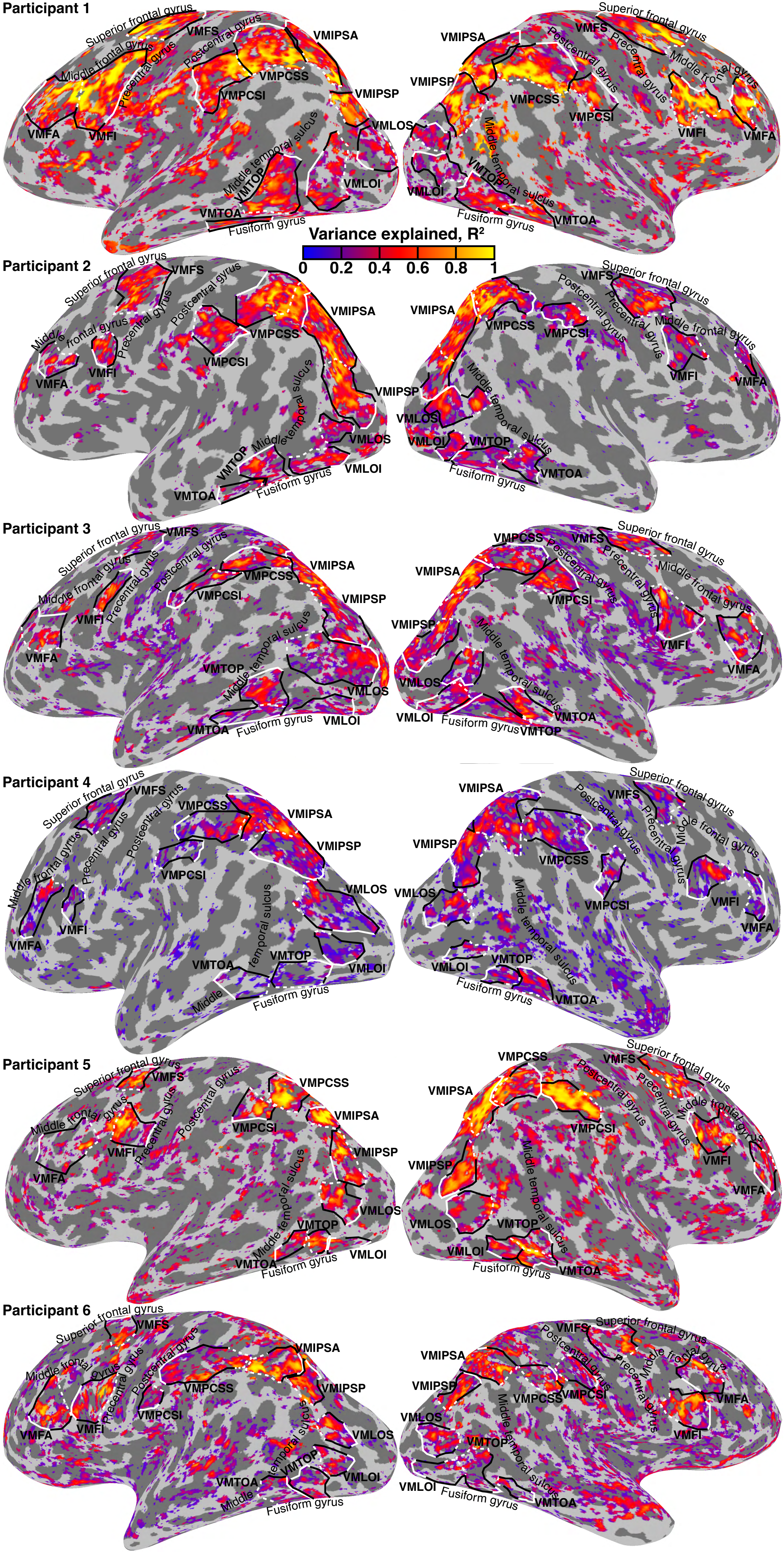


**Figure S6, Variance explained (R2) by the best fitting (logarithmic Gaussian tuned) response model projected onto the cortical surface of all hemispheres.** We observed consistent responsive locations in all participants. Participants showed individual differences in the goodness of model fits, resulting from both inherent individual differences in fMRI responses and improvements in our data acquisition protocol during the experiment. Therefore, participant-specific variance explained thresholds were used for ROI definition and visualization: 0.2 for P1, P2 and P5; 0.15 for P3 and P6; 0.1 for P4.Variance explained values (colors) are only shown for recording sites with convincing evidence of VSTM load tuning (preferences between 1 and 6, variance explained above these thresholds). The same threshold (0.2) was used for all quantitative analyses.

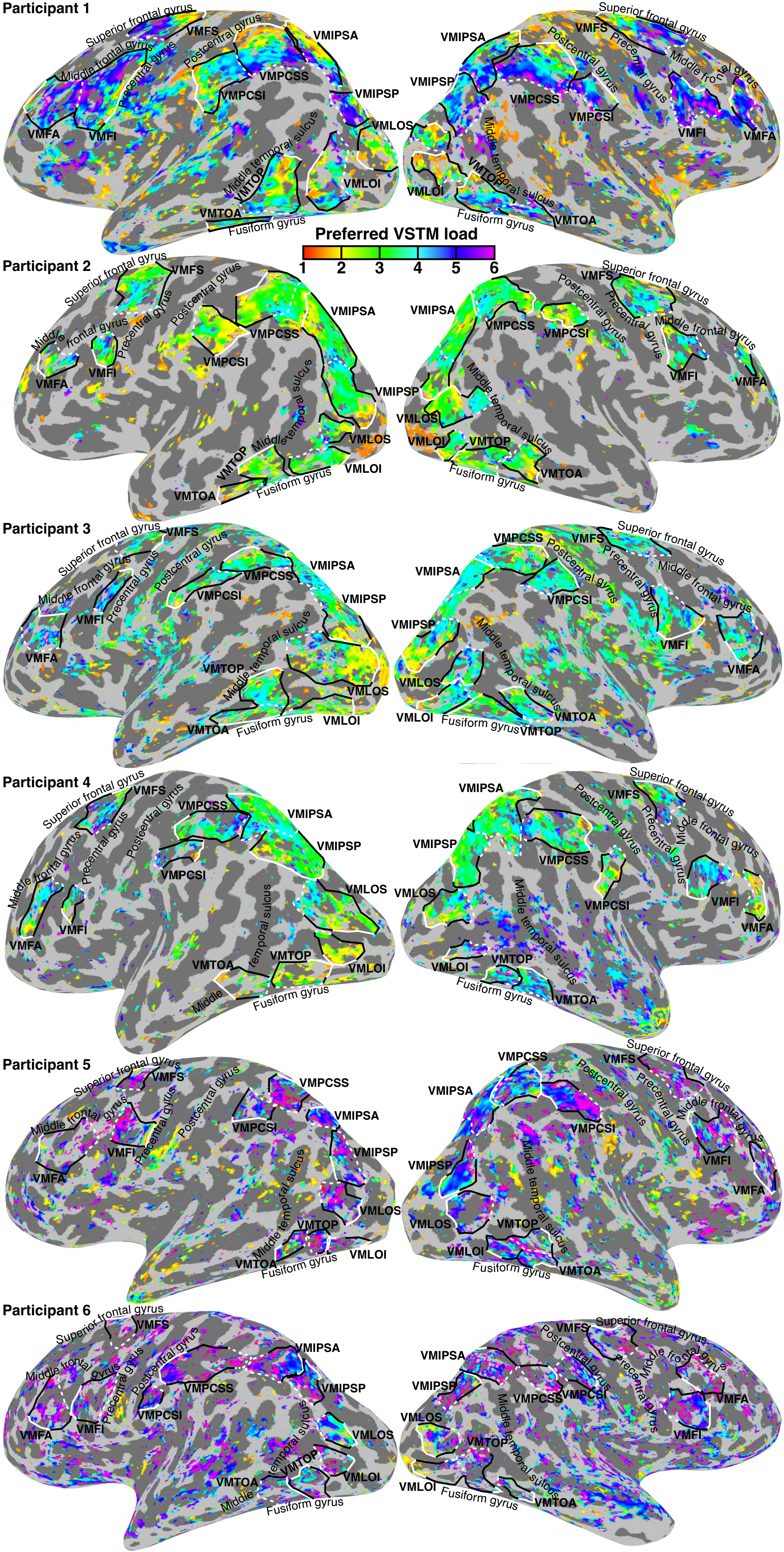


**Figure S7, VSTM load preferences from models fit to all data from the main experiment projected onto the cortical surface of all hemispheres, related to Figure 2.** Map borders are shown as black and white lines, with low and high VSTM load-preference ends in solid white and dashed white lines, respectively. Major gyri and sulci used for analysis of the orientation of topographic progressions are labelled in black text. Data are thresholded as described for Fig. S6. Image layout is consistent between Figs. S6, S7, S9, S10, S16, S17 and S18 allowing readers to compare responses between task conditions and control conditions by switching between pages.

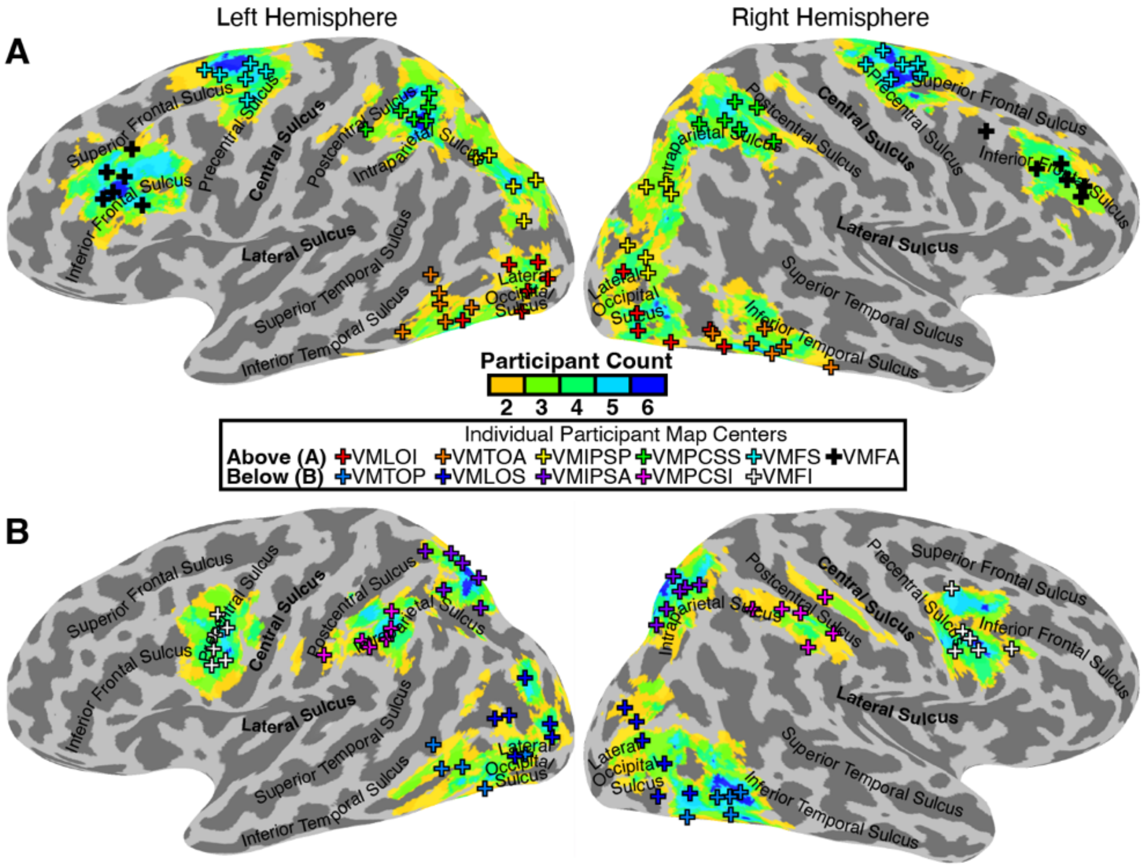


**Figure S8, Consistency and spatial overlap of VSTM load maps across participants.**
**(A)** Spatial overlap of 6 of the 11 identified VSTM load maps (VMLOI, VMTOA, VMIPSP, VMPCSS, VMFS, and VMFA) transformed onto an N27 template brain surface. Color scale indicates participant count per vertex. Colored crosses represent individual participant map centers. Maps are split across panels A and B to prevent overlapping boundaries from obscuring adjacent maps.
**(B)** Spatial overlap and individual participant centers for the remaining 5 interspersed maps (VMTOP, VMLOS, VMIPSA, VMPCSI, and VMFI).

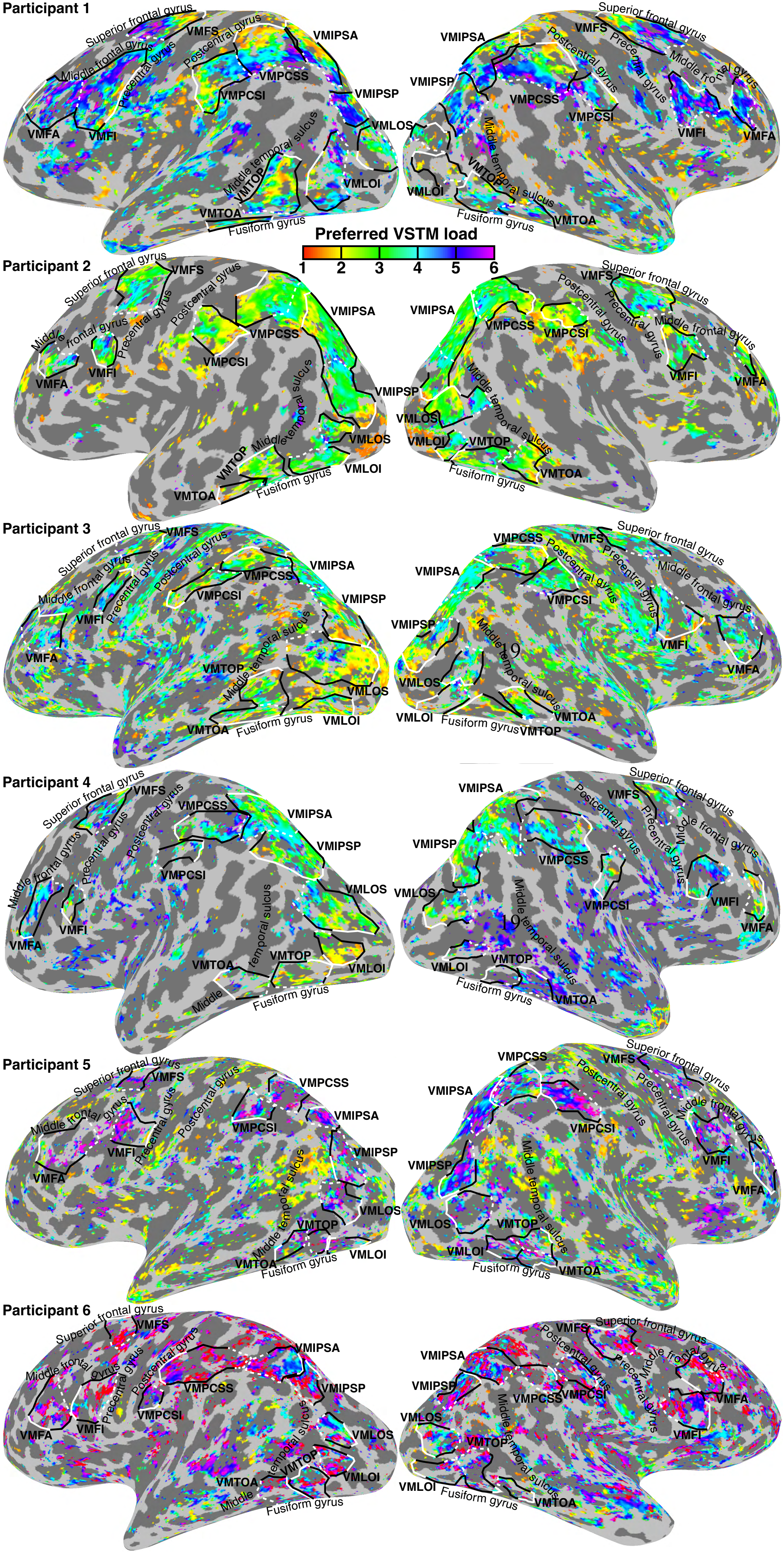


**Figure S9, VSTM load preferences from models fit to data from the constant performance condition, projected onto the cortical surface of all hemispheres, related to Figure 2.** Participant-specific variance explained thresholds were used for visualization: 0.2 for P1, P2 and P5; 0.15 for P3 and P6; 0.1 for P4. The same threshold (0.2) was used for all quantitative analyses. Map borders are taken from the models fit to all data (Fig. S7). These are shown as black and white lines, with low and high VSTM load-preference ends in solid white and dashed white lines, respectively.

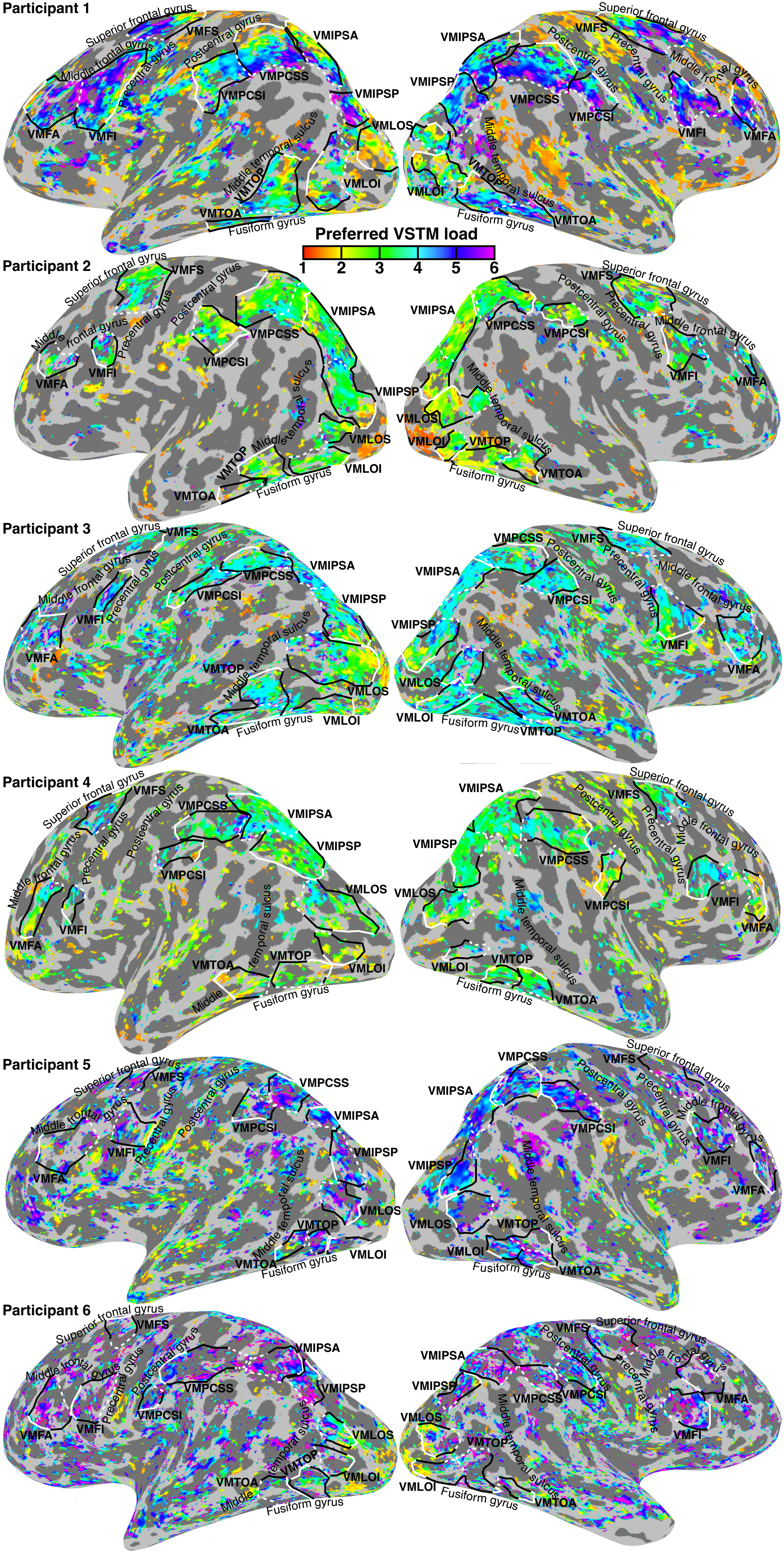


**Figure S10, VSTM load preferences from models fit to data from the constant orientation difference condition, projected onto the cortical surface of all hemispheres, related to Figure 2.** Participant-specific variance explained thresholds were used for visualization: 0.2 for P1, P2 and P5; 0.15 for P3 and P6; 0.1 for P4. The same threshold (0.2) was used for all quantitative analyses. Map borders are taken from the models fit to all data (Fig. S7). These are shown as black and white lines, with low and high VSTM load-preference ends in solid white and dashed white lines, respectively.

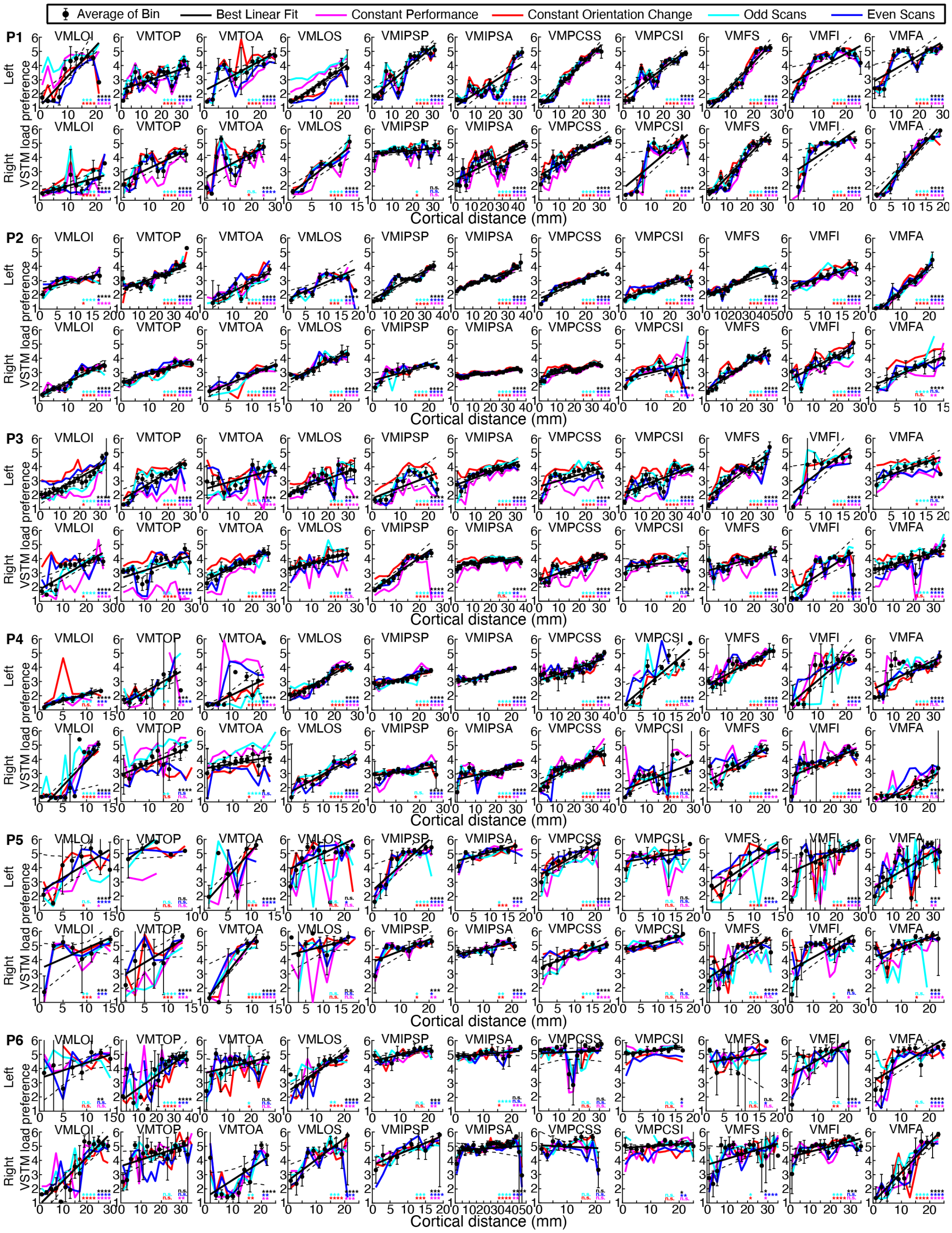


**Figure S11, VSTM load preferences were significantly correlated with cortical distance between the low and high load-preferring ends of each map in repeated measures, related to Figure 3A.** The same progression was found across repeated measures, and in both the constant performance and constant orientation difference task conditions (colored lines). Points are mean VSTM load preferences within each bin of 2 mm cortical distance. Correlations in unbinned data: *=$p$<0.05; **=$p$<0.01; ***=$p$<0.001; ****=$p$<0.0001; n.s. = no significant correlation).

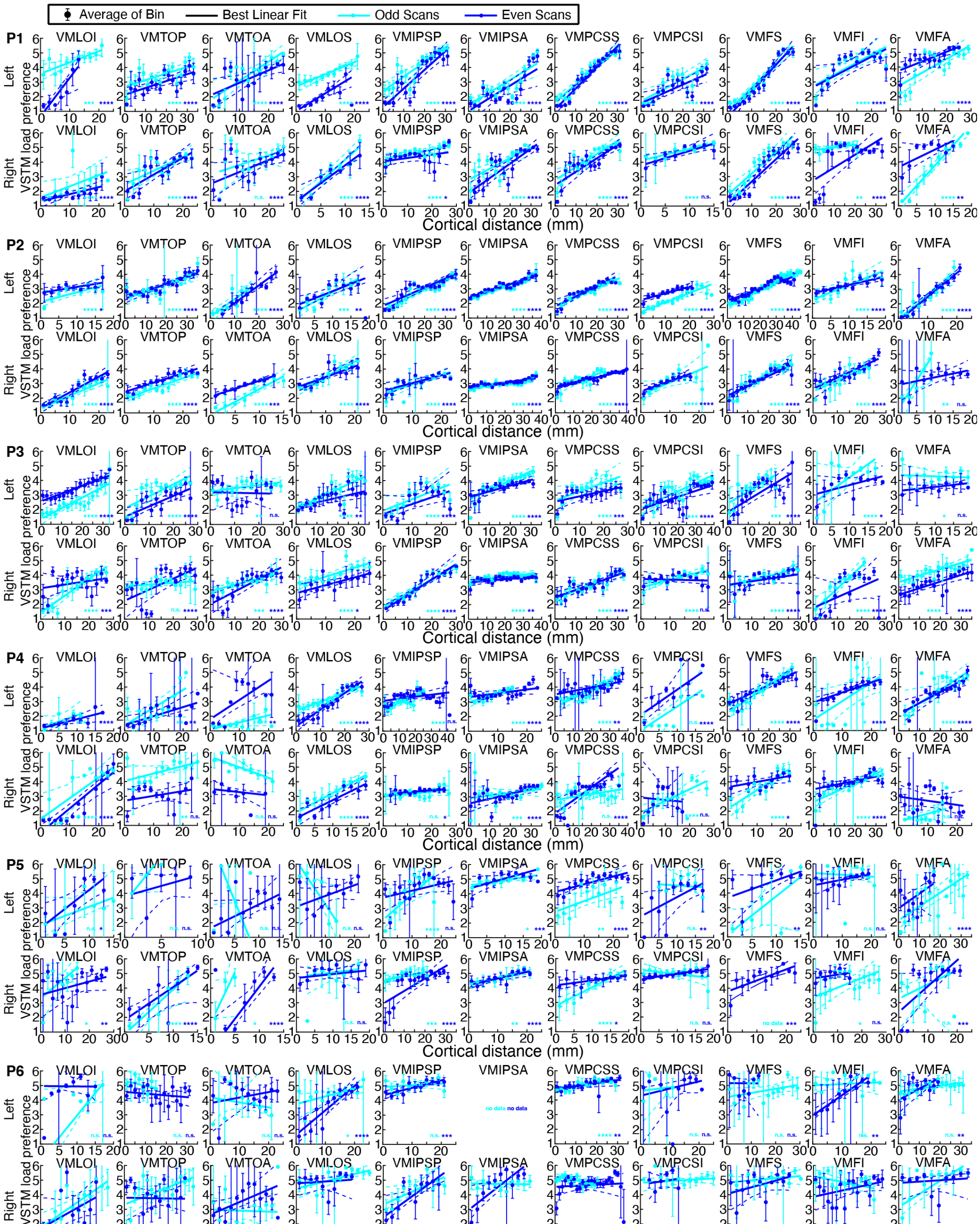


**Figure S12, VSTM load preferences from odd and even scans were significantly correlated with cortical distance between the low and high load-preferring ends of maps defined on the complementary data, related to Figure 3A.** The same progression was found across repeated measures (colored lines). Points are mean VSTM load preferences within each bin of 2 mm cortical distance. Correlations in unbinned data: *=$p$<0.05; **=$p$<0.01; ***=$p$<0.001; ****=$p$<0.0001; n.s. = no significant correlation; no data = no responsive locations in complementary map).

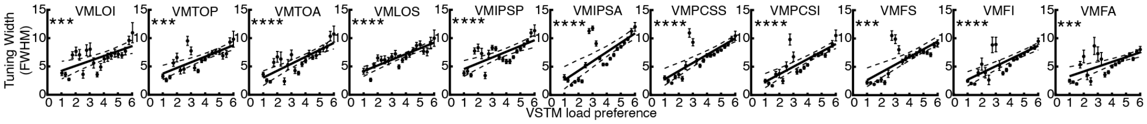


**Figure S13, Tuning width increases systematically with VSTM load preference across all maps.** Tuning width (full width at half maximum, FWHM) plotted as a function of VSTM load preference for voxels pooled across all participants and hemispheres. Black data points indicate mean tuning width within 0.25-item load preference bins; error bars represent ±1 SEM. Solid black lines represent best linear fits across bins; dashed lines indicate 95% confidence intervals. Unbinned Spearman correlations were statistically significant across all 11 maps (***$p$<0.001, ****$p$<0.0001).

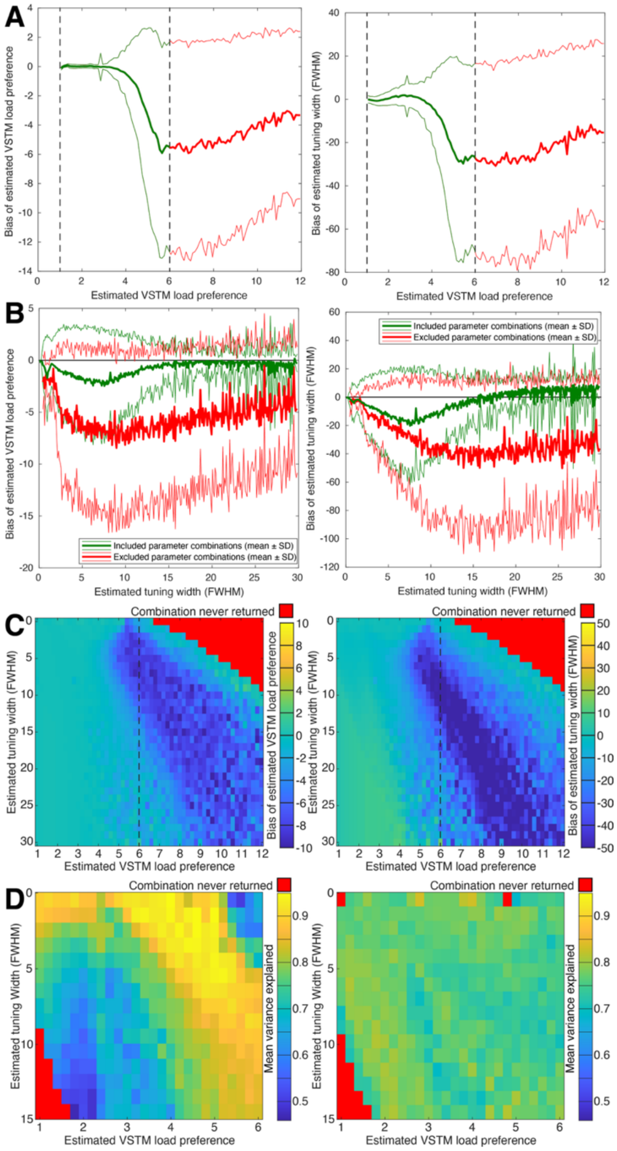


**Figure S14, Validation simulations reveal edge-of-range biases that underestimate true VSTM load preferences and tuning widths.** (**A**) Simulated fMRI responses following log-Gaussian response functions with true VSTM load preferences below 4 yield highly accurate parameter estimates. However, as true preferences increase beyond the tested range, functions fall largely outside the tested VSTM load window, leading to wider confidence intervals and a general underestimation of parameters. This underestimation occurs for both included parameters (estimated preferences between 4 and 6) and excluded parameters (estimated preferences above 6). (**B**, **C**) Due to these edge effects, simulated responses with estimated preferences above 6 (red, excluded) severely underestimate true values. Crucially, even within our analyzed range (green, estimated preferences below 6), parameters are systematically underestimated when true preferences exceed 4 or tuning widths fall below 10. Because these parameter combinations occur frequently in our fMRI data, these simulation biases demonstrate that the true experimental progressions of VSTM load preferences and tuning widths are likely steeper and stronger than our empirical models reveal. (**D**) When a constant level of noise is added, simulated fMRI responses yield higher variance explained for functions with higher VSTM load preferences (left). However, normalizing response amplitudes before adding noise eliminates this pattern (right). This indicates that the preference-dependent change in variance explained is driven by greater neural response modulations when high-preference functions interact with our VSTM load sequence.

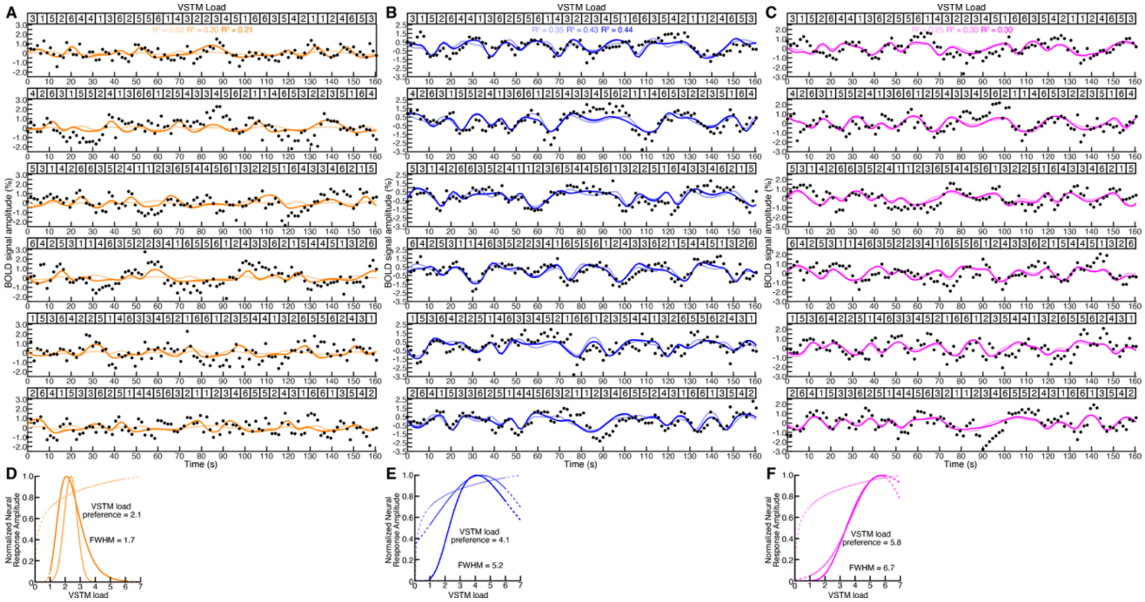


**Figure S15, Responses in the unpredictable trail order condition support VSTM load tuning.** (**A-C**) Examples of observed fMRI blood-oxygen-level-dependent (BOLD) responses (black dots) from the three example recording sites shown in Fig. 1C elicited by the sequence of changing VSTM loads in the unpredictable order control experiment (top inset). As in the main experiment, the largest response in the first recording site (A) occurs after trials with a VSTM load of 2, in the second site (B) after a VSTM load of 4, and in the second site (C) after a VSTM load of 6. We quantified these differences using the predictions of logarithm Gaussian tuned neural response models (dark colored lines), which fit the responses more closely than linear Gaussian tuned (medium colored lines) or logarithmic monotonic (light colored lines) response models. The logarithmic Gaussian VSTM load-tuned response model captured more of the variance ($R^2$) in the responses (dark text) outperforming the linear Gaussian (medium text) and logarithmic monotonic (light text) response models. (**D-F**) The best fitting neural response models for each response. The best fitting model (dark lines) describes a Gaussian tuning function of logarithmic VSTM load with two parameters: the VSTM load preference and tuning width defined by the full width at half maximum (FWHM). Different model parameters explain the response differences seen in (A-C). Linear Gaussian and logarithmic monotonic response functions are shown by medium and light lines respectively. Dashed lines extend the response functions beyond the tested VSTM loads.

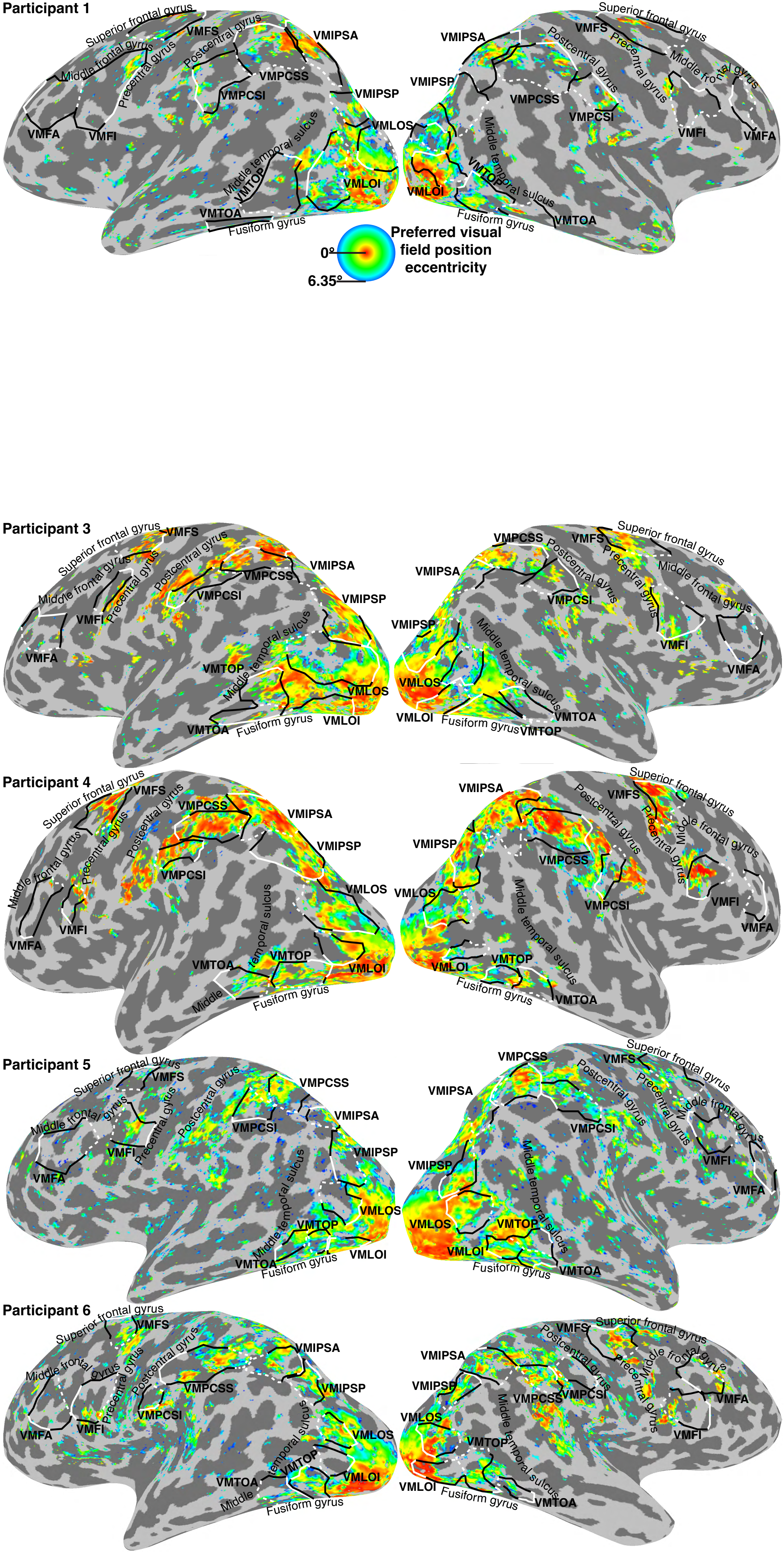


**Figure S16, Visual field eccentricity preferences from visual field mapping data, projected onto the cortical surface.** Visual field mapping data was not collected for P2. Black and white lines show the borders of VSTM load maps, for comparison to the locations and eccentricity preferences of visual spatially tuned responses.

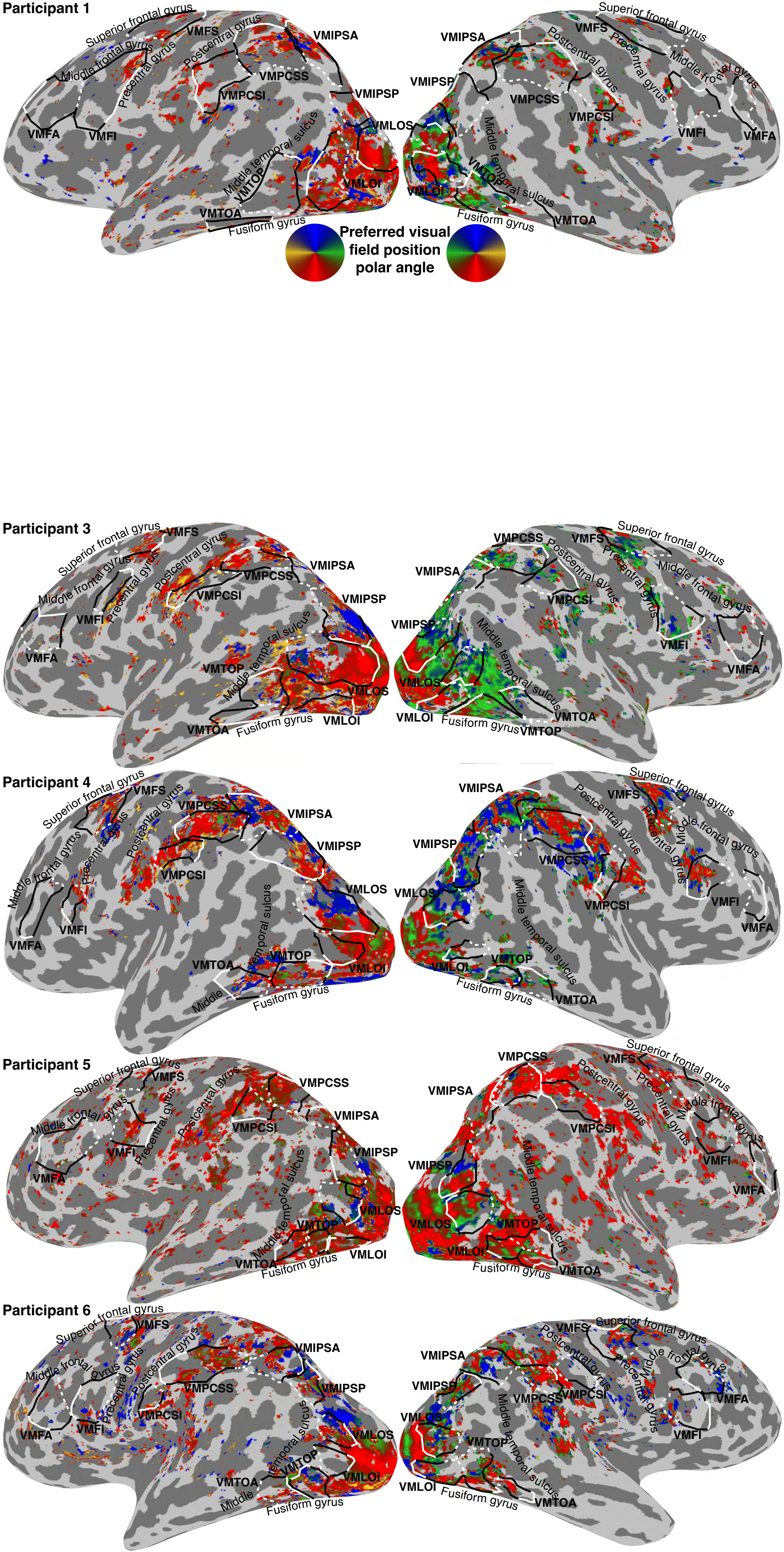


**Figure S17, Visual field polar angle preferences from visual field mapping data, projected onto the cortical surface.** Visual field mapping data was not collected for P2. Black and white lines show the borders of VSTM load maps, for comparison to the locations and polar angle preferences of visual spatially tuned responses.

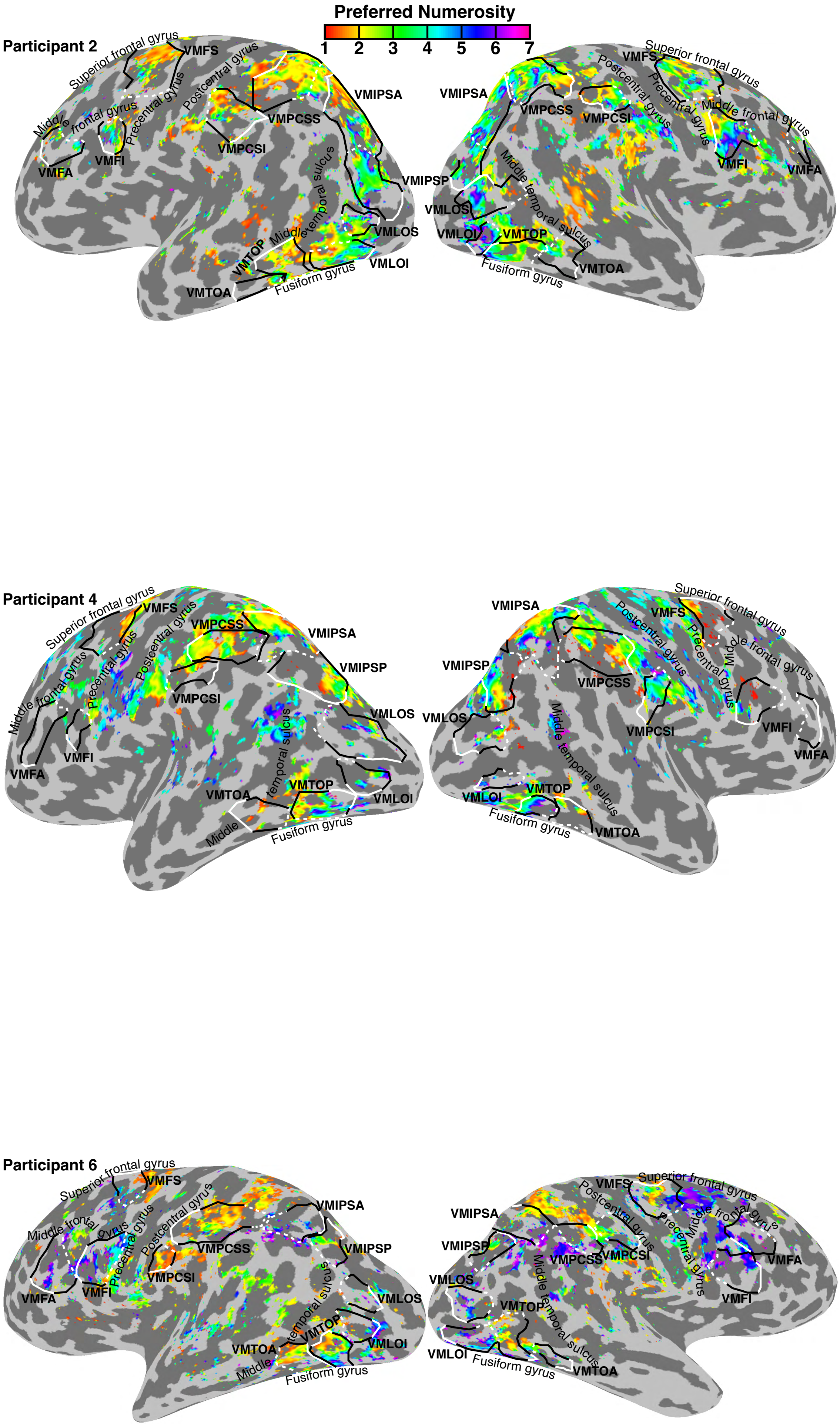


**Figure S18, Visual numerosity preferences from numerosity mapping data, projected onto the cortical surface.** Numerosity mapping data was only collected for P2, P4 and P6. Black and white lines show the borders of VSTM load maps, for comparison to the locations and numerosity preferences of numerosity tuned responses.

**Tables**

| | Surface-based transformation Mean (sd) center Talairach N27 x,y,z coordinates | | Corresponding Mean (sd) center MNI x,y,z coordinates | |
|---|---|---|---|---|
| Map | Left hemisphere | Right hemisphere | Left hemisphere | Right hemisphere |
| VMLOI | -33(5), -76(5), 3(6) | 37(5), -73(9), 1(9) | -33(5), -78(5), -2(7) | 38(5), -75(10), -3(10) |
| VMTOP | -41(5), -61(10), 2(4) | 43(2), -58(4), -3(3) | -41(5), -63(10), -1(6) | 44(2), -59(4), -7(3) |
| VMTOA | -45(2), -50(7), -4(4) | 49(4), -45(6), -5(5) | -46(2), -51(7), -8(5) | 50(4), -47(6), -8(5) |
| VMLOS | -30(4), -79(6), 17(6) | 36(10), -72(7), 14(4) | -31(4), -82(6), 14(7) | 36(10), -74(7), 11(5) |
| VMIPSP | -24(4), -67(10), 33(3) | 32(4), -69(7), 29(8) | -24(4), -70(10), 32(3) | 31(4), -72(7), 28(9) |
| VMIPSA | -18(5), -58(5), 54(3) | 18(8), -60(3), 55(5) | -18(5), -62(5), 55(4) | 18(8), -65(3), 56(5) |
| VMPCSS | -29(5), -43(7), 49(7) | 32(6), -44(7), 50(6) | -29(5), -47(8), 51(8) | 32(6), -48(8), 51(6) |
| VMPCSI | -46(2), -30(7), 40(7) | 44(7), -32(6), 44(4) | -47(2), -30(7), 40(7) | 44(7), -35(7), 46(4) |
| VMFS | -21(4), 2(6), 57(4) | 22(5), -4(6), 60(2) | -21(4), 0(6), 62(5) | 22(5), -7(6), 65(3) |
| VMFI | -39(4), 4(1), 37(6) | 39(6), 10(6), 35(5) | -39(5), 3(1), 41(7) | 40(6), 8(6), 38(5) |
| VMFA | -37(6), 36(10), 32(6) | 40(4), 34(10), 29(5) | -37(6), 36(10), 37(6) | 40(4), 33(11), 34(5) |

**Table S1, Distribution of coordinates of each map's center from each participant after transformation onto standard template brains.** Given in both Talairach (N27) coordinates and corresponding Montreal Neurological Institute (MNI) template coordinates.

| P | H | VMLOI | VMTOP | VMTOA | VMLOS | VMIPSP | VMIPSA | VMPCSS | VMPCSI | VMFS | VMFI | VMFA | p<0.05 | % |
|---|---|---|---|---|---|---|---|---|---|---|---|---|---|---|
| P1 | L | 7.2e-05 | 4.5e-49 | 8.3e-17 | 1.2e-11 | 1e-71 | 1.2e-74 | 5.8e-119 | 4e-77 | 4.9e-96 | 5.7e-94 | 1e-48 | 11 | 100 |
| P1 | R | 0.003 | 7.9e-38 | 2.2e-28 | 5.7e-19 | 3.7e-39 | 1.2e-65 | 2e-63 | 2.3e-20 | 1.5e-98 | 1.1e-35 | 1.2e-72 | 11 | 100 |
| P2 | L | 6.4e-06 | 4.3e-17 | 0.00019 | 6.3e-10 | 3.2e-50 | 3.4e-84 | 5.1e-47 | 7.3e-18 | 5.5e-37 | 3.1e-08 | 1e-08 | 11 | 100 |
| P2 | R | 3.5e-29 | 1.5e-09 | 3.6e-09 | 9.5e-14 | 1.8e-33 | 3.8e-66 | 4.9e-43 | 1.9e-15 | 1.6e-25 | 0.00013 | 0.014 | 11 | 100 |
| P3 | L | 0.019 | 1.7e-13 | 1 | 2.3e-20 | 1.5e-20 | 2.6e-13 | 1.3e-19 | 2.9e-14 | 2.9e-08 | 2e-14 | 0.024 | 10 | 91 |
| P3 | R | 0.77 | 0.14 | 0.0033 | 0.076 | 5e-35 | 1.1e-10 | 5.1e-12 | 3.6e-12 | 3e-18 | 1.1e-23 | 2.7e-07 | 8 | 73 |
| P4 | L | 0.018 | 0.002 | 0.15 | 7e-14 | 9.8e-13 | 3.3e-12 | 0.00013 | 0.068 | 0.0012 | 1 | 0.0027 | 8 | 73 |
| P4 | R | 2.6e-05 | 0.53 | 0.097 | 4.6e-11 | 0.00019 | 2.9e-08 | 1.1e-08 | 2.6e-08 | 0.0017 | 7.6e-06 | 0.92 | 8 | 73 |
| P5 | L | 0.34 | 0.046 | 0.63 | 0.0031 | 0.00019 | 5.2e-06 | 0.0012 | 0.0026 | 0.067 | 0.019 | 3.9e-05 | 8 | 73 |
| P5 | R | 9e-08 | 0.023 | 1.6e-08 | 1 | 0.00022 | 2.6e-19 | 3.3e-05 | 2e-05 | 8.5e-10 | 5.3e-08 | 0.65 | 9 | 82 |
| P6 | L | 0.94 | 0.12 | 0.043 | 0.0014 | 0.6 | 0.0055 | 1.4e-06 | 1 | 1 | 0.0035 | 0.001 | 6 | 55 |
| P6 | R | 0.45 | 1 | 2.3e-06 | 3.5e-05 | 7.6e-07 | 7.9e-10 | 0.00025 | 1 | 0.46 | 1.9e-16 | 0.00064 | 7 | 64 |
| <0.05 | | 8 | 8 | 8 | 10 | 11 | 12 | 12 | 9 | 9 | 11 | 10 | 108 | Tot |
| % | | 67 | 67 | 67 | 83 | 92 | 100 | 100 | 75 | 75 | 92 | 83 | 82 | % |

| P | H | VMLOI | VMTOP | VMTOA | VMLOS | VMIPSP | VMIPSA | VMPCSS | VMPCSI | VMFS | VMFI | VMFA |
|---|---|---|---|---|---|---|---|---|---|---|---|---|
| P1 | L | 0.576 | 0.867 | 0.907 | 0.613 | 0.914 | 0.904 | 0.941 | 0.915 | 0.937 | 0.915 | 0.842 |
| P1 | R | 0.495 | 0.913 | 0.942 | 0.899 | 0.840 | 0.863 | 0.878 | 0.876 | 0.945 | 0.799 | 0.963 |
| P2 | L | 0.547 | 0.707 | 0.694 | 0.848 | 0.842 | 0.825 | 0.716 | 0.717 | 0.695 | 0.594 | 0.621 |
| P2 | R | 0.934 | 0.654 | 0.814 | 0.765 | 0.846 | 0.793 | 0.696 | 0.778 | 0.735 | 0.583 | 0.601 |
| P3 | L | 0.255 | 0.716 | -0.129 | 0.811 | 0.836 | 0.530 | 0.735 | 0.771 | 0.635 | 0.901 | 0.337 |
| P3 | R | 0.046 | 0.204 | 0.354 | 0.276 | 0.794 | 0.379 | 0.542 | 0.609 | 0.718 | 0.791 | 0.564 |
| P4 | L | 0.684 | 0.915 | 0.541 | 0.798 | 0.571 | 0.556 | 0.576 | 0.740 | 0.541 | -0.071 | 0.512 |
| P4 | R | 0.895 | 0.149 | 0.303 | 0.854 | 0.377 | 0.472 | 0.697 | 0.900 | 0.564 | 0.626 | 0.055 |
| P5 | L | 0.328 | 0.899 | 0.611 | 0.706 | 0.654 | 0.497 | 0.458 | 0.602 | 0.504 | 0.740 | 0.800 |
| P5 | R | 0.806 | 0.620 | 0.847 | -0.057 | 0.472 | 0.705 | 0.434 | 0.551 | 0.704 | 0.682 | 0.091 |
| P6 | L | 0.065 | 0.427 | 0.499 | 0.513 | 0.103 | 0.362 | 0.902 | -0.022 | -0.217 | 0.665 | 0.504 |
| P6 | R | 0.227 | -0.295 | 0.816 | 0.647 | 0.718 | 0.548 | 0.745 | -0.122 | 0.169 | 0.843 | 0.662 |

| P | H | VMLOI | VMTOP | VMTOA | VMLOS | VMIPSP | VMIPSA | VMPCSS | VMPCSI | VMFS | VMFI | VMFA |
|---|---|---|---|---|---|---|---|---|---|---|---|---|
| P1 | L | 44 | 161 | 44 | 104 | 183 | 203 | 255 | 196 | 211 | 240 | 180 |
| P1 | R | 36 | 97 | 60 | 52 | 146 | 220 | 198 | 63 | 205 | 159 | 128 |
| P2 | L | 63 | 108 | 25 | 34 | 186 | 340 | 297 | 109 | 253 | 76 | 73 |
| P2 | R | 65 | 71 | 36 | 68 | 121 | 306 | 294 | 73 | 147 | 40 | 17 |
| P3 | L | 91 | 81 | 41 | 85 | 77 | 171 | 112 | 69 | 65 | 39 | 48 |
| P3 | R | 64 | 58 | 71 | 46 | 159 | 282 | 145 | 111 | 111 | 108 | 75 |
| P4 | L | 12 | 8 | 9 | 60 | 137 | 139 | 41 | 7 | 35 | 7 | 34 |
| P4 | R | 14 | 25 | 34 | 37 | 99 | 130 | 54 | 22 | 30 | 45 | 13 |
| P5 | L | 12 | 5 | 3 | 16 | 29 | 80 | 50 | 24 | 15 | 10 | 20 |
| P5 | R | 31 | 14 | 29 | 7 | 60 | 124 | 90 | 56 | 60 | 52 | 35 |
| P6 | L | 7 | 16 | 18 | 38 | 37 | 61 | 17 | 5 | 9 | 18 | 42 |
| P6 | R | 16 | 8 | 24 | 36 | 38 | 113 | 20 | 9 | 26 | 59 | 24 |

**Table S2, Descriptive statistics for correlations between VSTM load preferences estimated from odd and even numbered scan runs.** Top: *p*-values of correlations in each map for each hemisphere, after correction for false discovery rate and upsampling between the native fMRI scan resolution and the participant's cortical surface model. Middle: Pearson's correlation coefficient (*r*) in each map. Bottom: Number of selected measurement in each map after correction for upsampling. Selected measurements must have >20% variance explained by the response model in both sets of scanning runs.

| P | H | VMLOI | VMTOP | VMTOA | VMLOS | VMIPSP | VMIPSA | VMPCSS | VMPCSI | VMFS | VMFI | VMFA | p<0.05 | % |
|---|---|---|---|---|---|---|---|---|---|---|---|---|---|---|
| P1 | L | 7e-09 | 7.2e-29 | 1.8e-14 | 3.2e-10 | 9.7e-66 | 2.3e-53 | 1.7e-93 | 3.8e-54 | 1.4e-79 | 1.1e-104 | 1e-43 | 11 | 100 |
| P1 | R | 0.0067 | 8.7e-19 | 6.7e-13 | 2.5e-09 | 6.3e-33 | 1.7e-44 | 2.3e-45 | 7.8e-18 | 8.2e-80 | 1.8e-32 | 1.5e-49 | 11 | 100 |
| P2 | L | 7.7e-05 | 5.1e-11 | 3.8e-07 | 2.3e-08 | 5.8e-49 | 2.4e-64 | 3.8e-44 | 5e-13 | 3.5e-44 | 3.9e-08 | 2.9e-08 | 11 | 100 |
| P2 | R | 6.8e-28 | 7.5e-10 | 0.0017 | 2.3e-07 | 1.1e-17 | 2.6e-44 | 5.8e-22 | 0.094 | 2.4e-10 | 5.4e-05 | 0.022 | 10 | 91 |
| P3 | L | 0.27 | 4.2e-05 | 1 | 6e-11 | 1.7e-08 | 2.4e-05 | 1.6e-07 | 2.9e-09 | 1.3e-06 | 1.6e-10 | 0.63 | 8 | 73 |
| P3 | R | 1 | 1 | 0.00056 | 0.081 | 1.4e-14 | 1.8e-06 | 4.2e-05 | 5.6e-05 | 2.1e-06 | 1e-11 | 0.036 | 8 | 73 |
| P4 | L | 0.032 | 0.025 | 0.15 | 5e-12 | 3.4e-07 | 3e-17 | 4.7e-06 | 0.54 | 0.005 | 0.89 | 0.032 | 8 | 73 |
| P4 | R | 1.5e-07 | 0.99 | 0.76 | 3.3e-06 | 0.0018 | 8.8e-07 | 1e-12 | 0.005 | 3.9e-05 | 3.5e-07 | 0.13 | 8 | 73 |
| P5 | L | 0.13 | 0.48 | 0.017 | 0.52 | 0.0058 | 0.0035 | 0.00019 | 1.1e-05 | 0.053 | 0.00015 | 1.6e-05 | 7 | 64 |
| P5 | R | 2.4e-05 | 0.061 | 0.00019 | 1 | 0.00055 | 1.4e-19 | 3.3e-07 | 9.7e-07 | 0.00032 | 9e-10 | 0.43 | 8 | 73 |
| P6 | L | 1 | 0.89 | 0.0051 | 0.00074 | 0.2 | 0.098 | 4.6e-09 | 0.82 | 1 | 2.7e-09 | 3.3e-08 | 5 | 45 |
| P6 | R | 0.12 | 0.6 | 1.6e-08 | 4.4e-07 | 2.6e-06 | 1.1e-14 | 8.5e-05 | 0.025 | 0.12 | 1.5e-14 | 1.1e-05 | 8 | 73 |
| <0.05 | | 7 | 6 | 9 | 9 | 11 | 11 | 12 | 9 | 9 | 11 | 9 | 103 | Tot |
| % | | 58 | 50 | 75 | 75 | 92 | 92 | 100 | 75 | 75 | 92 | 75 | 78 | % |

| P | H | VMLOI | VMTOP | VMTOA | VMLOS | VMIPSP | VMIPSA | VMPCSS | VMPCSI | VMFS | VMFI | VMFA |
|---|---|---|---|---|---|---|---|---|---|---|---|---|
| P1 | L | 0.742 | 0.745 | 0.840 | 0.591 | 0.892 | 0.834 | 0.909 | 0.848 | 0.915 | 0.927 | 0.819 |
| P1 | R | 0.417 | 0.810 | 0.821 | 0.773 | 0.814 | 0.782 | 0.814 | 0.874 | 0.914 | 0.778 | 0.912 |
| P2 | L | 0.493 | 0.602 | 0.819 | 0.792 | 0.842 | 0.763 | 0.703 | 0.664 | 0.734 | 0.597 | 0.615 |
| P2 | R | 0.928 | 0.677 | 0.496 | 0.605 | 0.684 | 0.693 | 0.520 | 0.197 | 0.509 | 0.621 | 0.518 |
| P3 | L | 0.126 | 0.466 | -0.012 | 0.650 | 0.600 | 0.336 | 0.522 | 0.655 | 0.589 | 0.883 | 0.083 |
| P3 | R | -0.089 | -0.116 | 0.401 | 0.311 | 0.575 | 0.299 | 0.355 | 0.399 | 0.470 | 0.625 | 0.243 |
| P4 | L | 0.643 | 0.754 | 0.547 | 0.760 | 0.437 | 0.658 | 0.662 | 0.725 | 0.480 | 0.069 | 0.391 |
| P4 | R | 0.931 | 0.016 | 0.071 | 0.711 | 0.322 | 0.433 | 0.813 | 0.572 | 0.702 | 0.690 | 0.416 |
| P5 | L | 0.487 | 0.362 | 0.906 | 0.212 | 0.438 | 0.379 | 0.532 | 0.726 | 0.549 | 0.893 | 0.810 |
| P5 | R | 0.733 | 0.757 | 0.655 | -0.039 | 0.449 | 0.719 | 0.548 | 0.619 | 0.531 | 0.763 | 0.143 |
| P6 | L | -0.179 | 0.055 | 0.619 | 0.520 | 0.252 | 0.229 | 0.864 | 0.084 | -0.095 | 0.908 | 0.713 |
| P6 | R | 0.387 | 0.217 | 0.882 | 0.758 | 0.701 | 0.679 | 0.779 | 0.484 | 0.311 | 0.812 | 0.784 |

| P | H | VMLOI | VMTOP | VMTOA | VMLOS | VMIPSP | VMIPSA | VMPCSS | VMPCSI | VMFS | VMFI | VMFA |
|---|---|---|---|---|---|---|---|---|---|---|---|---|
| P1 | L | 47 | 161 | 53 | 100 | 191 | 206 | 247 | 196 | 203 | 248 | 179 |
| P1 | R | 44 | 79 | 51 | 44 | 138 | 215 | 191 | 56 | 205 | 158 | 128 |
| P2 | L | 62 | 104 | 27 | 36 | 182 | 338 | 294 | 97 | 259 | 75 | 71 |
| P2 | R | 65 | 68 | 40 | 65 | 124 | 308 | 308 | 82 | 144 | 38 | 21 |
| P3 | L | 91 | 75 | 42 | 85 | 78 | 161 | 94 | 69 | 61 | 31 | 47 |
| P3 | R | 76 | 86 | 75 | 36 | 158 | 261 | 135 | 102 | 98 | 101 | 82 |
| P4 | L | 12 | 9 | 9 | 61 | 133 | 134 | 41 | 3 | 35 | 12 | 33 |
| P4 | R | 17 | 22 | 32 | 35 | 98 | 126 | 52 | 24 | 29 | 45 | 16 |
| P5 | L | 12 | 7 | 6 | 14 | 41 | 62 | 47 | 30 | 14 | 12 | 21 |
| P5 | R | 27 | 7 | 29 | 7 | 59 | 120 | 80 | 55 | 44 | 48 | 40 |
| P6 | L | 10 | 16 | 20 | 41 | 31 | 59 | 29 | 15 | 9 | 24 | 48 |
| P6 | R | 19 | 10 | 25 | 34 | 37 | 104 | 20 | 23 | 29 | 60 | 24 |

**Table S3, Descriptive statistics for correlations between VSTM load preferences estimated from constant performance and constant orientation conditions.** Top: *p*-values of correlations in each map for each hemisphere, after correction for false discovery rate and upsampling between the native fMRI scan resolution and the participant's cortical surface model. Middle: Pearson's correlation coefficient (*r*) in each map. Bottom: Number of selected measurement in each map after correction for upsampling. Selected measurements must have >20% variance explained by the response model in both conditions.

| P | H | VMLOI | VMTOP | VMTOA | VMLOS | VMIPSP | VMIPSA | VMPCSS | VMPCSI | VMFS | VMFI | VMFA | p<0.05 | % |
|---|---|---|---|---|---|---|---|---|---|---|---|---|---|---|
| P1 | L | 1.5e-10 | 1.3e-05 | 1.4e-10 | 2.4e-19 | 5.1e-22 | 1e-27 | 5.7e-72 | 6.9e-31 | 1.9e-59 | 1.1e-14 | 5.7e-08 | 11 | 100 |
| P1 | R | 1.5e-05 | 1.5e-07 | 0.0006 | 9.7e-13 | 0.072 | 6.8e-27 | 1.9e-29 | 3.8e-06 | 5.4e-40 | 4.4e-08 | 1.4e-28 | 10 | 91 |
| P2 | L | 8e-06 | 5.6e-17 | 4.3e-09 | 8.3e-06 | 9.7e-34 | 7.4e-40 | 9e-43 | 1.4e-16 | 1.3e-18 | 5.6e-08 | 2.8e-25 | 11 | 100 |
| P2 | R | 1.3e-19 | 7e-18 | 2.5e-12 | 2.1e-14 | 5.6e-16 | 9.8e-15 | 1.6e-13 | 2.5e-07 | 5e-21 | 2e-07 | 1.1e-05 | 11 | 100 |
| P3 | L | 1.9e-12 | 2.4e-21 | 4.1e-05 | 1.7e-06 | 1.7e-11 | 6.4e-12 | 3.5e-12 | 1.4e-15 | 1.3e-18 | 4.7e-12 | 1.5e-06 | 11 | 100 |
| P3 | R | 2.8e-19 | 1.7e-09 | 2.1e-14 | 0.0035 | 1.7e-43 | 1.8e-07 | 9e-11 | 2.1e-05 | 1.2e-09 | 8.4e-22 | 5.9e-07 | 11 | 100 |
| P4 | L | 2.3e-09 | 0.0008 | 3.8e-14 | 1.8e-22 | 1.1e-14 | 1.6e-14 | 7.4e-09 | 2.5e-13 | 2.9e-15 | 1.1e-06 | 5.9e-09 | 11 | 100 |
| P4 | R | 2.7e-22 | 7.4e-06 | 0.032 | 4.6e-10 | 4.5e-05 | 1e-08 | 4.1e-13 | 9.6e-06 | 1.4e-11 | 1.9e-09 | 1e-05 | 11 | 100 |
| P5 | L | 3.8e-05 | 0.2 | 9.7e-06 | 0.055 | 6.1e-15 | 0.00011 | 1.3e-06 | 0.56 | 0.00013 | 1.4e-06 | 0.00081 | 8 | 73 |
| P5 | R | 0.00096 | 7.6e-05 | 6.4e-12 | 0.5 | 0.00011 | 0.00085 | 6.5e-07 | 0.025 | 1.3e-05 | 0.004 | 0.00026 | 10 | 91 |
| P6 | L | 0.0059 | 2.2e-06 | 0.054 | 4.6e-09 | 0.00025 | 0.051 | 1 | 0.045 | 1 | 0.00017 | 3e-05 | 7 | 64 |
| P6 | R | 5.6e-17 | 0.054 | 4.8e-06 | 4.7e-07 | 0.00015 | 1.7e-05 | 0.76 | 0.032 | 0.045 | 0.00048 | 1.3e-07 | 9 | 82 |
| <0.05 | | 12 | 10 | 11 | 10 | 11 | 11 | 10 | 11 | 11 | 12 | 12 | 121 | Tot |
| % | | 100 | 83 | 92 | 83 | 92 | 92 | 83 | 92 | 92 | 100 | 100 | 92 | % |

| P | H | VMLOI | VMTOP | VMTOA | VMLOS | VMIPSP | VMIPSA | VMPCSS | VMPCSI | VMFS | VMFI | VMFA |
|---|---|---|---|---|---|---|---|---|---|---|---|---|
| P1 | L | 0.648 | 0.288 | 0.672 | 0.671 | 0.568 | 0.628 | 0.811 | 0.660 | 0.791 | 0.425 | 0.328 |
| P1 | R | 0.460 | 0.420 | 0.336 | 0.707 | 0.131 | 0.578 | 0.627 | 0.472 | 0.688 | 0.379 | 0.723 |
| P2 | L | 0.398 | 0.576 | 0.678 | 0.504 | 0.681 | 0.583 | 0.625 | 0.618 | 0.457 | 0.490 | 0.767 |
| P2 | R | 0.722 | 0.623 | 0.689 | 0.658 | 0.580 | 0.376 | 0.363 | 0.411 | 0.584 | 0.559 | 0.559 |
| P3 | L | 0.524 | 0.670 | 0.410 | 0.352 | 0.537 | 0.411 | 0.511 | 0.631 | 0.648 | 0.741 | 0.434 |
| P3 | R | 0.715 | 0.463 | 0.638 | 0.276 | 0.713 | 0.263 | 0.394 | 0.330 | 0.427 | 0.623 | 0.365 |
| P4 | L | 0.759 | 0.493 | 0.832 | 0.762 | 0.475 | 0.514 | 0.593 | 0.832 | 0.767 | 0.771 | 0.635 |
| P4 | R | 0.926 | 0.517 | 0.261 | 0.661 | 0.309 | 0.366 | 0.574 | 0.495 | 0.662 | 0.600 | 0.479 |
| P5 | L | 0.551 | 0.298 | 0.841 | 0.264 | 0.680 | 0.346 | 0.459 | 0.080 | 0.521 | 0.613 | 0.355 |
| P5 | R | 0.408 | 0.592 | 0.730 | 0.134 | 0.411 | 0.243 | 0.411 | 0.250 | 0.325 | 0.274 | 0.346 |
| P6 | L | 0.469 | 0.566 | 0.189 | 0.597 | 0.434 | 0.201 | -0.001 | 0.316 | -0.013 | 0.413 | 0.370 |
| P6 | R | 0.801 | 0.258 | 0.554 | 0.514 | 0.442 | 0.300 | 0.040 | 0.256 | 0.199 | 0.299 | 0.569 |

| P | H | VMLOI | VMTOP | VMTOA | VMLOS | VMIPSP | VMIPSA | VMPCSS | VMPCSI | VMFS | VMFI | VMFA |
|---|---|---|---|---|---|---|---|---|---|---|---|---|
| P1 | L | 81 | 230 | 74 | 144 | 252 | 250 | 312 | 246 | 280 | 315 | 273 |
| P1 | R | 84 | 151 | 104 | 79 | 194 | 298 | 267 | 91 | 286 | 204 | 176 |
| P2 | L | 123 | 184 | 61 | 73 | 247 | 438 | 396 | 151 | 349 | 115 | 129 |
| P2 | R | 119 | 160 | 82 | 110 | 170 | 414 | 405 | 153 | 224 | 78 | 56 |
| P3 | L | 164 | 160 | 97 | 183 | 141 | 268 | 169 | 134 | 151 | 65 | 118 |
| P3 | R | 120 | 160 | 119 | 114 | 282 | 400 | 262 | 165 | 195 | 198 | 185 |
| P4 | L | 46 | 44 | 53 | 117 | 247 | 203 | 83 | 50 | 76 | 30 | 71 |
| P4 | R | 53 | 70 | 70 | 73 | 174 | 240 | 140 | 75 | 85 | 87 | 80 |
| P5 | L | 51 | 21 | 19 | 55 | 105 | 123 | 106 | 59 | 50 | 54 | 88 |
| P5 | R | 64 | 40 | 67 | 30 | 86 | 191 | 142 | 84 | 179 | 112 | 110 |
| P6 | L | 34 | 63 | 107 | 84 | 69 | 98 | 102 | 42 | 33 | 80 | 125 |
| P6 | R | 74 | 58 | 62 | 89 | 71 | 206 | 66 | 73 | 106 | 137 | 77 |

**Table S4, Descriptive statistics for correlations between VSTM load preferences estimated from all data and the cortical surface distance across the map, related to Figure 3A.** Top: *p*-values of correlations in each map for each hemisphere, after correction for false discovery rate and upsampling between the native fMRI scan resolution and the participant's cortical surface model. Middle: Pearson's correlation coefficient (*r*) in each map. Bottom: Number of selected measurement in each map after correction for upsampling. Selected measurements must have >20% variance explained by the response model fit to these data.

| P | H | VMLOI | VMTOP | VMTOA | VMLOS | VMIPSP | VMIPSA | VMPCSS | VMPCSI | VMFS | VMFI | VMFA | p<0.05 | % |
|---|---|---|---|---|---|---|---|---|---|---|---|---|---|---|
| P1 | L | 1.8e-05 | 0.0076 | 3.2e-06 | 8.4e-13 | 3.9e-22 | 3.4e-23 | 6.9e-53 | 4.4e-32 | 4.4e-50 | 7.2e-11 | 1.4e-08 | 11 | 100 |
| P1 | R | 0.048 | 2.3e-05 | 0.016 | 2.1e-07 | 0.14 | 1.3e-24 | 1.2e-28 | 0.0016 | 4.3e-44 | 2.3e-05 | 5.6e-29 | 10 | 91 |
| P2 | L | 3.3e-05 | 4.4e-22 | 2.8e-07 | 0.00081 | 1.7e-27 | 1.5e-32 | 1.3e-35 | 7.8e-21 | 3.3e-24 | 5.2e-09 | 6e-18 | 11 | 100 |
| P2 | R | 2.2e-12 | 4.9e-12 | 2e-10 | 2.8e-09 | 6.3e-10 | 1.3e-12 | 3.5e-20 | 3.4e-29 | 2.7e-19 | 8.3e-05 | 0.0017 | 11 | 100 |
| P3 | L | 0.025 | 4e-06 | 9.5e-05 | 2.3e-05 | 1.7e-06 | 7e-11 | 0.00089 | 1.3e-14 | 7e-11 | 4.1e-07 | 0.0013 | 11 | 100 |
| P3 | R | 2.3e-05 | 0.94 | 3.7e-15 | 0.15 | 6.3e-21 | 1.9e-05 | 0.0006 | 0.0089 | 1.3e-06 | 5.3e-14 | 8.8e-08 | 9 | 82 |
| P4 | L | 0.006 | 0.0043 | 2e-05 | 1.2e-16 | 1.4e-14 | 1e-09 | 7e-06 | 0.29 | 2.7e-07 | 0.00048 | 0.45 | 9 | 82 |
| P4 | R | 6.8e-20 | 0.15 | 0.04 | 1.4e-06 | 1.6e-08 | 0.004 | 1.2e-06 | 0.5 | 5.3e-05 | 2.2e-07 | 0.062 | 8 | 73 |
| P5 | L | 0.045 | 0.44 | 0.0011 | 0.15 | 1.8e-08 | 0.0013 | 1.2e-05 | 1 | 0.014 | 2e-06 | 0.0089 | 8 | 73 |
| P5 | R | 0.014 | 0.00012 | 3.4e-06 | 0.13 | 0.0021 | 0.00018 | 3.1e-05 | 1 | 0.17 | 0.045 | 4.5e-05 | 8 | 73 |
| P6 | L | 0.33 | 0.016 | 0.38 | 0.00081 | 0.0074 | 1.6e-06 | 1 | 0.033 | 0.15 | 0.00017 | 2.8e-09 | 7 | 64 |
| P6 | R | 1.3e-09 | 0.0014 | 0.0012 | 0.0064 | 0.00025 | 0.0042 | 1 | 0.15 | 0.013 | 0.69 | 1.1e-09 | 8 | 73 |
| <0.05 | | 11 | 9 | 11 | 9 | 11 | 12 | 10 | 7 | 10 | 11 | 10 | 111 | Tot |
| % | | 92 | 75 | 92 | 75 | 92 | 100 | 83 | 58 | 83 | 92 | 83 | 84 | % |

| P | H | VMLOI | VMTOP | VMTOA | VMLOS | VMIPSP | VMIPSA | VMPCSS | VMPCSI | VMFS | VMFI | VMFA |
|---|---|---|---|---|---|---|---|---|---|---|---|---|
| P1 | L | 0.502 | 0.194 | 0.558 | 0.606 | 0.583 | 0.596 | 0.760 | 0.690 | 0.771 | 0.386 | 0.353 |
| P1 | R | 0.249 | 0.421 | 0.277 | 0.638 | 0.119 | 0.590 | 0.672 | 0.383 | 0.738 | 0.312 | 0.733 |
| P2 | L | 0.445 | 0.696 | 0.670 | 0.470 | 0.669 | 0.553 | 0.606 | 0.689 | 0.537 | 0.549 | 0.712 |
| P2 | R | 0.682 | 0.634 | 0.660 | 0.586 | 0.498 | 0.355 | 0.453 | 0.773 | 0.602 | 0.476 | 0.464 |
| P3 | L | 0.194 | 0.434 | 0.452 | 0.367 | 0.432 | 0.458 | 0.299 | 0.679 | 0.602 | 0.756 | 0.331 |
| P3 | R | 0.414 | 0.010 | 0.728 | 0.186 | 0.585 | 0.258 | 0.248 | 0.253 | 0.424 | 0.576 | 0.461 |
| P4 | L | 0.442 | 0.590 | 0.697 | 0.755 | 0.554 | 0.491 | 0.591 | 0.417 | 0.642 | 0.700 | 0.110 |
| P4 | R | 0.923 | 0.219 | 0.316 | 0.635 | 0.508 | 0.238 | 0.547 | 0.122 | 0.586 | 0.604 | 0.284 |
| P5 | L | 0.286 | 0.237 | 0.756 | 0.232 | 0.561 | 0.344 | 0.467 | -0.162 | 0.400 | 0.690 | 0.341 |
| P5 | R | 0.409 | 0.614 | 0.642 | 0.359 | 0.368 | 0.306 | 0.397 | -0.036 | 0.140 | 0.220 | 0.400 |
| P6 | L | 0.145 | 0.277 | 0.097 | 0.400 | 0.329 | 0.461 | -0.093 | 0.365 | 0.229 | 0.406 | 0.499 |
| P6 | R | 0.668 | 0.462 | 0.434 | 0.339 | 0.445 | 0.206 | -0.091 | 0.217 | 0.238 | 0.040 | 0.612 |

| P | H | VMLOI | VMTOP | VMTOA | VMLOS | VMIPSP | VMIPSA | VMPCSS | VMPCSI | VMFS | VMFI | VMFA |
|---|---|---|---|---|---|---|---|---|---|---|---|---|
| P1 | L | 71 | 203 | 65 | 121 | 239 | 237 | 286 | 227 | 257 | 283 | 262 |
| P1 | R | 68 | 101 | 81 | 58 | 169 | 259 | 217 | 70 | 259 | 190 | 172 |
| P2 | L | 86 | 150 | 50 | 51 | 210 | 408 | 357 | 145 | 319 | 104 | 113 |
| P2 | R | 86 | 101 | 78 | 92 | 146 | 399 | 388 | 147 | 190 | 67 | 46 |
| P3 | L | 143 | 112 | 74 | 135 | 122 | 195 | 130 | 104 | 103 | 35 | 98 |
| P3 | R | 105 | 120 | 89 | 68 | 223 | 290 | 203 | 114 | 130 | 151 | 131 |
| P4 | L | 40 | 23 | 32 | 88 | 173 | 147 | 53 | 9 | 56 | 22 | 55 |
| P4 | R | 48 | 48 | 46 | 51 | 117 | 156 | 74 | 36 | 44 | 66 | 47 |
| P5 | L | 53 | 14 | 16 | 43 | 92 | 90 | 86 | 48 | 40 | 40 | 62 |
| P5 | R | 38 | 36 | 46 | 21 | 72 | 156 | 111 | 62 | 105 | 90 | 105 |
| P6 | L | 53 | 80 | 93 | 72 | 70 | 106 | 96 | 37 | 43 | 87 | 134 |
| P6 | R | 69 | 48 | 57 | 68 | 68 | 206 | 58 | 50 | 116 | 125 | 87 |

**Table S5, Descriptive statistics for correlations between VSTM load preferences estimated from the constant performance condition and the cortical surface distance across the map, related to Figure 3A.** Top: *p*-values of correlations in each map for each hemisphere, after correction for false discovery rate and upsampling between the native fMRI scan resolution and the participant's cortical surface model. Middle: Pearson's correlation coefficient (*r*) in each map. Bottom: Number of selected measurement in each map after correction for upsampling. Selected measurements must have >20% variance explained.

| P | H | VMLOI | VMTOP | VMTOA | VMLOS | VMIPSP | VMIPSA | VMPCSS | VMPCSI | VMFS | VMFI | VMFA | p<0.05 | % |
|---|---|---|---|---|---|---|---|---|---|---|---|---|---|---|
| P1 | L | 5.2e-09 | 1.5e-07 | 1.6e-12 | 2.9e-13 | 4.7e-21 | 7.7e-24 | 1.5e-67 | 4.9e-22 | 5e-49 | 1.7e-16 | 2.1e-05 | 11 | 100 |
| P1 | R | 2.5e-07 | 3.5e-10 | 0.038 | 1.7e-10 | 0.036 | 1.2e-19 | 2.9e-21 | 1.6e-06 | 1.9e-30 | 4.6e-06 | 7.7e-24 | 11 | 100 |
| P2 | L | 0.021 | 7.1e-12 | 1e-07 | 0.00094 | 1.4e-25 | 7.9e-32 | 1.4e-24 | 3.2e-05 | 1.4e-11 | 2e-06 | 5.9e-15 | 11 | 100 |
| P2 | R | 5.6e-18 | 8.2e-15 | 3.3e-05 | 3.9e-08 | 1.7e-11 | 6.2e-08 | 0.00084 | 1 | 1.2e-05 | 3e-05 | 0.053 | 9 | 82 |
| P3 | L | 4.6e-07 | 3.9e-09 | 0.067 | 8.6e-06 | 8.6e-07 | 1.3e-07 | 1.9e-06 | 2.1e-10 | 2.1e-05 | 1.7e-10 | 0.019 | 10 | 91 |
| P3 | R | 0.013 | 2e-07 | 7.4e-05 | 6.4e-06 | 5.7e-28 | 0.073 | 2.9e-08 | 0.0018 | 4.9e-06 | 1.7e-16 | 0.082 | 9 | 82 |
| P4 | L | 0.2 | 0.036 | 1.4e-08 | 3.8e-17 | 3.6e-09 | 1.6e-12 | 5.4e-06 | 1.2e-13 | 2.5e-07 | 0.007 | 1.3e-12 | 10 | 91 |
| P4 | R | 4.5e-12 | 0.59 | 0.032 | 3.1e-08 | 0.035 | 6.1e-12 | 6.9e-11 | 6.1e-10 | 1.9e-14 | 2.3e-07 | 1.3e-05 | 10 | 91 |
| P5 | L | 0.013 | 0.2 | 0.00038 | 0.89 | 1.4e-05 | 0.00039 | 1.6e-06 | 0.12 | 9.5e-06 | 2.5e-05 | 0.01 | 8 | 73 |
| P5 | R | 0.00054 | 0.00028 | 1.2e-09 | 0.44 | 0.036 | 0.0045 | 0.015 | 0.063 | 5.3e-09 | 0.036 | 0.013 | 9 | 82 |
| P6 | L | 0.064 | 1.2e-09 | 0.041 | 1.5e-09 | 0.073 | 0.017 | 0.9 | 0.39 | 0.87 | 0.0037 | 0.013 | 6 | 55 |
| P6 | R | 6.5e-11 | 1 | 0.0041 | 8.9e-06 | 0.0006 | 5e-05 | 0.69 | 0.42 | 0.01 | 5.7e-06 | 1.3e-06 | 8 | 73 |
| <0.05 | | 10 | 9 | 11 | 10 | 11 | 11 | 10 | 7 | 11 | 12 | 10 | 112 | Tot |
| % | | 83 | 75 | 92 | 83 | 92 | 92 | 83 | 58 | 92 | 100 | 83 | 85 | % |

| P | H | VMLOI | VMTOP | VMTOA | VMLOS | VMIPSP | VMIPSA | VMPCSS | VMPCSI | VMFS | VMFI | VMFA |
|---|---|---|---|---|---|---|---|---|---|---|---|---|
| P1 | L | 0.588 | 0.361 | 0.731 | 0.536 | 0.560 | 0.603 | 0.805 | 0.596 | 0.762 | 0.456 | 0.273 |
| P1 | R | 0.513 | 0.488 | 0.214 | 0.637 | 0.157 | 0.507 | 0.542 | 0.485 | 0.636 | 0.330 | 0.697 |
| P2 | L | 0.245 | 0.546 | 0.750 | 0.454 | 0.649 | 0.580 | 0.537 | 0.408 | 0.394 | 0.492 | 0.702 |
| P2 | R | 0.755 | 0.642 | 0.554 | 0.589 | 0.539 | 0.300 | 0.188 | -0.069 | 0.325 | 0.524 | 0.368 |
| P3 | L | 0.397 | 0.468 | 0.191 | 0.338 | 0.411 | 0.339 | 0.374 | 0.551 | 0.372 | 0.712 | 0.233 |
| P3 | R | 0.217 | 0.372 | 0.355 | 0.406 | 0.632 | 0.091 | 0.357 | 0.250 | 0.332 | 0.538 | 0.132 |
| P4 | L | 0.292 | 0.379 | 0.688 | 0.746 | 0.412 | 0.504 | 0.541 | 0.851 | 0.646 | 0.602 | 0.775 |
| P4 | R | 0.896 | 0.068 | 0.283 | 0.634 | 0.171 | 0.454 | 0.533 | 0.649 | 0.737 | 0.549 | 0.564 |
| P5 | L | 0.435 | 0.271 | 0.869 | 0.026 | 0.424 | 0.338 | 0.427 | 0.207 | 0.581 | 0.641 | 0.286 |
| P5 | R | 0.418 | 0.632 | 0.661 | 0.169 | 0.228 | 0.213 | 0.208 | 0.208 | 0.416 | 0.206 | 0.257 |
| P6 | L | 0.396 | 0.655 | 0.229 | 0.644 | 0.237 | 0.273 | 0.016 | 0.161 | 0.031 | 0.335 | 0.235 |
| P6 | R | 0.629 | -0.033 | 0.402 | 0.484 | 0.424 | 0.310 | 0.056 | 0.104 | 0.300 | 0.433 | 0.639 |

| P | H | VMLOI | VMTOP | VMTOA | VMLOS | VMIPSP | VMIPSA | VMPCSS | VMPCSI | VMFS | VMFI | VMFA |
|---|---|---|---|---|---|---|---|---|---|---|---|---|
| P1 | L | 88 | 214 | 72 | 170 | 251 | 239 | 301 | 226 | 261 | 312 | 250 |
| P1 | R | 95 | 157 | 101 | 86 | 192 | 294 | 274 | 94 | 270 | 197 | 163 |
| P2 | L | 95 | 144 | 39 | 53 | 214 | 357 | 328 | 103 | 291 | 89 | 98 |
| P2 | R | 96 | 123 | 52 | 78 | 143 | 334 | 333 | 101 | 185 | 60 | 30 |
| P3 | L | 160 | 152 | 98 | 176 | 142 | 246 | 163 | 121 | 131 | 64 | 108 |
| P3 | R | 143 | 196 | 126 | 123 | 251 | 410 | 243 | 164 | 193 | 213 | 185 |
| P4 | L | 22 | 33 | 56 | 95 | 202 | 184 | 66 | 48 | 55 | 20 | 61 |
| P4 | R | 34 | 76 | 62 | 66 | 165 | 221 | 138 | 77 | 82 | 82 | 55 |
| P5 | L | 34 | 26 | 12 | 46 | 103 | 113 | 124 | 61 | 53 | 38 | 87 |
| P5 | R | 69 | 30 | 71 | 25 | 91 | 190 | 147 | 86 | 193 | 113 | 102 |
| P6 | L | 24 | 73 | 85 | 75 | 61 | 82 | 85 | 34 | 41 | 79 | 120 |
| P6 | R | 93 | 46 | 53 | 81 | 66 | 174 | 62 | 69 | 79 | 108 | 50 |

**Table S6, Descriptive statistics for correlations between VSTM load preferences estimated from the constant orientation change condition and the cortical surface distance across the map, related to Figure 3A.** Top: *p*-values of correlations in each map for each hemisphere, after correction for false discovery rate and upsampling between the native fMRI scan resolution and the participant's cortical surface model. Middle: Pearson's correlation coefficient (*r*) in each map. Bottom: Number of selected measurement in each map after correction for upsampling. Selected measurements must have >20% variance explained.

| P | H | VMLOI | VMTOP | VMTOA | VMLOS | VMIPSP | VMIPSA | VMPCSS | VMPCSI | VMFS | VMFI | VMFA | p<0.05 | % |
|---|---|---|---|---|---|---|---|---|---|---|---|---|---|---|
| P1 | L | 0.00023 | 4.8e-05 | 6.6e-10 | 1.6e-07 | 1.8e-17 | 1.2e-28 | 5.8e-51 | 2.6e-29 | 2e-43 | 5.1e-15 | 5.5e-08 | 11 | 100 |
| P1 | R | 6.5e-05 | 2.6e-07 | 0.087 | 2.8e-11 | 0.044 | 4.9e-26 | 2.2e-22 | 0.00038 | 9.8e-38 | 0.00096 | 6.2e-25 | 10 | 91 |
| P2 | L | 4.2e-05 | 3.6e-12 | 7.4e-06 | 9.8e-05 | 2.9e-28 | 1.5e-30 | 6e-31 | 9.1e-19 | 5e-24 | 1.1e-08 | 4.4e-18 | 11 | 100 |
| P2 | R | 3.3e-14 | 1.9e-16 | 3.4e-08 | 1.3e-13 | 3.1e-14 | 2e-12 | 6.1e-09 | 1e-11 | 1.1e-11 | 1e-06 | 1.3e-05 | 11 | 100 |
| P3 | L | 2.9e-10 | 2.1e-10 | 2.6e-10 | 2.1e-10 | 3.5e-08 | 2.2e-12 | 1e-13 | 8e-13 | 7.5e-12 | 7.2e-09 | 6.5e-05 | 11 | 100 |
| P3 | R | 5.5e-17 | 0.077 | 0.00017 | 4.4e-05 | 4.5e-33 | 1.3e-06 | 3.2e-10 | 6.1e-09 | 3.1e-06 | 3.1e-21 | 4.2e-05 | 10 | 91 |
| P4 | L | 0.00059 | 0.0071 | 4.4e-05 | 7.7e-12 | 5.5e-09 | 3.3e-12 | 1.5e-08 | 2.1e-10 | 1.3e-09 | 0.028 | 5.4e-05 | 11 | 100 |
| P4 | R | 1.1e-12 | 0.0022 | 2.2e-06 | 4.2e-08 | 0.054 | 8.1e-05 | 1.2e-06 | 2.3e-05 | 1.6e-08 | 4.4e-11 | 0.076 | 9 | 82 |
| P5 | L | 0.16 | 0.21 | 0.32 | 0.13 | 6.2e-08 | 0.019 | 3.4e-06 | 1 | 0.0098 | 0.00026 | 0.021 | 6 | 55 |
| P5 | R | 0.0016 | 1.2e-05 | 3.5e-06 | 0.17 | 0.014 | 0.0016 | 3.6e-08 | 0.3 | 0.053 | 0.039 | 0.087 | 7 | 64 |
| P6 | L | 0.94 | 2.1e-06 | 0.032 | 0.0016 | 0.099 | 4.2e-05 | 0.9 | 0.58 | 1 | 0.13 | 0.15 | 4 | 36 |
| P6 | R | 1.6e-05 | 0.11 | 0.0011 | 0.00079 | 0.00047 | 8.4e-06 | 0.83 | 1 | 0.043 | 0.00064 | 1.6e-07 | 8 | 73 |
| <0.05 | | 10 | 9 | 10 | 10 | 10 | 12 | 10 | 8 | 10 | 11 | 9 | 109 | Tot |
| % | | 83 | 75 | 83 | 83 | 83 | 100 | 83 | 67 | 83 | 92 | 75 | 83 | % |

| P | H | VMLOI | VMTOP | VMTOA | VMLOS | VMIPSP | VMIPSA | VMPCSS | VMPCSI | VMFS | VMFI | VMFA |
|---|---|---|---|---|---|---|---|---|---|---|---|---|
| P1 | L | 0.409 | 0.287 | 0.686 | 0.435 | 0.518 | 0.647 | 0.746 | 0.670 | 0.728 | 0.450 | 0.343 |
| P1 | R | 0.532 | 0.438 | 0.208 | 0.708 | 0.158 | 0.580 | 0.583 | 0.386 | 0.687 | 0.246 | 0.707 |
| P2 | L | 0.458 | 0.581 | 0.665 | 0.566 | 0.668 | 0.561 | 0.586 | 0.694 | 0.547 | 0.577 | 0.746 |
| P2 | R | 0.723 | 0.711 | 0.713 | 0.730 | 0.612 | 0.378 | 0.330 | 0.641 | 0.490 | 0.628 | 0.750 |
| P3 | L | 0.495 | 0.511 | 0.622 | 0.513 | 0.509 | 0.454 | 0.570 | 0.624 | 0.580 | 0.710 | 0.389 |
| P3 | R | 0.688 | 0.169 | 0.358 | 0.409 | 0.682 | 0.262 | 0.421 | 0.467 | 0.361 | 0.640 | 0.347 |
| P4 | L | 0.613 | 0.481 | 0.477 | 0.606 | 0.423 | 0.508 | 0.625 | 0.791 | 0.724 | 0.567 | 0.529 |
| P4 | R | 0.883 | 0.373 | 0.548 | 0.612 | 0.170 | 0.286 | 0.491 | 0.570 | 0.658 | 0.699 | 0.231 |
| P5 | L | 0.272 | 0.424 | 0.508 | 0.280 | 0.621 | 0.242 | 0.470 | -0.102 | 0.479 | 0.743 | 0.368 |
| P5 | R | 0.449 | 0.675 | 0.628 | 0.312 | 0.297 | 0.249 | 0.477 | 0.136 | 0.206 | 0.226 | 0.214 |
| P6 | L | 0.021 | 0.564 | 0.256 | 0.353 | 0.219 | 0.441 | 0.024 | 0.145 | -0.017 | 0.203 | 0.151 |
| P6 | R | 0.532 | 0.246 | 0.426 | 0.400 | 0.397 | 0.325 | 0.032 | -0.064 | 0.259 | 0.368 | 0.747 |

| P | H | VMLOI | VMTOP | VMTOA | VMLOS | VMIPSP | VMIPSA | VMPCSS | VMPCSI | VMFS | VMFI | VMFA |
|---|---|---|---|---|---|---|---|---|---|---|---|---|
| P1 | L | 81 | 206 | 66 | 141 | 247 | 240 | 292 | 224 | 266 | 289 | 252 |
| P1 | R | 53 | 134 | 74 | 70 | 176 | 287 | 242 | 85 | 272 | 187 | 163 |
| P2 | L | 78 | 127 | 39 | 44 | 217 | 368 | 336 | 128 | 304 | 88 | 100 |
| P2 | R | 85 | 104 | 48 | 79 | 133 | 344 | 313 | 95 | 180 | 53 | 27 |
| P3 | L | 152 | 144 | 89 | 143 | 110 | 229 | 152 | 113 | 124 | 53 | 105 |
| P3 | R | 118 | 120 | 111 | 99 | 243 | 352 | 217 | 148 | 168 | 181 | 142 |
| P4 | L | 29 | 32 | 71 | 111 | 186 | 175 | 71 | 46 | 55 | 16 | 55 |
| P4 | R | 38 | 69 | 69 | 70 | 140 | 194 | 93 | 51 | 62 | 71 | 65 |
| P5 | L | 30 | 11 | 6 | 32 | 66 | 101 | 94 | 38 | 30 | 20 | 42 |
| P5 | R | 49 | 36 | 48 | 22 | 73 | 166 | 127 | 65 | 96 | 91 | 70 |
| P6 | L | 28 | 65 | 76 | 82 | 62 | 85 | 52 | 19 | 42 | 60 | 98 |
| P6 | R | 62 | 46 | 59 | 71 | 78 | 191 | 67 | 23 | 66 | 87 | 38 |

**Table S7, Descriptive statistics for correlations between VSTM load preferences estimated from the odd numbered scans and the cortical surface distance across the map, related to Figure 3A.** Top: *p*-values of correlations in each map for each hemisphere, after correction for false discovery rate and upsampling between the native fMRI scan resolution and the participant's cortical surface model. Middle: Pearson's correlation coefficient (*r*) in each map. Bottom: Number of selected measurement in each map after correction for upsampling. Selected measurements must have >20% variance explained.

| P | H | VMLOI | VMTOP | VMTOA | VMLOS | VMIPSP | VMIPSA | VMPCSS | VMPCSI | VMFS | VMFI | VMFA | p<0.05 | % |
|---|---|---|---|---|---|---|---|---|---|---|---|---|---|---|
| P1 | L | 3.4e-10 | 2.3e-05 | 4.3e-07 | 3e-20 | 1.3e-21 | 2e-20 | 3.6e-77 | 1.7e-27 | 1.8e-56 | 1.4e-11 | 3.7e-07 | 11 | 100 |
| P1 | R | 2.4e-05 | 1.9e-09 | 8.5e-05 | 8.7e-10 | 0.33 | 4e-20 | 4.8e-29 | 0.00017 | 1.1e-37 | 8.7e-10 | 6.4e-24 | 10 | 91 |
| P2 | L | 0.013 | 7.4e-17 | 2e-06 | 0.0042 | 1.9e-25 | 1.1e-43 | 1.1e-31 | 8.9e-09 | 5.1e-11 | 7.1e-05 | 1.5e-14 | 11 | 100 |
| P2 | R | 3.2e-20 | 1.9e-12 | 1.5e-07 | 2.7e-09 | 8.3e-10 | 4e-11 | 1.5e-11 | 0.0045 | 6.5e-13 | 9e-07 | 0.22 | 10 | 91 |
| P3 | L | 2.2e-05 | 1e-09 | 1 | 0.0015 | 1e-07 | 1.1e-06 | 0.00015 | 7.3e-09 | 1.2e-06 | 4.6e-08 | 0.056 | 9 | 82 |
| P3 | R | 0.00016 | 8e-08 | 5.1e-11 | 0.0098 | 3.5e-27 | 0.0016 | 4.4e-06 | 1 | 8.6e-05 | 1.9e-14 | 4e-05 | 10 | 91 |
| P4 | L | 0.00018 | 8.9e-05 | 0.0048 | 2.7e-15 | 4.6e-11 | 9.9e-10 | 0.0042 | 1.6e-05 | 6.8e-07 | 5.8e-06 | 6.4e-08 | 11 | 100 |
| P4 | R | 4.1e-15 | 0.13 | 1 | 2.4e-08 | 2.5e-06 | 6.3e-07 | 8.4e-14 | 0.39 | 0.0097 | 4.8e-06 | 0.0014 | 8 | 73 |
| P5 | L | 1.3e-05 | 0.12 | 3.7e-05 | 0.041 | 1.4e-11 | 0.00027 | 1.2e-06 | 0.18 | 3.1e-06 | 7.8e-06 | 0.0026 | 9 | 82 |
| P5 | R | 0.00082 | 0.0017 | 3.5e-09 | 0.39 | 0.03 | 1.7e-06 | 0.016 | 0.082 | 8.6e-07 | 0.063 | 0.00034 | 8 | 73 |
| P6 | L | 0.16 | 0.00023 | 0.21 | 1.3e-08 | 0.00014 | 0.1 | 0.74 | 0.015 | 0.014 | 1.6e-07 | 2.8e-14 | 7 | 64 |
| P6 | R | 9.9e-14 | 0.97 | 0.0066 | 0.00062 | 7.5e-08 | 0.00079 | 1 | 0.0099 | 1.9e-05 | 0.25 | 6e-10 | 8 | 73 |
| <0.05 | | 11 | 9 | 9 | 11 | 11 | 11 | 10 | 8 | 12 | 10 | 10 | 112 | Tot |
| % | | 92 | 75 | 75 | 92 | 92 | 92 | 83 | 67 | 100 | 83 | 83 | 85 | % |

| P | H | VMLOI | VMTOP | VMTOA | VMLOS | VMIPSP | VMIPSA | VMPCSS | VMPCSI | VMFS | VMFI | VMFA |
|---|---|---|---|---|---|---|---|---|---|---|---|---|
| P1 | L | 0.635 | 0.298 | 0.623 | 0.660 | 0.580 | 0.573 | 0.838 | 0.647 | 0.795 | 0.398 | 0.319 |
| P1 | R | 0.463 | 0.501 | 0.385 | 0.676 | 0.076 | 0.532 | 0.643 | 0.409 | 0.695 | 0.430 | 0.691 |
| P2 | L | 0.253 | 0.607 | 0.668 | 0.388 | 0.644 | 0.626 | 0.588 | 0.493 | 0.374 | 0.387 | 0.668 |
| P2 | R | 0.797 | 0.617 | 0.584 | 0.589 | 0.489 | 0.345 | 0.343 | 0.246 | 0.499 | 0.540 | 0.183 |
| P3 | L | 0.349 | 0.545 | -0.038 | 0.269 | 0.479 | 0.332 | 0.330 | 0.561 | 0.501 | 0.689 | 0.223 |
| P3 | R | 0.362 | 0.472 | 0.613 | 0.297 | 0.657 | 0.175 | 0.310 | -0.016 | 0.327 | 0.594 | 0.350 |
| P4 | L | 0.664 | 0.739 | 0.569 | 0.772 | 0.437 | 0.470 | 0.388 | 0.716 | 0.628 | 0.699 | 0.651 |
| P4 | R | 0.878 | 0.214 | -0.152 | 0.709 | 0.406 | 0.386 | 0.642 | 0.145 | 0.353 | 0.488 | 0.470 |
| P5 | L | 0.574 | 0.324 | 0.776 | 0.261 | 0.581 | 0.330 | 0.419 | 0.166 | 0.519 | 0.559 | 0.275 |
| P5 | R | 0.439 | 0.487 | 0.685 | 0.182 | 0.242 | 0.345 | 0.211 | 0.196 | 0.328 | 0.172 | 0.308 |
| P6 | L | 0.220 | 0.404 | 0.124 | 0.651 | 0.452 | 0.167 | 0.032 | 0.452 | 0.355 | 0.494 | 0.529 |
| P6 | R | 0.714 | 0.011 | 0.369 | 0.389 | 0.617 | 0.241 | -0.172 | 0.317 | 0.337 | 0.096 | 0.567 |

| P | H | VMLOI | VMTOP | VMTOA | VMLOS | VMIPSP | VMIPSA | VMPCSS | VMPCSI | VMFS | VMFI | VMFA |
|---|---|---|---|---|---|---|---|---|---|---|---|---|
| P1 | L | 84 | 206 | 58 | 159 | 237 | 230 | 296 | 231 | 262 | 285 | 259 |
| P1 | R | 81 | 135 | 105 | 68 | 181 | 269 | 249 | 84 | 263 | 199 | 167 |
| P2 | L | 101 | 163 | 43 | 56 | 215 | 404 | 341 | 129 | 307 | 106 | 109 |
| P2 | R | 91 | 113 | 73 | 91 | 150 | 369 | 392 | 140 | 195 | 77 | 51 |
| P3 | L | 149 | 115 | 82 | 145 | 118 | 219 | 135 | 97 | 89 | 52 | 79 |
| P3 | R | 110 | 124 | 100 | 79 | 219 | 342 | 224 | 129 | 147 | 146 | 139 |
| P4 | L | 28 | 23 | 24 | 76 | 221 | 162 | 56 | 30 | 55 | 35 | 59 |
| P4 | R | 47 | 54 | 48 | 50 | 133 | 167 | 114 | 41 | 56 | 84 | 46 |
| P5 | L | 53 | 26 | 22 | 66 | 121 | 124 | 133 | 73 | 76 | 59 | 125 |
| P5 | R | 58 | 41 | 61 | 26 | 86 | 194 | 138 | 85 | 229 | 126 | 139 |
| P6 | L | 46 | 83 | 113 | 65 | 70 | 103 | 135 | 30 | 50 | 107 | 190 |
| P6 | R | 85 | 50 | 56 | 78 | 67 | 202 | 59 | 69 | 164 | 158 | 108 |

**Table S8, Descriptive statistics for correlations between VSTM load preferences estimated from the even numbered scans and the cortical surface distance across the map, related to Figure 3A.** Top: *p*-values of correlations in each map for each hemisphere, after correction for false discovery rate and upsampling between the native fMRI scan resolution and the participant's cortical surface model. Middle: Pearson's correlation coefficient (*r*) in each map. Bottom: Number of selected measurement in each map after correction for upsampling. Selected measurements must have >20% variance explained.

| P | H | VMLOI | VMTOP | VMTOA | VMLOS | VMIPSP | VMIPSA | VMPCSS | VMPCSI | VMFS | VMFI | VMFA | p<0.05 | % |
|---|---|---|---|---|---|---|---|---|---|---|---|---|---|---|
| P1 | L | 0.00026 | 5.7e-05 | 1.2e-16 | 1.8e-07 | 1.9e-17 | 1.2e-28 | 4.1e-56 | 7.7e-22 | 1.4e-43 | 5.1e-15 | 6e-08 | 11 | 100 |
| P1 | R | 7.6e-05 | 2.8e-07 | 0.094 | 2.7e-11 | 2.1e-05 | 4.7e-24 | 1.9e-30 | 0.00015 | 5.7e-42 | 0.0063 | 6.2e-25 | 10 | 91 |
| P2 | L | 5e-05 | 3.3e-12 | 2.1e-07 | 0.00012 | 2.9e-28 | 1.5e-30 | 1.7e-27 | 9.1e-19 | 4.9e-44 | 1.1e-08 | 1.3e-12 | 11 | 100 |
| P2 | R | 3.3e-14 | 2e-16 | 3.7e-08 | 1.3e-13 | 3.1e-14 | 1.8e-12 | 2.1e-05 | 1e-11 | 1e-11 | 6.4e-05 | 0.0015 | 11 | 100 |
| P3 | L | 3e-10 | 2.1e-10 | 0.0061 | 2e-10 | 3.8e-08 | 2e-12 | 1e-13 | 8e-13 | 0.00023 | 7.7e-09 | 0.046 | 11 | 100 |
| P3 | R | 9.1e-19 | 0.084 | 0.0002 | 5.3e-05 | 3.6e-33 | 1.5e-06 | 3.2e-10 | 6.4e-09 | 3.5e-06 | 2.3e-16 | 5e-05 | 10 | 91 |
| P4 | L | 0.00068 | 0.0079 | 5.3e-05 | 7.5e-12 | 5.9e-09 | 3.1e-12 | 6.4e-09 | 0.094 | 8e-13 | 0.03 | 6.3e-05 | 10 | 91 |
| P4 | R | 1e-12 | 0.0025 | 1 | 4.6e-08 | 0.059 | 0.00068 | 1 | 2.7e-05 | 1.7e-08 | 4.2e-11 | 0.083 | 7 | 64 |
| P5 | L | 0.18 | 0.23 | 0.36 | 1 | 6.7e-08 | 0.021 | 0.0066 | 1 | 0.011 | 1 | 0.023 | 5 | 45 |
| P5 | R | 0.0019 | 1.4e-05 | 0.012 | 0.19 | 0.00091 | 0.0019 | 3.9e-08 | 0.83 | NoData | 0.043 | 0.094 | 7 | 64 |
| P6 | L | 1 | 1 | 1 | 0.017 | 0.11 | NoData | 5.3e-05 | 1 | 0.63 | 1 | 0.3 | 2 | 18 |
| P6 | R | 0.0055 | 0.0005 | 0.51 | 0.083 | 0.00091 | 0.96 | 1 | 0.18 | 0.58 | 1 | 0.0094 | 4 | 36 |
| <0.05 | | 10 | 9 | 7 | 9 | 10 | 10 | 10 | 7 | 9 | 9 | 9 | 99 | Tot |
| % | | 83 | 75 | 58 | 75 | 83 | 83 | 83 | 58 | 75 | 75 | 75 | 75 | % |

| P | H | VMLOI | VMTOP | VMTOA | VMLOS | VMIPSP | VMIPSA | VMPCSS | VMPCSI | VMFS | VMFI | VMFA |
|---|---|---|---|---|---|---|---|---|---|---|---|---|
| P1 | L | 0.409 | 0.287 | 0.771 | 0.435 | 0.518 | 0.647 | 0.747 | 0.545 | 0.728 | 0.450 | 0.343 |
| P1 | R | 0.532 | 0.438 | 0.208 | 0.708 | 0.325 | 0.576 | 0.651 | 0.453 | 0.689 | 0.206 | 0.707 |
| P2 | L | 0.458 | 0.581 | 0.737 | 0.566 | 0.668 | 0.561 | 0.558 | 0.694 | 0.738 | 0.577 | 0.676 |
| P2 | R | 0.723 | 0.711 | 0.713 | 0.730 | 0.612 | 0.378 | 0.324 | 0.641 | 0.490 | 0.516 | 0.601 |
| P3 | L | 0.495 | 0.511 | 0.279 | 0.513 | 0.509 | 0.454 | 0.570 | 0.624 | 0.368 | 0.710 | 0.188 |
| P3 | R | 0.694 | 0.169 | 0.358 | 0.409 | 0.682 | 0.262 | 0.421 | 0.467 | 0.361 | 0.652 | 0.347 |
| P4 | L | 0.613 | 0.481 | 0.477 | 0.606 | 0.423 | 0.508 | 0.657 | 0.315 | 0.821 | 0.567 | 0.529 |
| P4 | R | 0.883 | 0.373 | -0.518 | 0.612 | 0.170 | 0.257 | -0.020 | 0.570 | 0.658 | 0.699 | 0.231 |
| P5 | L | 0.272 | 0.424 | 0.508 | -0.438 | 0.621 | 0.242 | 0.304 | 0.022 | 0.479 | -0.006 | 0.368 |
| P5 | R | 0.449 | 0.675 | 0.382 | 0.312 | 0.394 | 0.249 | 0.477 | 0.036 | NoData | 0.226 | 0.214 |
| P6 | L | -0.052 | -0.092 | 0.000 | 0.293 | 0.219 | NoData | 0.398 | -0.229 | 0.088 | 0.017 | 0.121 |
| P6 | R | 0.435 | 0.435 | 0.111 | 0.295 | 0.400 | 0.022 | -0.130 | 0.209 | 0.117 | -0.119 | 0.429 |

| P | H | VMLOI | VMTOP | VMTOA | VMLOS | VMIPSP | VMIPSA | VMPCSS | VMPCSI | VMFS | VMFI | VMFA |
|---|---|---|---|---|---|---|---|---|---|---|---|---|
| P1 | L | 81 | 206 | 83 | 141 | 247 | 240 | 319 | 277 | 266 | 289 | 252 |
| P1 | R | 53 | 134 | 74 | 70 | 178 | 268 | 252 | 70 | 300 | 192 | 163 |
| P2 | L | 78 | 127 | 39 | 44 | 217 | 368 | 333 | 128 | 259 | 88 | 90 |
| P2 | R | 85 | 104 | 48 | 79 | 133 | 344 | 179 | 95 | 180 | 58 | 27 |
| P3 | L | 152 | 144 | 105 | 143 | 110 | 229 | 152 | 113 | 103 | 53 | 127 |
| P3 | R | 128 | 120 | 111 | 99 | 243 | 352 | 217 | 148 | 168 | 131 | 142 |
| P4 | L | 29 | 32 | 71 | 111 | 186 | 175 | 66 | 33 | 51 | 16 | 55 |
| P4 | R | 38 | 69 | 49 | 70 | 140 | 186 | 129 | 51 | 62 | 71 | 65 |
| P5 | L | 30 | 11 | 6 | 23 | 66 | 101 | 86 | 35 | 30 | 16 | 42 |
| P5 | R | 49 | 36 | 46 | 22 | 73 | 166 | 127 | 90 | NoData | 91 | 70 |
| P6 | L | 21 | 33 | 19 | 72 | 62 | NoData | 105 | 12 | 47 | 25 | 89 |
| P6 | R | 43 | 65 | 50 | 40 | 71 | 68 | 135 | 49 | 35 | 96 | 39 |

**Table S9, Descriptive statistics for correlations between VSTM load preferences estimated from the odd numbered scans and the cortical surface distance across maps defined from the complementary data from even numbered scans.** Top: *p*-values of correlations in each map for each hemisphere, after correction for false discovery rate and upsampling between the native fMRI scan resolution and the participant's cortical surface model. Middle: Pearson's correlation coefficient (*r*) in each map. Bottom: Number of selected measurement in each map after correction for upsampling. Selected measurements must have >20% variance explained.

| P | H | VMLOI | VMTOP | VMTOA | VMLOS | VMIPSP | VMIPSA | VMPCSS | VMPCSI | VMFS | VMFI | VMFA | p<0.05 | % |
|---|---|---|---|---|---|---|---|---|---|---|---|---|---|---|
| P1 | L | 3.7e-05 | 2.8e-05 | 5.6e-07 | 4.2e-14 | 1.3e-21 | 2e-20 | 1.2e-79 | 7.5e-12 | 1.8e-56 | 1.4e-11 | 4e-06 | 11 | 100 |
| P1 | R | 1.2e-05 | 2.5e-09 | 9.8e-05 | 1.1e-09 | 0.035 | 1.3e-39 | 3.5e-28 | 0.063 | 9.3e-36 | 9.8e-05 | 0.01 | 10 | 91 |
| P2 | L | 0.015 | 8.4e-17 | 1e-06 | 0.0047 | 1.9e-25 | 1.1e-43 | 9e-32 | 9.1e-06 | 8.3e-14 | 8.5e-05 | 5.6e-15 | 11 | 100 |
| P2 | R | 3.4e-20 | 2e-12 | 0.001 | 3.5e-09 | 1e-09 | 4.3e-11 | 1e-22 | 5.3e-16 | 6.8e-13 | 1.2e-06 | 0.22 | 10 | 91 |
| P3 | L | 2.7e-05 | 1.3e-09 | 1 | 0.0018 | 1.3e-07 | 1.5e-06 | 0.00017 | 9.5e-09 | 1.5e-06 | 0.034 | 0.06 | 9 | 82 |
| P3 | R | 0.00097 | 1.1e-07 | 1.2e-07 | 0.011 | 3.5e-27 | 0.0019 | 5.6e-06 | 1 | 0.0001 | 2.1e-14 | 4.9e-05 | 10 | 91 |
| P4 | L | 1e-05 | 0.0047 | 0.0055 | 2.8e-15 | 0.054 | 7.8e-05 | 0.0047 | 1.9e-05 | 1.5e-06 | 7.3e-06 | 8.6e-08 | 10 | 91 |
| P4 | R | 2.6e-13 | 0.068 | 1 | 3.1e-08 | 0.019 | 8e-07 | 2.4e-13 | 1 | 0.011 | 6.1e-06 | 1 | 7 | 64 |
| P5 | L | 0.044 | 0.089 | 0.86 | 0.052 | 0.06 | 0.00033 | 1.3e-05 | 0.0039 | 0.0029 | 0.19 | 9e-05 | 6 | 55 |
| P5 | R | 0.0067 | 3.3e-07 | 1.5e-06 | 0.73 | 6e-07 | 2.1e-06 | 0.018 | 0.072 | 0.00053 | 1 | 0.00033 | 8 | 73 |
| P6 | L | 0.33 | 1 | 1 | 1.7e-08 | 0.00016 | NoData | 0.0026 | 0.099 | 0.71 | 0.0033 | 0.0092 | 5 | 45 |
| P6 | R | 0.18 | 1 | 0.026 | 0.94 | 0.00074 | 9.9e-08 | 0.00094 | 0.052 | 0.041 | 0.033 | 1 | 6 | 55 |
| <0.05 | | 10 | 8 | 8 | 9 | 10 | 11 | 12 | 6 | 11 | 10 | 8 | 103 | Tot |
| % | | 83 | 67 | 67 | 75 | 83 | 92 | 100 | 50 | 92 | 83 | 67 | 78 | % |

| P | H | VMLOI | VMTOP | VMTOA | VMLOS | VMIPSP | VMIPSA | VMPCSS | VMPCSI | VMFS | VMFI | VMFA |
|---|---|---|---|---|---|---|---|---|---|---|---|---|
| P1 | L | 0.502 | 0.298 | 0.623 | 0.568 | 0.580 | 0.573 | 0.816 | 0.413 | 0.795 | 0.398 | 0.308 |
| P1 | R | 0.436 | 0.501 | 0.385 | 0.676 | 0.162 | 0.680 | 0.635 | 0.217 | 0.662 | 0.262 | 0.253 |
| P2 | L | 0.253 | 0.607 | 0.688 | 0.388 | 0.644 | 0.626 | 0.588 | 0.394 | 0.424 | 0.387 | 0.678 |
| P2 | R | 0.797 | 0.617 | 0.391 | 0.589 | 0.489 | 0.345 | 0.578 | 0.664 | 0.499 | 0.540 | 0.194 |
| P3 | L | 0.349 | 0.545 | -0.038 | 0.269 | 0.479 | 0.332 | 0.330 | 0.561 | 0.501 | 0.279 | 0.223 |
| P3 | R | 0.310 | 0.472 | 0.547 | 0.297 | 0.657 | 0.175 | 0.310 | -0.016 | 0.327 | 0.594 | 0.350 |
| P4 | L | 0.669 | 0.627 | 0.569 | 0.772 | 0.155 | 0.317 | 0.388 | 0.716 | 0.630 | 0.699 | 0.651 |
| P4 | R | 0.894 | 0.234 | -0.091 | 0.709 | 0.218 | 0.386 | 0.705 | 0.008 | 0.353 | 0.488 | -0.170 |
| P5 | L | 0.344 | 0.406 | 0.047 | 0.285 | 0.203 | 0.330 | 0.381 | 0.459 | 0.395 | 0.189 | 0.405 |
| P5 | R | 0.303 | 0.586 | 0.626 | 0.073 | 0.433 | 0.345 | 0.211 | 0.176 | 0.328 | -0.072 | 0.416 |
| P6 | L | 0.197 | -0.023 | -0.060 | 0.651 | 0.452 | NoData | 0.262 | 0.315 | 0.091 | 0.430 | 0.258 |
| P6 | R | 0.225 | -0.121 | 0.330 | 0.020 | 0.389 | 0.617 | 0.241 | 0.272 | 0.302 | 0.238 | -0.111 |

| P | H | VMLOI | VMTOP | VMTOA | VMLOS | VMIPSP | VMIPSA | VMPCSS | VMPCSI | VMFS | VMFI | VMFA |
|---|---|---|---|---|---|---|---|---|---|---|---|---|
| P1 | L | 66 | 206 | 58 | 159 | 237 | 230 | 337 | 270 | 262 | 285 | 234 |
| P1 | R | 101 | 135 | 105 | 68 | 186 | 293 | 249 | 81 | 284 | 233 | 112 |
| P2 | L | 101 | 163 | 43 | 56 | 215 | 404 | 341 | 129 | 303 | 106 | 108 |
| P2 | R | 91 | 113 | 73 | 91 | 150 | 369 | 252 | 123 | 195 | 77 | 47 |
| P3 | L | 149 | 115 | 82 | 145 | 118 | 219 | 135 | 97 | 89 | 64 | 79 |
| P3 | R | 120 | 124 | 88 | 79 | 219 | 342 | 224 | 129 | 147 | 146 | 139 |
| P4 | L | 38 | 20 | 24 | 76 | 170 | 162 | 56 | 30 | 52 | 35 | 59 |
| P4 | R | 38 | 67 | 30 | 50 | 126 | 167 | 86 | 39 | 56 | 84 | 36 |
| P5 | L | 38 | 20 | 38 | 52 | 95 | 124 | 133 | 41 | 60 | 56 | 95 |
| P5 | R | 86 | 70 | 53 | 40 | 133 | 194 | 138 | 114 | 117 | 114 | 76 |
| P6 | L | 31 | 60 | 43 | 65 | 70 | NoData | 143 | 31 | 30 | 49 | 111 |
| P6 | R | 41 | 47 | 50 | 79 | 78 | 67 | 202 | 57 | 51 | 89 | 63 |

**Table S10, Descriptive statistics for correlations between VSTM load preferences estimated from the even numbered scans and the cortical surface distance across maps defined from the complementary data from odd numbered scans.** Top: *p*-values of correlations in each map for each hemisphere, after correction for false discovery rate and upsampling between the native fMRI scan resolution and the participant's cortical surface model. Middle: Pearson's correlation coefficient (*r*) in each map. Bottom: Number of selected measurement in each map after correction for upsampling. Selected measurements must have >20% variance explained.

| P | H | VMLOI | VMTOP | VMTOA | VMLOS | VMIPSP | VMIPSA | VMPCSS | VMPCSI | VMFS | VMFI | VMFA | p<0.05 | % |
|---|---|---|---|---|---|---|---|---|---|---|---|---|---|---|
| P1 | L | 0.14 | 1.8e-19 | 0.033 | 0.017 | 3.5e-16 | 1.8e-20 | 6.1e-42 | 1.9e-09 | 2.8e-33 | 9.1e-15 | 8e-07 | 10 | 91 |
| P1 | R | 0.017 | 0.046 | 0.083 | 0.085 | 4.2e-11 | 1.1e-17 | 8.2e-10 | 0.016 | 1.7e-34 | 5.4e-10 | 8e-14 | 9 | 82 |
| P2 | L | 0.062 | 3e-09 | 0.02 | 0.016 | 1.4e-17 | 2.6e-30 | 1.3e-12 | 4.9e-07 | 1.6e-19 | 2.5e-08 | 2.3e-06 | 10 | 91 |
| P2 | R | 0.012 | 1 | 0.056 | 0.99 | 0.0091 | 5.5e-10 | 0.0092 | 0.0071 | 0.00038 | 0.23 | 0.43 | 6 | 55 |
| P3 | L | 1 | 0.47 | 1 | 0.33 | 0.017 | 0.0001 | 0.0097 | 0.41 | 0.18 | 0.38 | 0.33 | 3 | 27 |
| P3 | R | 0.85 | 0.99 | 1 | 0.77 | 0.014 | 0.14 | 0.0082 | 7e-06 | 0.0092 | 0.61 | 0.61 | 4 | 36 |
| P4 | L | 0.33 | 1 | 0.47 | 0.77 | 0.61 | 0.27 | 0.92 | 0.65 | 0.19 | 1 | 1 | 0 | 0 |
| P4 | R | 0.72 | 1 | 1 | 0.61 | 0.27 | 0.1 | 0.046 | 0.071 | 0.85 | 0.081 | 1 | 1 | 9 |
| P5 | L | 0.77 | 0.84 | 0.98 | 0.99 | 1 | 1 | 1 | 0.77 | 0.77 | 1 | 1 | 0 | 0 |
| P5 | R | 0.14 | 0.85 | 0.78 | 1 | 0.33 | 0.98 | 0.58 | 0.39 | 0.64 | 0.61 | 0.62 | 0 | 0 |
| P6 | L | 0.34 | 0.09 | 0.82 | 0.042 | 0.096 | 1 | 1 | 1 | 1 | 0.033 | 0.028 | 3 | 27 |
| P6 | R | 0.62 | 1 | 0.00015 | 0.016 | 0.13 | 5e-07 | 0.058 | 1 | 0.069 | 0.13 | 0.016 | 4 | 36 |
| <0.05 | | 2 | 3 | 3 | 4 | 6 | 6 | 7 | 5 | 5 | 4 | 5 | 50 | Tot |
| % | | 17 | 25 | 25 | 33 | 50 | 50 | 58 | 42 | 42 | 33 | 42 | 38 | % |

| P | H | VMLOI | VMTOP | VMTOA | VMLOS | VMIPSP | VMIPSA | VMPCSS | VMPCSI | VMFS | VMFI | VMFA |
|---|---|---|---|---|---|---|---|---|---|---|---|---|
| P1 | L | 0.275 | 0.651 | 0.377 | 0.271 | 0.572 | 0.608 | 0.732 | 0.432 | 0.719 | 0.489 | 0.380 |
| P1 | R | 0.453 | 0.242 | 0.272 | 0.289 | 0.533 | 0.551 | 0.438 | 0.351 | 0.737 | 0.489 | 0.619 |
| P2 | L | 0.283 | 0.556 | 0.528 | 0.472 | 0.589 | 0.579 | 0.412 | 0.487 | 0.544 | 0.614 | 0.552 |
| P2 | R | 0.361 | -0.112 | 0.382 | 0.028 | 0.277 | 0.362 | 0.178 | 0.364 | 0.319 | 0.251 | 0.289 |
| P3 | L | -0.034 | 0.120 | -0.115 | 0.144 | 0.316 | 0.320 | 0.283 | 0.141 | 0.212 | 0.200 | 0.198 |
| P3 | R | 0.054 | 0.030 | -0.011 | 0.087 | 0.228 | 0.108 | 0.256 | 0.440 | 0.287 | 0.081 | 0.102 |
| P4 | L | 0.409 | 0.058 | 0.394 | 0.075 | 0.073 | 0.124 | 0.056 | 0.341 | 0.284 | -0.204 | -0.005 |
| P4 | R | 0.194 | -0.031 | -0.078 | 0.142 | 0.147 | 0.175 | 0.322 | 0.464 | 0.081 | 0.317 | -0.065 |
| P5 | L | 0.184 | 0.284 | 0.319 | 0.063 | 0.011 | -0.100 | 0.013 | 0.123 | 0.159 | -0.059 | -0.184 |
| P5 | R | 0.335 | 0.132 | 0.107 | -0.058 | 0.174 | 0.024 | 0.096 | 0.162 | 0.101 | 0.121 | 0.140 |
| P6 | L | 0.557 | 0.517 | 0.125 | 0.391 | 0.333 | -0.009 | 0.011 | -0.081 | 0.052 | 0.579 | 0.396 |
| P6 | R | 0.216 | 0.085 | 0.745 | 0.464 | 0.307 | 0.478 | 0.506 | 0.062 | 0.430 | 0.243 | 0.558 |

| P | H | VMLOI | VMTOP | VMTOA | VMLOS | VMIPSP | VMIPSA | VMPCSS | VMPCSI | VMFS | VMFI | VMFA |
|---|---|---|---|---|---|---|---|---|---|---|---|---|
| P1 | L | 44 | 161 | 44 | 104 | 183 | 203 | 255 | 196 | 211 | 240 | 180 |
| P1 | R | 36 | 97 | 60 | 52 | 146 | 220 | 198 | 63 | 205 | 159 | 128 |
| P2 | L | 63 | 108 | 25 | 34 | 186 | 340 | 297 | 109 | 253 | 76 | 73 |
| P2 | R | 65 | 71 | 36 | 68 | 121 | 306 | 294 | 73 | 147 | 40 | 17 |
| P3 | L | 91 | 81 | 41 | 85 | 77 | 171 | 112 | 69 | 65 | 39 | 48 |
| P3 | R | 64 | 58 | 71 | 46 | 159 | 282 | 145 | 111 | 111 | 108 | 75 |
| P4 | L | 12 | 8 | 9 | 60 | 137 | 139 | 41 | 7 | 35 | 7 | 34 |
| P4 | R | 14 | 25 | 34 | 37 | 99 | 130 | 54 | 22 | 30 | 45 | 13 |
| P5 | L | 12 | 5 | 3 | 16 | 29 | 80 | 50 | 24 | 15 | 10 | 20 |
| P5 | R | 31 | 14 | 29 | 7 | 60 | 124 | 90 | 56 | 60 | 52 | 35 |
| P6 | L | 7 | 16 | 18 | 38 | 37 | 61 | 17 | 5 | 9 | 18 | 42 |
| P6 | R | 16 | 8 | 24 | 36 | 38 | 113 | 20 | 9 | 26 | 59 | 24 |

**Table S11, Descriptive statistics for correlations between tuning widths estimated from odd and even numbered scan runs.** Top: *p*-values of correlations in each map for each hemisphere, after correction for false discovery rate and upsampling between the native fMRI scan resolution and the participant's cortical surface model. Middle: Pearson's correlation coefficient (*r*) in each map. Bottom: Number of selected measurement in each map after correction for upsampling. Selected measurements must have >20% variance explained by the response model in both sets of scanning runs.

| P | H | VMLOI | VMTOP | VMTOA | VMLOS | VMIPSP | VMIPSA | VMPCSS | VMPCSI | VMFS | VMFI | VMFA | p<0.05 | % |
|---|---|---|---|---|---|---|---|---|---|---|---|---|---|---|
| P1 | L | 0.038 | 0.00031 | 0.19 | 5.2e-05 | 6.4e-13 | 1.9e-14 | 2.5e-25 | 5.2e-05 | 7.6e-36 | 1.9e-16 | 0.00016 | 10 | 91 |
| P1 | R | 0.19 | 0.32 | 0.25 | 0.29 | 0.00055 | 2.5e-08 | 0.35 | 0.32 | 1e-15 | 0.021 | 6.7e-10 | 5 | 45 |
| P2 | L | 0.049 | 1.5e-11 | 0.00032 | 0.0032 | 1.7e-21 | 8.5e-33 | 1.4e-19 | 1.1e-07 | 2e-24 | 0.00028 | 0.00032 | 11 | 100 |
| P2 | R | 1.4e-09 | 1 | 0.056 | 0.052 | 7.4e-05 | 1.2e-13 | 4.1e-06 | 0.15 | 7.3e-06 | 0.26 | 0.09 | 5 | 45 |
| P3 | L | 0.79 | 0.84 | 1 | 1 | 1 | 1 | 0.72 | 0.75 | 1 | 0.11 | 1 | 0 | 0 |
| P3 | R | 0.58 | 1 | 1 | 1 | 1 | 1 | 0.13 | 0.034 | 0.19 | 1 | 0.84 | 1 | 9 |
| P4 | L | 0.15 | 1 | 0.035 | 1 | 1 | 0.33 | 0.58 | 0.55 | 0.4 | 1 | 0.33 | 1 | 9 |
| P4 | R | 0.32 | 0.25 | 1 | 0.56 | 1 | 0.73 | 1 | 0.081 | 0.97 | 0.043 | 0.29 | 1 | 9 |
| P5 | L | 0.19 | 1 | 0.97 | 0.49 | 1 | 0.56 | 0.63 | 1 | 0.48 | 0.6 | 1 | 0 | 0 |
| P5 | R | 0.23 | 0.73 | 0.79 | 1 | 0.33 | 1.4e-05 | 0.015 | 0.36 | 0.86 | 0.081 | 0.72 | 2 | 18 |
| P6 | L | 0.25 | 0.26 | 0.92 | 0.13 | 0.25 | 0.075 | 0.39 | 0.58 | 0.46 | 1 | 0.017 | 1 | 9 |
| P6 | R | 0.97 | 0.25 | 7e-05 | 0.024 | 0.25 | 4.1e-06 | 0.25 | 1 | 0.036 | 0.13 | 0.0029 | 5 | 45 |
| <0.05 | | 3 | 2 | 3 | 3 | 4 | 6 | 4 | 3 | 5 | 4 | 5 | 42 | Tot |
| % | | 25 | 17 | 25 | 25 | 33 | 50 | 33 | 25 | 42 | 33 | 42 | 32 | % |

| P | H | VMLOI | VMTOP | VMTOA | VMLOS | VMIPSP | VMIPSA | VMPCSS | VMPCSI | VMFS | VMFI | VMFA |
|---|---|---|---|---|---|---|---|---|---|---|---|---|
| P1 | L | 0.366 | 0.310 | 0.239 | 0.428 | 0.509 | 0.519 | 0.614 | 0.312 | 0.750 | 0.509 | 0.307 |
| P1 | R | 0.264 | 0.155 | 0.218 | 0.218 | 0.321 | 0.391 | 0.093 | 0.182 | 0.540 | 0.220 | 0.537 |
| P2 | L | 0.306 | 0.625 | 0.691 | 0.535 | 0.647 | 0.601 | 0.511 | 0.538 | 0.593 | 0.449 | 0.454 |
| P2 | R | 0.694 | 0.005 | 0.370 | 0.296 | 0.380 | 0.423 | 0.281 | 0.207 | 0.394 | 0.248 | 0.468 |
| P3 | L | 0.059 | 0.057 | -0.023 | -0.060 | -0.096 | 0.014 | 0.069 | 0.075 | -0.058 | 0.372 | -0.015 |
| P3 | R | 0.101 | -0.145 | -0.168 | -0.043 | -0.001 | 0.001 | 0.170 | 0.255 | 0.175 | -0.018 | 0.054 |
| P4 | L | 0.560 | 0.046 | 0.798 | -0.137 | 0.011 | 0.115 | 0.138 | 0.851 | 0.202 | 0.060 | 0.233 |
| P4 | R | 0.347 | 0.347 | 0.036 | 0.159 | -0.018 | 0.057 | -0.011 | 0.446 | 0.059 | 0.367 | 0.377 |
| P5 | L | 0.519 | 0.055 | 0.166 | 0.295 | 0.021 | 0.117 | 0.116 | 0.031 | 0.299 | 0.262 | 0.004 |
| P5 | R | 0.322 | 0.297 | 0.104 | -0.106 | 0.174 | 0.417 | 0.320 | 0.170 | 0.069 | 0.316 | 0.110 |
| P6 | L | 0.517 | 0.395 | 0.089 | 0.309 | 0.282 | 0.291 | 0.229 | 0.243 | 0.403 | -0.081 | 0.403 |
| P6 | R | 0.071 | 0.521 | 0.756 | 0.457 | 0.259 | 0.468 | 0.359 | -0.036 | 0.466 | 0.255 | 0.644 |

| P | H | VMLOI | VMTOP | VMTOA | VMLOS | VMIPSP | VMIPSA | VMPCSS | VMPCSI | VMFS | VMFI | VMFA |
|---|---|---|---|---|---|---|---|---|---|---|---|---|
| P1 | L | 47 | 161 | 53 | 100 | 191 | 206 | 247 | 196 | 203 | 248 | 179 |
| P1 | R | 44 | 79 | 51 | 44 | 138 | 215 | 191 | 56 | 205 | 158 | 128 |
| P2 | L | 62 | 104 | 27 | 36 | 182 | 338 | 294 | 97 | 259 | 75 | 71 |
| P2 | R | 65 | 68 | 40 | 65 | 124 | 308 | 308 | 82 | 144 | 38 | 21 |
| P3 | L | 91 | 75 | 42 | 85 | 78 | 161 | 94 | 69 | 61 | 31 | 47 |
| P3 | R | 76 | 86 | 75 | 36 | 158 | 261 | 135 | 102 | 98 | 101 | 82 |
| P4 | L | 12 | 9 | 9 | 61 | 133 | 134 | 41 | 3 | 35 | 12 | 33 |
| P4 | R | 17 | 22 | 32 | 35 | 98 | 126 | 52 | 24 | 29 | 45 | 16 |
| P5 | L | 12 | 7 | 6 | 14 | 41 | 62 | 47 | 30 | 14 | 12 | 21 |
| P5 | R | 27 | 7 | 29 | 7 | 59 | 120 | 80 | 55 | 44 | 48 | 40 |
| P6 | L | 10 | 16 | 20 | 41 | 31 | 59 | 29 | 15 | 9 | 24 | 48 |
| P6 | R | 19 | 10 | 25 | 34 | 37 | 104 | 20 | 23 | 29 | 60 | 24 |

**Table S12, Descriptive statistics for correlations between tuning widths estimated from constant performance and constant orientation conditions.** Top: *p*-values of correlations in each map for each hemisphere, after correction for false discovery rate and upsampling between the native fMRI scan resolution and the participant's cortical surface model. Middle: Pearson's correlation coefficient (*r*) in each map. Bottom: Number of selected measurement in each map after correction for upsampling. Selected measurements must have >20% variance explained by the response model in both conditions.

| P | H | VMLOI | VMTOP | VMTOA | VMLOS | VMIPSP | VMIPSA | VMPCSS | VMPCSI | VMFS | VMFI | VMFA | p<0.05 | % |
|---|---|---|---|---|---|---|---|---|---|---|---|---|---|---|
| P1 | L | 7.3e-05 | 1.9e-49 | 6.4e-17 | 1e-11 | 3e-72 | 3.1e-75 | 2.5e-120 | 8.9e-78 | 5.8e-97 | 8.8e-95 | 4.6e-49 | 11 | 100 |
| P1 | R | 0.0032 | 4.1e-38 | 1.4e-28 | 4.1e-19 | 1.9e-39 | 4.4e-66 | 7.5e-64 | 1.6e-20 | 1.3e-99 | 6e-36 | 3.4e-73 | 11 | 100 |
| P3 | L | NoData | 1.4e-13 | 1 | 1.6e-20 | 1e-20 | 2.2e-13 | 8.9e-20 | 2.4e-14 | 2.8e-08 | 1 | 1 | 7 | 64 |
| P3 | R | 0.91 | 0.17 | 0.0035 | 0.087 | 2.9e-35 | 9.7e-11 | 4.4e-12 | 3.1e-12 | 2.2e-18 | 7.4e-24 | NoData | 7 | 64 |
| P4 | L | NoData | NoData | NoData | 5.8e-14 | 8.3e-13 | 2.9e-12 | 0.00014 | NoData | 0.0012 | 1 | 0.0029 | 6 | 55 |
| P4 | R | NoData | NoData | 1 | 3.9e-11 | 0.0002 | 2.8e-08 | 1e-08 | 2.4e-08 | 0.0017 | 7.6e-06 | NoData | 7 | 64 |
| P5 | L | NoData | NoData | NoData | 1 | 1 | 5.1e-06 | 0.0013 | 1 | 0.077 | 0.021 | 3.9e-05 | 4 | 36 |
| P5 | R | 8.7e-08 | 1 | 1 | NoData | 0.00023 | 1.9e-19 | 3.4e-05 | 2e-05 | 7.5e-10 | 5.2e-08 | NoData | 7 | 64 |
| P6 | L | 1 | 0.13 | 0.049 | 0.0015 | 0.7 | 0.006 | 1.4e-06 | NoData | 0.74 | 1 | 0.0011 | 5 | 45 |
| P6 | R | 0.53 | 0.63 | 2.2e-06 | 3.5e-05 | 7.5e-07 | 7e-10 | 0.00026 | 0.95 | 0.54 | 1.4e-16 | 0.00067 | 7 | 64 |
| <0.05 | | 3 | 3 | 5 | 7 | 8 | 10 | 10 | 6 | 7 | 7 | 6 | 72 | Tot |
| % | | 30 | 30 | 50 | 70 | 80 | 100 | 100 | 60 | 70 | 70 | 60 | 65 | % |

| P | H | VMLOI | VMTOP | VMTOA | VMLOS | VMIPSP | VMIPSA | VMPCSS | VMPCSI | VMFS | VMFI | VMFA |
|---|---|---|---|---|---|---|---|---|---|---|---|---|
| P1 | L | 0.446 | 0.850 | 0.876 | 0.702 | 0.828 | 0.913 | 0.871 | 0.923 | 0.955 | 0.667 | 0.833 |
| P1 | R | 0.505 | 0.932 | 0.825 | 0.999 | 0.579 | 0.770 | 0.470 | 0.622 | 0.925 | 0.549 | 0.893 |
| P3 | L | NoData | 0.501 | -0.167 | 0.736 | 0.717 | 0.861 | 0.661 | 0.979 | 0.473 | -0.020 | -0.556 |
| P3 | R | 0.122 | 0.694 | 0.642 | 0.991 | 0.888 | 0.816 | 0.478 | 0.803 | 0.871 | 0.786 | NoData |
| P4 | L | NoData | NoData | NoData | 0.729 | 0.789 | 0.765 | 0.879 | NoData | 0.489 | 0.247 | 0.573 |
| P4 | R | NoData | NoData | -0.078 | 0.928 | 0.672 | 0.685 | 0.727 | 0.938 | 0.784 | 0.682 | NoData |
| P5 | L | NoData | NoData | NoData | -0.518 | -0.213 | 0.380 | 0.096 | -0.017 | 0.542 | 0.256 | 0.140 |
| P5 | R | 0.056 | -0.056 | -0.178 | NoData | 0.009 | 0.362 | 0.273 | 0.227 | 0.320 | 0.185 | NoData |
| P6 | L | 0.055 | 0.112 | 0.069 | 0.463 | 0.104 | 0.330 | 0.328 | NoData | 0.514 | -0.108 | 0.015 |
| P6 | R | 0.957 | 0.523 | 0.880 | 0.412 | 0.410 | 0.351 | 0.261 | 0.905 | 0.591 | 0.752 | 0.165 |

| P | H | VMLOI | VMTOP | VMTOA | VMLOS | VMIPSP | VMIPSA | VMPCSS | VMPCSI | VMFS | VMFI | VMFA |
|---|---|---|---|---|---|---|---|---|---|---|---|---|
| P1 | L | 12 | 50 | 24 | 12 | 133 | 68 | 161 | 113 | 107 | 155 | 134 |
| P1 | R | 2 | 27 | 24 | 2 | 113 | 150 | 146 | 43 | 101 | 140 | 73 |
| P3 | L | NoData | 9 | 3 | 5 | 12 | 53 | 25 | 5 | 1 | 13 | 1 |
| P3 | R | 11 | 2 | 1 | 1 | 39 | 74 | 36 | 17 | 5 | 43 | NoData |
| P4 | L | NoData | NoData | NoData | 29 | 40 | 88 | 13 | NoData | 17 | 2 | 32 |
| P4 | R | NoData | NoData | 3 | 16 | 53 | 57 | 33 | 4 | 7 | 32 | NoData |
| P5 | L | NoData | NoData | NoData | 4 | 10 | 46 | 16 | 10 | 1 | 5 | 4 |
| P5 | R | 11 | 3 | 6 | NoData | 20 | 71 | 20 | 60 | 13 | 20 | NoData |
| P6 | L | 3 | 12 | 15 | 42 | 25 | 46 | 4 | NoData | 3 | 12 | 23 |
| P6 | R | 5 | 2 | 12 | 31 | 19 | 79 | 13 | 1 | 3 | 46 | 5 |

**Table S13, Descriptive statistics for correlations between VSTM load preferences estimated from the main experiment and the unpredictable order control.** Top: *p*-values of correlations in each map for each hemisphere, after correction for false discovery rate and upsampling between the native fMRI scan resolution and the participant's cortical surface model. Middle: Pearson's correlation coefficient (*r*) in each map. Bottom: Number of selected measurement in each map after correction for upsampling. Selected measurements must have >10% variance explained by the response model in both experiments.

**Video S1, VSTM task in one cycle of constant performance condition of the main experiment, at the orientation difference thresholds of an example participant, related to Figure 1A-B,.** Each trial started with an array of six colored bars with one to six different orientations. Participants were asked to briefly remember this six-item display. This display was followed by a backwards mask to avoid afterimages, a blank screen, then a probe bar with the same color as one of the memorized bars, then another blank screen. Participants responded whether the orientation of the probe bar was the same or different from the memorized bar of that color.

**Video S2, VSTM task in one cycle of the reverse display role control experiment, at the same orientation difference thresholds used in Movie S1.** In each trial, participants were asked to briefly remember the one-item display. The array of six colored bars should be compared to the bar in the previous, remembered (rather than the next, upcoming) one-item display. Participants responded whether the orientation of the probe bar was the same or different from the bar of that color in the next six-item display.

**Video S3, VSTM task in one cycle of the constant performance condition in the unpredictable trial order control experiment, at the orientation difference thresholds of the example participant used in Movie S1.** The structure of each trial was the same as in the main experiment: participants responded whether the orientation of the probe bar was the same or different from the bar of that color in the previous six-item display. However, the VSTM load on each trial was pseudo-random and unpredictable to the participant.
